# Age-related microbiome metabolites alter RNA splicing and chromatin accessibility in the brain

**DOI:** 10.1101/2025.10.03.680371

**Authors:** Meenakshi Chakraborty, Sophia M. Shi, Imani E. Porter, Daniel J. Richard, Georgi K. Marinov, Mukundh N. Murthy, Ashley A. Moore, Jenna L. E. Blum, Aravind Natarajan, James W. Jahng, Joseph C. Wu, Sydney X. Lu, Shawn M. Davidson, William J. Greenleaf, Nay L. Saw, Mehrdad Shamloo, Anne Brunet, Tony Wyss-Coray, Ami S. Bhatt

## Abstract

The gut microbiome generates diverse metabolites that can enter the bloodstream and alter host biology, including brain function. Hundreds of physiologically relevant, gut-brain signaling molecules likely exist; however, there has been no systematic, high-throughput effort to identify and validate them. Here, we integrate computational, *in vitro*, and *in vivo* approaches to pinpoint microbiome-derived metabolites whose blood levels change during aging, and that induce molecular changes in the mouse brain. First, we mine large-scale metabolomics datasets from human cohorts (each *n ≥* 1200) to identify 30 microbiome-associated metabolites whose blood levels change with age. We then screen this panel in an *in vitro* transcriptomic assay to identify metabolites that perturb genes linked to age-related neurodegeneration. To assess *in vivo* relevance, we then test four metabolites in male mice by acute exposure, using multi-omic approaches to evaluate the metabolites’ impact on cellular functions in the brain. With RNA-seq, we confirm known effects of trimethylamine N-oxide (TMAO), including changes in mitochondrial pathways, and further discover its effects on the pathways of glycolysis, GABAergic signaling, and RNA splicing. Additionally, using both RNA- and ATAC-seq, we identify glycodeoxycholate (GDCA), a microbiome-derived secondary bile acid, as a potent regulator of chromatin accessibility and of genes involved in protecting the brain from age-related stressors. GDCA also acutely reduces locomotion in male but not female mice. In summary, we present a generalizable framework for identifying microbiome metabolites that impact host biology, and apply it to identify age-related microbial metabolites that affect processes related to brain aging and neurodegeneration.

## Introduction

Aging is the greatest risk factor for many neurodegenerative diseases. Remarkably, age-related conditions that often precede these diseases, such as mild cognitive impairment (MCI), can be reversible^1,2^. Thus, identifying potentially modifiable age-related factors that influence brain function could alter and perhaps improve the natural history of disease initiation and progression.

One physiological compartment that changes with aging is the gut microbiome^3–5^, likely due to factors such as increased permeability of the gut lining, increased medication usage, immunosenescence, and accumulated exposure to environmental toxins. In light of growing interest in the microbiota-gut-brain axis^6,7^, a natural question arises: how do age-related microbiome changes affect the brain? Studies involving fecal microbiota transfer between young and aged mice suggest that the aged microbiome can have both beneficial and detrimental effects on the brain, depending on the experimental context and the specific outcome examined, such as neuroinflammation, neurogenesis, or behavior^8–12^. However, the microbiome-related molecules and underlying molecular mechanisms that drive these phenotypes remain largely uncharacterized in both animals and humans, which limits translation of this knowledge to the clinic. This gap in the gut-brain field is partly due to the underutilization of *in vitro* systems, which can help create a pipeline from hypothesis-generating data analyses to *in vivo* validation.

The microbiome is known to substantially impact the levels of blood metabolites, with hundreds of metabolites differing in abundance between germ-free mice and their conventionally raised counterparts^13,14^. This is significant because blood metabolites enable the proper functioning of host organs; when their levels change, this can have profound effects on physiology^15–17^. We therefore hypothesized that age-related microbiome changes affect the brain by altering the levels of bioactive blood metabolites. These metabolites could impact the blood-brain barrier - a key regulator of brain homeostasis whose functionality declines with age^18^ - and perhaps even cross it to act directly on neurons and glia. Importantly, age-related microbial metabolites would not necessarily be expected to be detrimental to brain function and may even be beneficial, given previous work suggesting that some age-associated microbiome changes may protect against inflammaging and frailty^19–21^.

To better understand how microbial metabolites might affect barrier and brain function, we leveraged public datasets^14,22–28^ to identify a panel of 30 blood metabolites whose levels: 1) change with human aging, and 2) are significantly associated with the presence and/or composition of the gut microbiome. We then screened these candidates *in vitro* for their ability to alter the expression of genes in pathways related to age-related neurodegeneration, and validated four of the most interesting metabolites *in vivo* through multi-omic analyses of the male mouse brain. Our transcriptomics results confirm the known effects of trimethylamine N-oxide (TMAO) on the brain - TMAO being one of a small number of microbiome-derived metabolites with established links to brain function^7^ - and identify novel effects, including impacts on RNA splicing. In addition, we find that glycodeoxycholate (GDCA), a microbiome-derived bile acid, alters chromatin accessibility in the brain and silences many genes that maintain brain homeostasis after stress and injury. Finally, in behavioral experiments, we find that GDCA, whose blood levels are correlated with neurodegeneration and cognitive impairment in multiple human cohorts^29–37^, acutely reduces locomotion in male but not female mice. Together, our findings on TMAO and GDCA, our curated panel of age- and microbiome-associated blood metabolites, and the integrative framework we present - combining computational, *in vitro*, and *in vivo* approaches - provide a foundation for substantial advances in identifying microbiome-derived molecules that play key roles in host physiology.

## Results

### Identifying microbiome-associated metabolites whose blood levels change with age

There have been extensive efforts to characterize the specific metabolomic changes that occur with aging^22^. In addition, recent work has identified metabolites associated with microbiome composition and activity^14,26–28^. We leveraged these data to build a panel of metabolites for medium-throughput *in vitro* testing (Fig. 1; Supplementary Table 1; Methods).

**Fig. 1.**
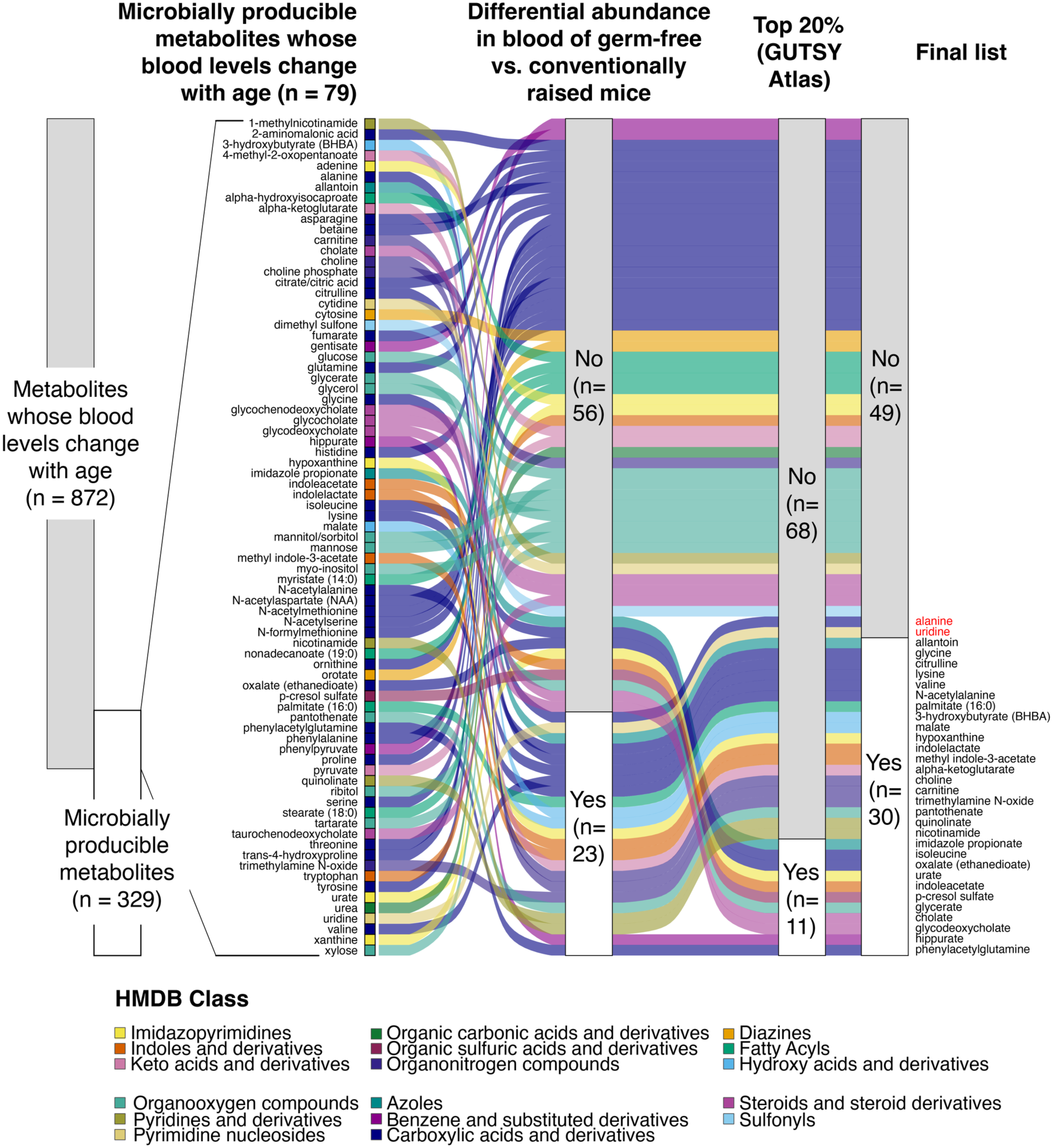
Workflow used to define a panel of 30 microbiome-associated metabolites whose blood levels change with age. First, metabolites whose blood levels changed with age in large cohort(s) were overlapped with a curated list of metabolites that are producible by microbes, either alone or via host-microbe cometabolism. Metabolites were retained for the final panel if they met at least one of two criteria: **(1)** showed significantly different blood levels in germ-free vs. conventionally raised mice (based on Lai *et al.*, 2021^14^), and/or **(2)** ranked in the top 20% for the proportion of variance in blood levels “explained” by the gut microbiome (metric from the GUTSY Atlas^27^). After excluding alanine and uridine (see Results), the final panel comprised 30 metabolites.

Specifically, we first referenced a review on the metabolomics of aging^22^ to identify three large cohort studies^23–25^ that each analyzed blood samples from at least 1,200 participants. Pooling data across these studies yielded hundreds of metabolites whose blood levels changed with age in at least one study. Separately, we curated a list of human-associated metabolites that have been reported to be produced by microbes, either alone or via host-microbe “cometabolism” (where both host and microbial metabolism contribute to the final product). Intersecting this curated list with the age-associated metabolites yielded 79 unique overlapping metabolites.

For experimental feasibility, we further narrowed the list, prioritizing metabolites most strongly linked to the gut microbiome. We used two key resources: **(1)** a study comparing blood metabolite levels between germ-free and conventionally raised mice^14^; and **(2)** the GUTSY Atlas^27^, which reported the proportion of variance in blood metabolite levels “explained” by the gut microbiome for 1,168 metabolites based on paired stool-blood data from over 8,000 humans. Metabolites were retained for the final panel if they met at least one of two criteria: **(1)** showed significantly different blood levels between germ-free and conventionally raised mice^14^; **(2)** ranked in the top 20% of blood metabolites for the GUTSY variance-explained metric.

We eliminated L-alanine because it is expected to have minimal impact on cellular processes^38^, and uridine because its blood levels increased with age in one study^23^ but decreased with age in another^25^. The final panel comprised 30 metabolites, 26 of which increased with age and 4 of which decreased with age (Supplementary Table 1). The metabolites encompassed a broad spectrum of categories, spanning six super classes, thirteen classes, and sixteen sub classes within the Human Metabolome Database (HMDB)^39^ taxonomy (Fig. S1). These metabolites include compounds that have been associated with detrimental effects on host health, as well as beneficial or compensatory effects, depending on context^40–44^.

### *In vitro* screening reveals that microbiome-associated metabolites perturb genes and pathways linked to age-related neurodegeneration

While *in vivo* validation remains the gold standard for assessing physiological relevance, we began with *in vitro* screening, as testing dozens of metabolites *in vivo* for potential effects on the brain would not have been practically feasible. We chose human brain endothelial cells as our model system because they line the brain’s blood vessels and are the first brain cell type encountered by blood metabolites. In fact, many blood metabolites never reach other brain cell types, since endothelial cells are a key component of the blood-brain barrier (BBB), which selectively regulates entry into the brain parenchyma. Importantly, even metabolites that do not cross the BBB could significantly impact brain function, since the BBB is itself a key regulator of brain homeostasis^18^.

We chose to use hCMEC/D3 cells, an immortalized human brain endothelial cell line widely used in BBB studies^45^. To conduct the screen, each metabolite was added individually to low-passage hCMEC/D3 cells in triplicate, alongside matched vehicle controls containing the corresponding solvent in the media (Fig. 2A; Supplementary Table 3). After a 3-hour incubation (chosen over longer timepoints to minimize metabolite clearance), cells were lysed in TRIzol for downstream RNA extraction and sequencing. Each metabolite was added at a representative physiological concentration, primarily based on its listed blood concentrations in HMDB (Methods; Supplementary Table 1; Supplementary Note 1). In addition to vehicle controls, we included lipopolysaccharide (LPS) controls in each sequencing batch, given the known effects of LPS on human brain endothelial cells^46^. As expected, across sequencing batches, LPS induced the expression of genes in KEGG pathways associated with inflammation, such as “TNF signaling pathway” and “NOD-like receptor signaling pathway” (Fig. 2B; Fig. S2; Supplementary Table 2).

**Fig. 2.**
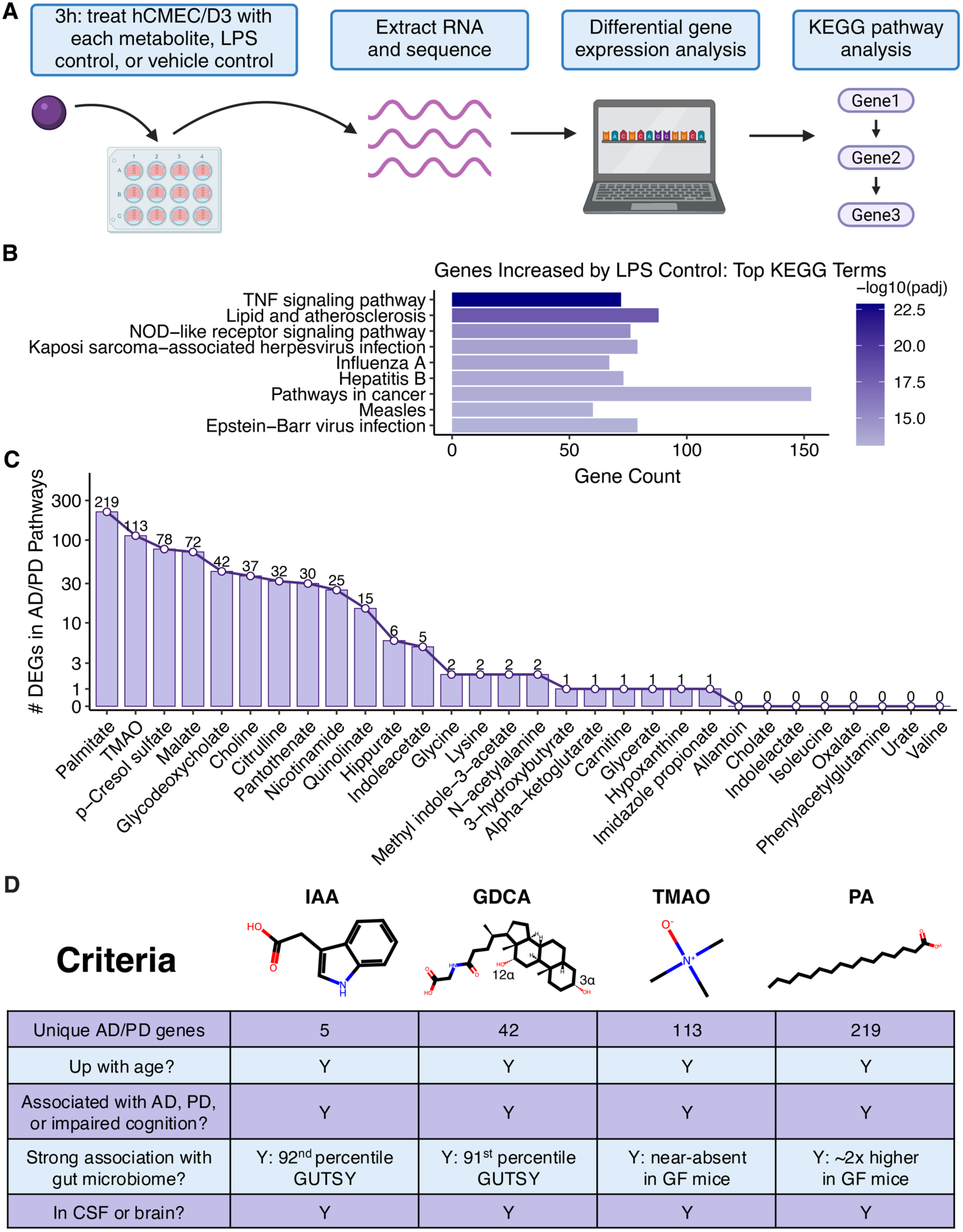
*In vitro* screening reveals that microbiome-associated metabolites perturb genes and pathways linked to age-associated neurodegenerative disease. **(A)** Schematic illustrating the setup of the *in vitro* transcriptomic screen. The full panel of 30 metabolites was distributed across four sequencing batches, with each metabolite included in one batch. Within each batch, at least *n* = 3 wells were treated with each assigned metabolite, its respective vehicle control, or the LPS control (see Methods and Supplementary Table 3). **(B)** Top enriched KEGG pathways for the genes significantly upregulated by the LPS control in the first sequencing batch. The analogous results for batches #2-4 are presented in Fig. S2. The non-specific “KEGG root term” was filtered out. **(C)** Number of differentially expressed genes (DEGs) per metabolite in the KEGG pathways for Alzheimer’s Disease (AD) and/or Parkinson’s Disease (PD). **(D)** Identity and characteristics of the four metabolites chosen for subsequent *in vivo* validation of their effects on the brain. IAA = indoleacetic acid, GDCA = glycodeoxycholate, TMAO = trimethylamine N-oxide, PA = palmitic acid, AD = Alzheimer’s Disease, PD = Parkinson’s Disease, GF = germ-free, CSF = cerebrospinal fluid. GUTSY refers to the GUTSY Atlas^27^, which reports microbiome-attributable variance in blood levels for 1,168 blood metabolites. Structure diagrams were produced with rdkit^61^. Relevant studies (e.g., those reporting metabolite detection in CSF or brain) are cited in the text.

The number of differentially expressed genes (DEGs), relative to vehicle controls, varied considerably across the 30 test metabolites, reflecting substantial differences in the magnitude of their transcriptional impact (Fig. S3; Supplementary Tables 4-5). The median number of DEGs was 45.5, with an interquartile range (IQR) of 18.5 to 359.5, and a total range of 3 to 6,419 genes affected. Twelve metabolites perturbed ≥5 genes in the KEGG pathways for “Alzheimer’s Disease” (AD) and/or “Parkinson’s Disease” (PD), which together contain several hundred genes and represent the two most common age-related neurodegenerative disorders (Fig. 2C; Supplementary Table 6). The five-gene threshold separated metabolites perturbing several AD/PD pathway genes from those perturbing few or none (Fig. 2C). The affected AD/PD genes converged on processes including energy metabolism, proteasome-mediated protein degradation, and inflammatory signaling, whose disruptions are central to neurodegenerative disease^47^.

We prioritized four metabolites for *in vivo* validation of their effects on brain function: indoleacetic acid (IAA), glycodeoxycholate (GDCA), trimethylamine N-oxide (TMAO), and palmitic acid (PA). Several previous studies have investigated TMAO’s effects on the brain^48^, so we included it partially as a control and also to expand the scope of knowledge about its impact on the brain. All four chosen metabolites share the following characteristics (Fig. 2D). They **(1)** impact the expression of at least 5 unique genes in the AD and/or PD KEGG pathways in our transcriptomic screen (Fig. 2C), **(2)** have been reported to increase in the blood with age (Supplementary Table 1), **(3)** have been previously associated with AD, PD, and/or impaired cognition in humans^29–37,48–53^, **(4)** are strongly associated with production or regulation by the gut microbiome^14,27,54,55^, and **(5)** have been detected in human cerebrospinal fluid and/or brain^56–60^.

### TMAO disrupts transcriptional homeostasis and RNA splicing in the mouse brain

To interrogate the effects of each of the four chosen metabolites on cellular functions in the brain *in vivo*, we conducted bulk transcriptomics on mouse brain hemispheres following intravenous retro-orbital injection of each metabolite or vehicle (2-hour treatment; *n =* 4 adult male mice per injection group; Fig. 3A). Doses were chosen to reflect physiologically relevant exposures, by converting blood concentrations observed in humans to mouse-equivalent doses using well-established scaling guidelines (Methods). The experiment design was informed by prior work on TMAO^62^, such that TMAO could serve as a positive control. The main modification was the use of intravenous retro-orbital injection - a well-established technique^18,63–65^ - in place of intraperitoneal injection, to enable direct entry into systemic circulation and more precise control of blood metabolite levels.

**Fig. 3.**
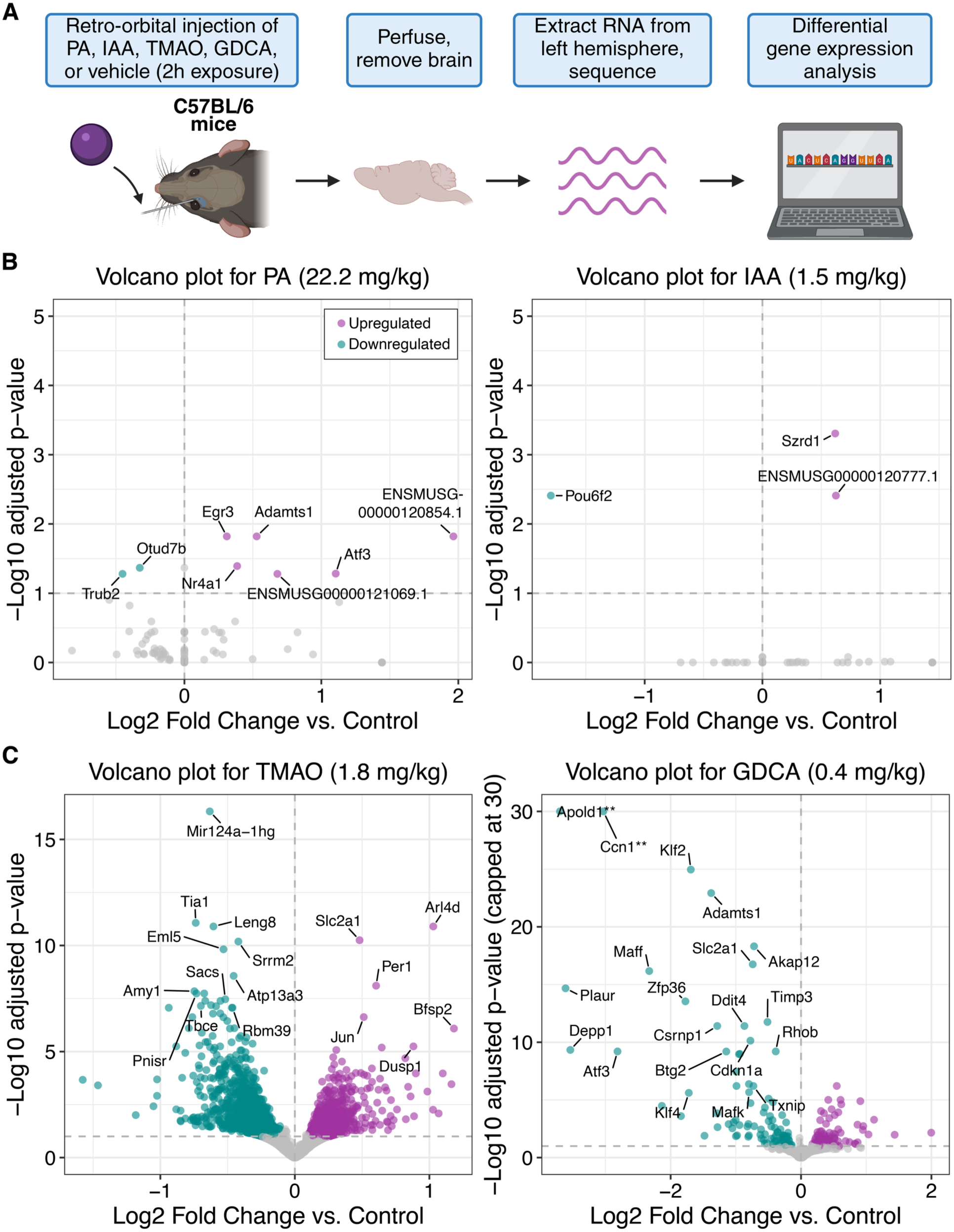
The four metabolites selected for *in vivo* experiments impact brain gene expression to different extents. **(A)** Schematic illustrating the setup of the *in vivo* transcriptomics experiment; *n =* 4 C57BL/6 male mice were injected with each metabolite or vehicle. **(B)** Volcano plots showingthe results of differential gene expression analysis between PA- and IAA-treated brains and their vehicle controls. **(C)** Volcano plots showing the results of differential gene expression analysis between TMAO- and GDCA-treated brains and their vehicle controls. In both **(B)** and **(C)**, the y-axis represents the −log10(adjusted *p*-value) from DESeq2, and the x-axis represents the log2FoldChange vs. vehicle control, after applying the recommended log2FoldChange shrinkage via lfcShrink^69^. Purple dots represent significantly upregulated genes, with an adjusted *p* ≤ 0.1 (DESeq2 default) and shrunken log2FoldChange ≥ 0.1. Teal dots represent significantly downregulated genes (adjusted *p* ≤ 0.1 and shrunken log2FoldChange ≤ −0.1). The GDCA volcano plot y-axis was capped at 30 to prevent distortion by extreme values. *Apold1* and *Ccn1* are marked with double asterisks because their significance exceeded this limit; their uncapped adjusted p-values were 1.82E-108 and 3.50E-70, respectively (Supplementary Table 7).

For TMAO, the chosen mouse dose was 1.8 mg/kg, the same as the original study^62^, approximating 30μM concentration in human blood. The remaining doses were 22.2 mg/kg for PA (∼100μM human), 1.5 mg/kg for IAA (∼10μM human), and 0.4 mg/kg for GDCA (∼1μM human). All compounds were non-cytotoxic in hCMEC/D3 cells at concentrations equal to or above their selected human-equivalent blood concentrations (Fig. S7). For PA - unlike for IAA and GDCA - the human-equivalent concentration used for the mouse experiment was lower than the concentration used in the *in vitro* transcriptomic screen. This was because the original concentration of 500μM, while in the physiologically relevant range^66,67^, was cytotoxic to hCMEC/D3 cells (Fig. S7), likely explaining the disproportionately large number of DEGs observed for PA *in vitro* (Fig. S3). Overall, the cytotoxicity testing (Fig. S7) ensured that the mouse exposures reflected physiologically relevant human levels while avoiding human concentrations that are overtly cytotoxic.

PA and IAA had negligible effects on gene expression in the mouse brain, with fewer than ten differentially expressed genes per metabolite (Fig. 3B; Supplementary Table 7). By contrast, TMAO and GDCA each had a significant transcriptional impact in the brain, modulating the expression of 2,305 and 179 genes, respectively (Fig. 3C; Supplementary Table 7; one TMAO-treated mouse was excluded after QC, see Methods). We first evaluated the TMAO results to verify concordance with the literature on this metabolite, which reports predominantly detrimental effects on brain function, with some evidence of potential benefits^48,62^.

In agreement with prior literature^62^, we observed a coordinated downregulation of mitochondrially-encoded genes involved in oxidative phosphorylation, including *mt-Nd1, mt-Nd2, mt-Nd3, mt-Co2, mt-Co3, mt-Cytb, mt-Atp6,* and *mt-Atp8* (Supplementary Table 7). However, gene set enrichment analysis (GSEA) based on hallmark gene sets revealed a more complex rewiring of bioenergetic pathways (Fig. 4A; Supplementary Table 8). Despite the clear downregulation of mitochondrial oxidative phosphorylation, the term “oxidative phosphorylation” was positively enriched in GSEA when comparing TMAO-treated brains to controls. This result was due to the upregulation of several nuclear-encoded genes involved in oxidative phosphorylation (e.g., *Cox4i1, Cox6a1, Cox6b1, Cox7a2l, Ndufa7, Ndufa8, Ndufb2, Ndufb6, Ndufb8, Ndufc2*), perhaps reflecting a compensatory response, similar to that observed in patients with macro-deletion of mtDNA^68^.

**Fig. 4.**
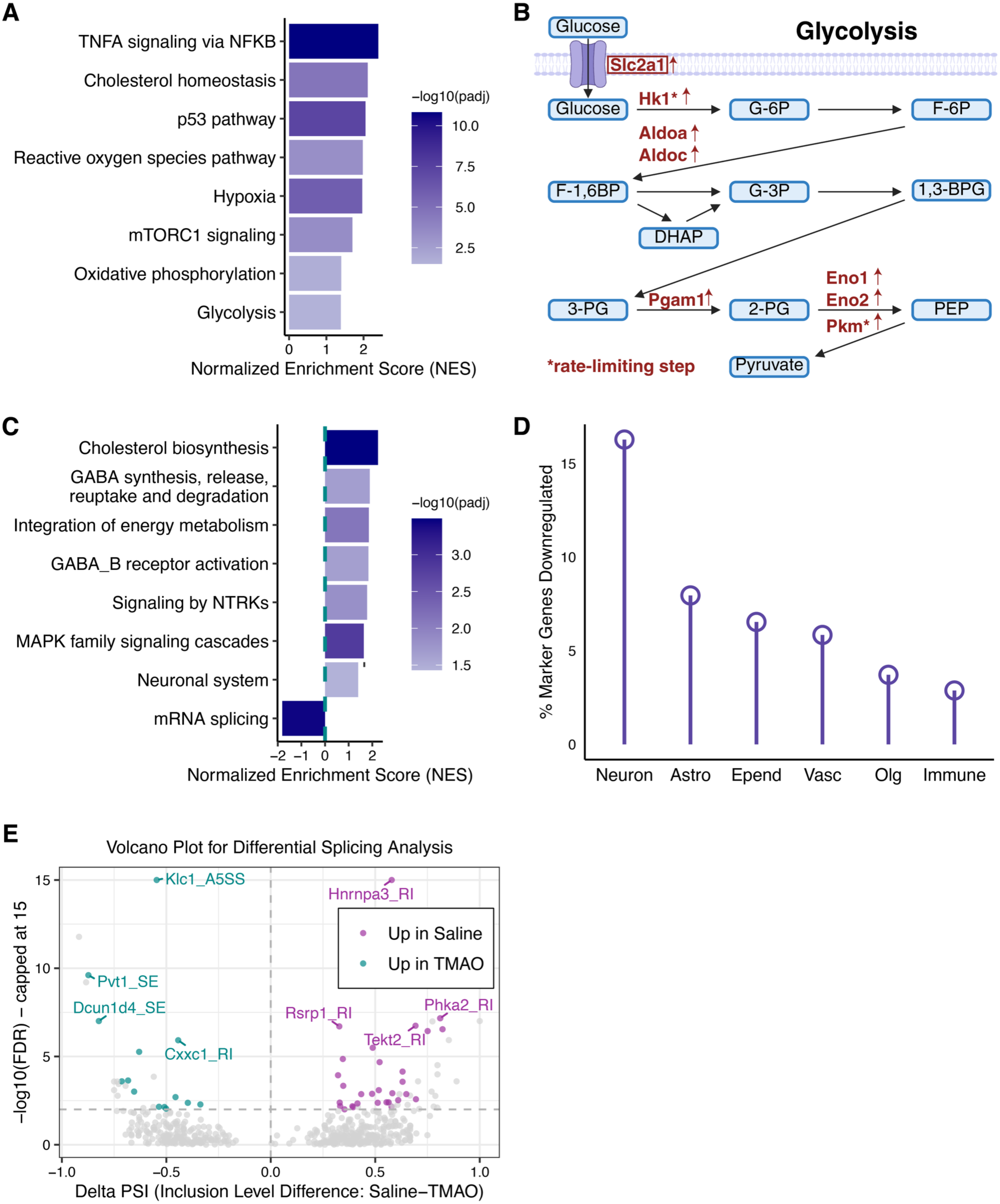
TMAO disrupts transcriptional homeostasis and RNA splicing in the mouse brain. **(A)** Selected significantly enriched terms from gene set enrichment analysis (GSEA) based on hallmark gene sets; full results in Supplementary Table 8. **(B)** The glycolysis pathway, annotated with the glycolytic genes upregulated by TMAO. Asterisks indicate genes whose products catalyze rate-limiting steps of glycolysis. The box around *Slc2a1* denotes its particularly strong statistical significance (*padj* 5.62E-11). Base figure adapted from Hu *et al.*, 2014^86^. **(C)** Selected significantly enriched terms from GSEA using the REACTOME pathway database; full results in Supplementary Table 8. **(D)** Percentage of marker genes in each brain cell class that are downregulated by TMAO. Neuron = neuronal lineage, Astro = astrocyte lineage and stem cells, Epend = ependymal cells, Vasc = vasculature cells, Olg = oligodendrocyte lineage, Immune = immune cells. **(E)** Results of differential splicing analysis comparing brains from TMAO-exposed and control mice. Each point represents a splicing event. The x-axis represents the inclusion level difference (ΔPSI) and the y-axis represents the −log10(FDR), capped at 15 for visual clarity. Events significantly enriched in control or TMAO are shown in purple and teal, respectively, with corresponding metrics listed in Supplementary Table 10. Non-significant events are shown in gray. To reduce visual clutter, points were filtered to those with −log10(FDR) > 0. RI = retained intron, A5SS = alternative 5’ splice site, SE = skipped exon.

In addition, GSEA highlighted the term “glycolysis” as enriched (Fig. 4A). Indeed, eight mouse glycolytic genes were upregulated by TMAO, including *Hk1* and *Pkm*, which encode enzymes that catalyze rate-limiting steps of glycolysis^70^ (Fig. 4B; Supplementary Table 7). Furthermore, *Slc2a1* (i.e., *Glut1*), which encodes a major glucose transporter in the brain that supports glycolytic flux, was the second most significantly upregulated gene by TMAO (adjusted *p*-value 5.62E-11; Fig. 3C and Supplementary Table 7). These findings agreed with data from our transcriptomic screen in hCMEC/D3 cells, where TMAO upregulated human glycolytic genes including *HK1*, *PKM*, *PFKP*, *GPI*, and *ALDOA* (Supplementary Table 5). Overall, our results align with previous findings that TMAO results in mitochondrial dysfunction in the brain^48^. Furthermore, our findings demonstrate transcriptional upregulation of the glycolytic pathway in the brain.

Mitochondrial dysfunction can cause oxidative stress. Consistent with this, the “reactive oxygen species pathway” was significantly more highly expressed (Fig. 4A) in the brain transcriptome of TMAO-exposed animals vs. controls. In addition, *Txnip* - which encodes a known inducer of oxidative stress and activator of the inflammasome^71^ - was significantly upregulated by TMAO (Supplementary Table 7). This finding is consistent with the most enriched GSEA term, “TNFα signaling via NF-кB” (Fig. 4A), which reflects inflammatory signaling. Together, these results support previous studies showing that TMAO promotes oxidative stress and inflammation, including in the brain^42,48,72^.

GSEA using the REACTOME pathway database provided additional insights, including the positive enrichment of genes involved in synaptic transmission and GABAergic signaling, especially genes related to GABA_B_ receptor activation (Fig. 4C; Supplementary Table 8). By contrast, we observed a marked downregulation of genes encoding GABA_A_ receptor subunits (Fig. S8). These results suggest that TMAO may shift inhibitory neural circuits from fast GABA_A_-mediated responses towards slower, GABA_B_-mediated responses. Although previous literature has demonstrated negative effects of TMAO on synaptic function^48^, this is the first observation, to our knowledge, of specific perturbation of GABAergic signaling pathways. To assess the extent of TMAO’s impact on neuronal cells vs. other brain cell types, we computed “marker genes” for six different brain cell classes, using a well-characterized single-cell dataset from the mouse brain^73^ (Methods; Supplementary Table 9). TMAO downregulated over 15% of the marker genes for cells in the neuronal lineage, representing more than twice its impact on any of the other five cell classes (Fig. 4D). Of note, several of the downregulated GABA_A_ receptor genes (e.g., *Gabra1*, *Gabra2*, *Gabrg2*, and *Gabrb3*, see Fig. S8) were among these neuronal markers, with minimal expression in other cell classes (Supplementary Table 9).

GSEA also highlighted the depletion of genes associated with “mRNA splicing” (Fig. 4C). Strikingly, several of the genes that were most significantly downregulated by TMAO (e.g., *Tia1*, *Srrm2*, *Pnisr*, and *Rbm39*, see Fig. 3C) regulate alternative splicing^74–77^. TMAO also significantly downregulated *Leng8*, which normally prevents partially processed transcripts from leaking into the cytoplasm^78^. These results were intriguing given the well-established link between changes in splicing patterns and the development of neurological disease^79^.

We next sought to determine whether any genes were differentially spliced in TMAO-treated animals vs. controls. rMATS-turbo^80^ nominated 126 differential splicing events between TMAO- and vehicle-treated brains, which we then narrowed to 44 using conservative filters (Fig. 4E; Supplementary Table 10; Methods). Manual inspection in Integrative Genomics Viewer (IGV) revealed clear patterns consistent with differential splicing, including events in the RNA processing factors *Rsrp1*, *Cpsf4*, and *Ythdf3*, as well as in *Kmt2b*, a gene whose aberrant splicing has been linked to dystonia^81,82^ (Figs. S9-S10). The *Rsrp1* event - selected because of its relatively high read coverage - was further validated by RT-PCR (Fig. S9). This event may be related to the observed downregulation of *Tia1*, whose human homolog is known to regulate *RSRP1* splicing in HepG2, a human liver cancer-derived cell line (ENCODE file ENCFF258ZJM^83^; RI event #6165), though a causal role for *Tia1* was not demonstrated here.

To more robustly test TMAO’s ability to alter splicing patterns, we performed deeper RNA-seq (targeting 100M reads per sample), in keeping with standard practices in the splicing field. RNA-seq was performed on SH-SY5Y human neuroblastoma cells treated with TMAO (30μM, 3h exposure) or the splicing inhibitor control pladienolide B (100nM, 4h exposure), both with 3 replicates per condition and the appropriate vehicle controls (see Supplementary Table 3).

As expected, pladienolide B caused widespread splicing disruption in SH-SY5Y cells, with rMATS-turbo identifying thousands of differential splicing events under the same conservative filters used for the mouse brain dataset. For instance, *DNAJB1* and *RIOK3* showed increased intron retention, as previously reported in other contexts^84,85^ (Fig. S11; Supplementary Table 11). TMAO treatment produced 101 differential splicing events (Supplementary Table 11), more than twice the number observed in the mouse brain. IGV visualization again showed read-coverage patterns consistent with differential splicing (Fig. S12).

Comparing the rMATS-turbo results for SH-SY5Y cells and mouse brain, TMAO-associated splicing changes appeared largely context- and species-specific (Supplementary Tables 10-11). However, shared features, including altered intron retention in JAK3/Jak3 and events involving RNA processing factors, suggest that some aspects of the splicing response may be conserved or functionally related across systems. Together, these data support a previously unrecognized role for TMAO in modulating alternative splicing patterns, with the full scope and functional impact of these changes remaining to be determined. Overall, our findings confirm^42,48,62^ and broaden current understanding of TMAO’s effects on the brain.

### GDCA remodels chromatin and silences genes that protect the brain from stress and injury

Like TMAO, GDCA is a well-known microbiome-derived metabolite. It is a secondary conjugated bile acid, formed when the host conjugates microbially produced DCA with glycine. Though GDCA has been reported to influence metabolic health with potential beneficial effects^87–89^, its effects on the host are understudied compared to TMAO and are likely context-dependent. Blood levels of GDCA have been associated with neurodegeneration and/or cognitive impairment in at least nine studies^29–37^ (summarized in Supplementary Table 12), suggesting a potential impact on brain function.

GDCA strongly suppressed the expression of many genes in the mouse brain (Fig. 3C; Supplementary Table 7). The most significantly downregulated gene was *Apold1* (log2FoldChange −3.7, adjusted *p*-value 1.82E-108), a gene whose expression is critical for stroke recovery in mice^90^. *Ccn1*, the second most downregulated gene (log2FoldChange −3.0, adjusted *p*-value 3.50E-70), similarly promotes wound healing and tissue repair^91^. This pattern extended beyond individual genes: many of the genes that were strongly downregulated by GDCA (e.g., *Klf2*, *Klf4*, *Adamts1*, *Akap12*, *Maff*, *Plaur*, *Ddit4*, *Atf3*, *Btg2*) are involved in cellular responses to injury and stress, including oxidative stress, hypoxia, DNA damage, and stroke^92–100^. These observations were supported by GSEA based on hallmark gene sets. GSEA demonstrated significant negative enrichment of the terms “TNFα signaling via NF-кB” and “hypoxia” (Fig. 5A), as well as nominal negative enrichment of “p53 pathway” (Supplementary Table 8), consistent with reduced activity of stress-response pathways. These changes could reflect a reduced need for stress signaling; alternatively, they could indicate disruption of pathways needed to maintain cellular homeostasis. Marker gene analysis (Fig. 5B) suggested perturbations to vascular homeostasis in particular. This included downregulation of vascular markers *Klf2* and *Klf4*, which have been shown to control endothelial function and vascular integrity^101,102^.

**Fig. 5.**
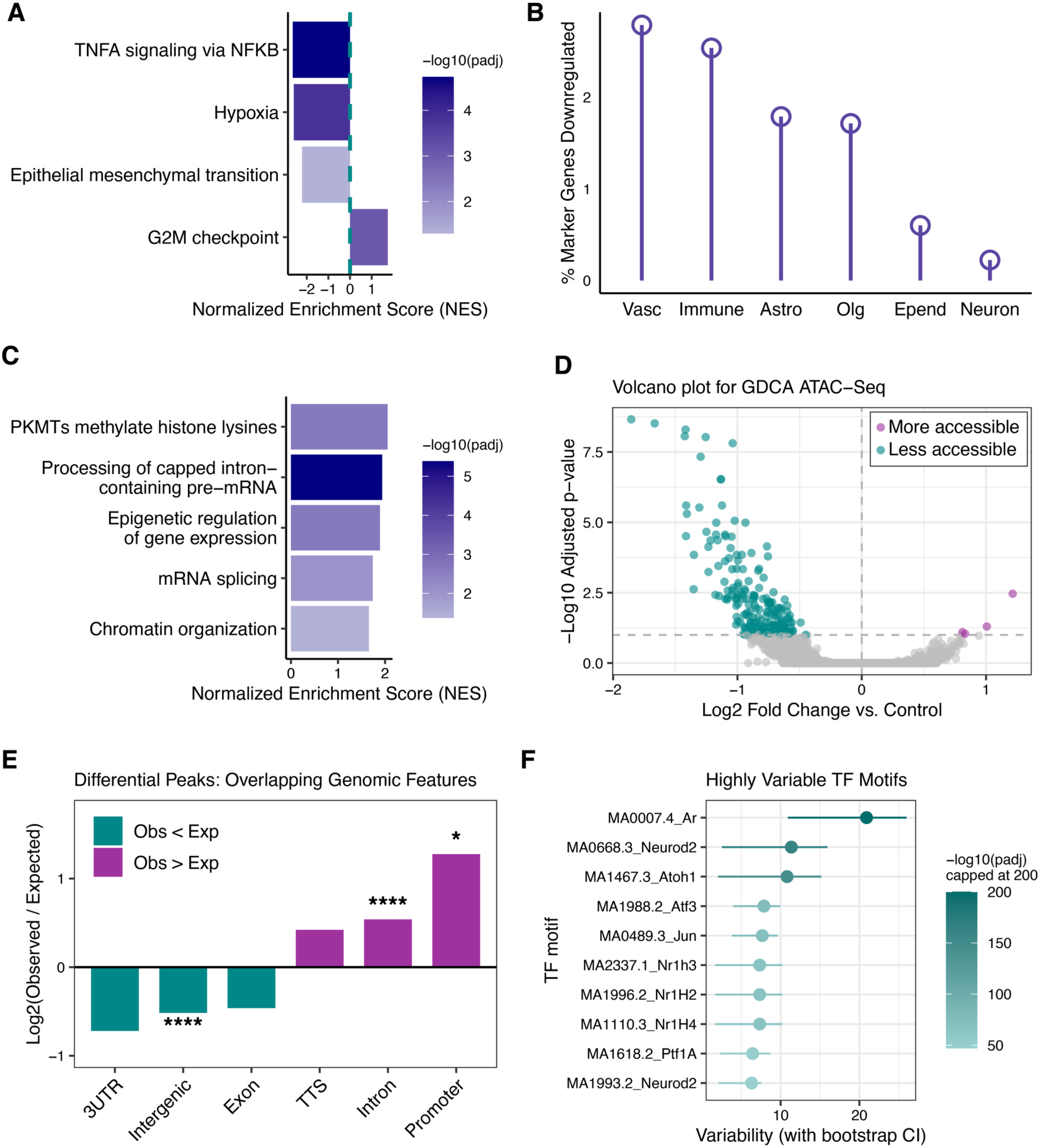
GDCA suppresses stress-response genes and remodels chromatin in the brain. **(A)** All significantly enriched terms from gene set enrichment analysis (GSEA) based on hallmark gene sets; full results in Supplementary Table 8. **(B)** Percentage of marker genes in each brain cell class that are downregulated by GDCA. Vasc = vasculature cells, Immune = immune cells, Astro = astrocyte lineage and stem cells, Olg = oligodendrocyte lineage, Epend = ependymal cells, Neuron = neuronal lineage. **(C)** All significantly enriched terms from GSEA using the REACTOME pathway database; full results in Supplementary Table 8. **(D)** Volcano plot of differential chromatin accessibility between GDCA-treated and vehicle control mouse brains, based on ATAC-seq of flash-frozen right hemispheres from the initial RNA-seq cohort (see Methods). Y-axis: - log10(adjusted p-value) from DESeq2; x-axis: log2FoldChange vs. vehicle control. Purple dots represent genomic peaks with higher accessibility in brains from GDCA-exposed mice (adjusted *p* ≤ 0.1 and positive log2FoldChange). Teal dots represent genomic peaks with lower accessibility in brains from GDCA-exposed mice (adjusted *p* ≤ 0.1 and negative log2FoldChange). **(E)** Observed versus expected proportions of differentially accessible peaks overlapping various genomic features. Expected proportions were calculated based on the reference genome. Introns and promoters are significantly enriched among differential peaks, whereas intergenic regions are depleted. The symbol * indicates *p* ≤ 0.05 and **** indicates *p* ≤ 0.0001. *p*-values were calculated by HOMER using a one-sided binomial test (upper-tailed when observed > expected and lower-tailed when observed < expected). **(F)** Transcription factor (TF) motifs with the highest variability in accessibility across ATAC-seq samples, as quantified by chromVAR^109^. Dots represent point estimates of motif variability and horizontal lines indicate the corresponding 95% bootstrap confidence intervals. The color scale shows the significance of variability as −log10(adjusted *p*-value from chromVAR). To facilitate visualization, −log10(padj) values were capped at 200; the true padj for the variability of MA0007.4_Ar across samples is 0 (Supplementary Table 13).

GSEA using the REACTOME pathway database (Fig. 5C; Supplementary Table 8) demonstrated the positive enrichment of several terms related to chromatin dynamics, including “PKMTs methylate histone lysines,” “epigenetic regulation of gene expression,” and “chromatin organization.” Based on this result, we postulated that GDCA achieves its notable gene regulatory effects, including the strong silencing of many genes, by inducing changes in chromatin organization and accessibility. To test this hypothesis, we conducted bulk ATAC-Seq on the flash-frozen right brain hemispheres of the GDCA- and vehicle-treated mice.

Consistent with our prediction that GDCA induces chromatin changes, ATAC-seq identified hundreds of genomic regions with differential accessibility between the brains of GDCA-and vehicle-treated mice (Fig. 5D; Fig. S13A-B; Supplementary Table 13). 203 of the 207 differentially accessible peaks (98%) were less accessible in the GDCA-exposed brains. Moreover, using rGREAT to annotate differential peaks with genes whose regulatory domains they overlap, we found that eleven peaks overlapped regulatory domains of genes that were also differentially expressed in RNA-seq (Supplementary Table 13). These genes included several known stress-response genes: *Ddit4*^97^, *Errfi1*^103^, *Rhob*^104^, and *Sgk1*^105^. In 10/11 cases, the direction of change was concordant across datasets - decreased gene expression and decreased accessibility of the associated peak - supporting the idea that the strong transcriptional changes induced by GDCA are driven, at least in part, by chromatin remodeling.

We further annotated the differentially accessible peaks based on overlapping genomic features, and compared the proportion of peaks in each category to the proportions expected from the reference genome, using HOMER^106^. This analysis revealed enrichment of differential peaks in promoters and introns (Fig. 5E; Fig. S13C; Supplementary Table 13). Promoters and introns frequently harbor regulatory elements; for example, introns may contain enhancers^107^. Therefore, the enrichment of these features among differential peaks further supports a direct link between GDCA-induced changes in chromatin accessibility and effects on gene expression. Consistent with this interpretation, 4 of the 5 most significantly differential intronic peaks fully overlapped elements annotated as distal enhancers in the ENCODE SCREEN database (https://screen.wenglab.org/)^108^.

Promoters and enhancers commonly contain motifs that bind transcription factors (TFs) critical for the regulation of gene expression. We therefore used chromVAR^109^ to assess whether GDCA altered the accessibility of any TF motifs from the “CORE” collection of the well-established JASPAR^110^ database. We found that TF motif accessibility patterns quantified by chromVAR clearly separated GDCA-exposed brains from controls (Fig. S14A). To identify motifs contributing to this separation, we next examined chromVAR “variability” scores for each motif, where a value near 1 indicates that motif-associated peaks are no more variable across samples than background peak sets matched for GC content and average accessibility. The TF motif with the highest variability across samples was MA0007.4, which is recognized by the *Ar* transcription factor that coordinates the cellular response to androgens (Fig. 5F; Supplementary Table 13). This *Ar* motif was consistently less accessible in the GDCA-exposed brains than in controls (Fig. S14B).

A recent study showed that certain microbiome-derived bile acids, which share the sterol structure of androgens, can antagonize the human androgen receptor^111^. This is in addition to their established roles as ligands for other receptors, including FXR and TGR5^112^. Thus, GDCA may block mouse *Ar* from nuclear translocation and binding to its target motifs, thereby reducing motif accessibility - since TF binding usually opens chromatin^113^ - and suppressing androgen-responsive transcription. Consistent with this model, several of the genes strongly downregulated by GDCA (e.g., *Adamts1*, *Akap12*, *Sgk1*) belong to the “Androgen Response” hallmark gene set from MSigDB. GSEA likewise showed negative enrichment of this gene set (normalized enrichment score = −2.17; Supplementary Table 8), although this did not reach statistical significance after multiple testing correction (raw *p* = 0.05, adjusted *p* = 0.16), possibly reflecting a lag between certain chromatin changes and downstream transcriptional effects.

### GDCA acutely reduces locomotion in male but not female mice

Growing evidence indicates that androgens support both cognitive and motor function^114–116^. In addition, increased chromatin accessibility can be beneficial for cognitive function^117^, in stark contrast to the accessibility reductions observed after GDCA treatment. Therefore, we hypothesized that GDCA impairs cognitive and motor performance and may also influence additional aspects of mouse behavior.

To test this, we first administered GDCA for three days (0.4 mg/kg/day, intravenous retro-orbital injection) and assessed behavior. There were no detectable differences between GDCA-and vehicle-treated mice, as assessed in an activity chamber on day 4 or in a Y-maze on day 5 (*n =* 15 adult male mice per group; see Fig. S15 and Supplementary Table 14). We reasoned that the administered GDCA would likely be cleared by the day of testing (e.g., via hepatic uptake^118^), such that systemic levels were no longer elevated. Therefore, in a second cohort of adult male mice, we evaluated the impact of acute exposure. Here, GDCA was administered via oral gavage, because retro-orbital injection, which is invasive, is not compatible with same-day behavioral testing. Gavage doses (10 or 50 mg/kg) were based on prior mouse studies involving GDCA^87,119^ and previous estimates that only a small percentage (5-10%) of bile acids escape the enterohepatic circulation^88,112^.

One hour after administration of GDCA or vehicle, mice were tested in the Y-maze (*n =* 15 per group). Like vehicle-treated mice, GDCA-treated mice alternated between maze arms above chance levels (Fig. S16A; Supplementary Table 14), indicating no impairment in spatial working memory. However, GDCA reduced locomotion in a dose-dependent manner, as evidenced by fewer entries into maze arms and lower total distance traveled within the Y-maze (Figs. 6A-D; Supplementary Table 14). Statistical significance was reached at 50 mg/kg, accompanied by detectable brain levels of GDCA and a reduction in core body temperature (Figs. 6C-E; Supplementary Tables 14-15). Total distance traveled in the Y-maze also trended towards a GDCA-induced reduction within the setup of cohort 1 (*p*=0.054; Fig. S15K; Supplementary Table 14).

**Fig. 6.**
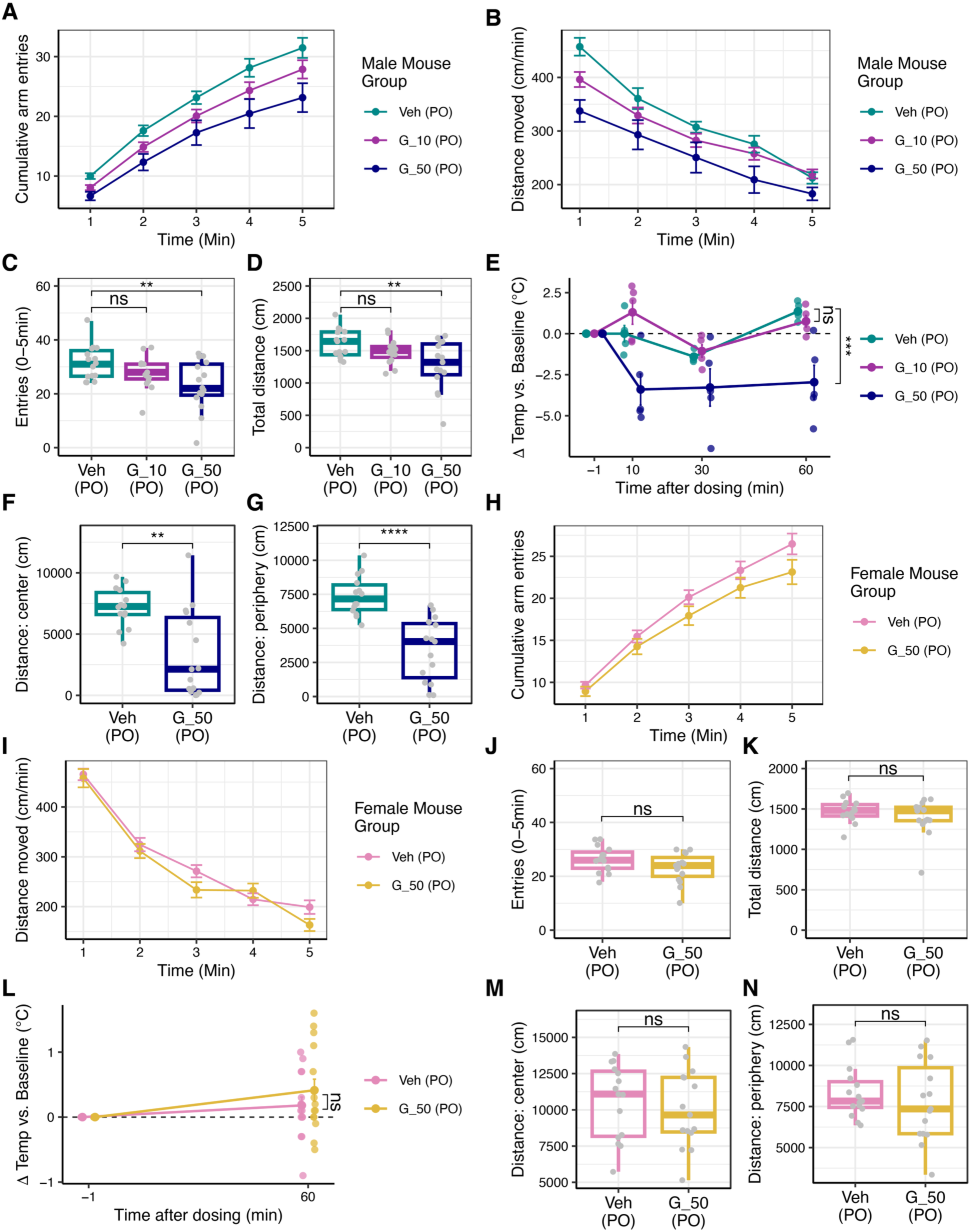
Oral GDCA acutely reduces locomotion in male mice. Panels (A)-(G) show experiments in male mice, and panels (H)-(N) show analogous experiments in female mice. In all panels, Veh (PO), G_10 (PO), and G_50 (PO) indicate oral gavage with vehicle, 10 mg/kg GDCA, and 50 mg/kg GDCA, respectively. PO = *per os*. **(A)** Cumulative arm entries in a Y-maze over five minutes, measured one hour after gavage (*n =* 15 per group; lines show group means ± SEM at each time point). **(B)** Distance moved per minute in the Y-maze, plotted as in (A). **(C)** Boxplot of total arm entries in the Y-maze. Groups were compared by one-way ANOVA with Dunnett’s post hoc test (statistics shown). **(D)** Boxplot of total distance traveled in the Y-maze. Statistics as in (C). **(E)** Change in body temperature after gavage; lines show group means ± SEM at each time point, and points show individual mice. All *n =* 15 mice per group were measured at baseline (time −1); *n =* 5 mice per group were measured at each subsequent time point. At the 60-minute time point, groups were compared by one-way ANOVA with Dunnett’s post hoc test (statistics shown). **(F)** Total distance moved in the center of an activity chamber over one hour, beginning one hour after gavage (*n =* 14 mice in vehicle group, *n =* 15 mice in GDCA group; see Methods). Groups were compared using Welch’s two-sample, two-sided *t*-test. **(G)** Same as (F), but for the periphery of the activity chamber. **(H)**-**(N)** are analogous to (A)-(G), respectively, for female mice; displayed comparisons were performed using Welch’s two-sample, two-sided *t*-test, and sample size was *n =* 15 per group. In all panels, ns indicates not statistically significant, * indicates *p* ≤ 0.05, ** indicates *p* ≤ 0.01, *** indicates *p* ≤ 0.001, and **** indicates *p* ≤ 0.0001. Full statistics are reported in Supplementary Table 14. In all boxplots, boxes represent the interquartile range (IQR), horizontal lines indicate medians, and whiskers extend to the most extreme values within 1.5 x IQR of the hinges.

We confirmed GDCA’s ability to reduce locomotion in a third cohort of adult male mice. Mice gavaged with 50 mg/kg GDCA or vehicle (*n =* 14-15 per group, see Methods) were assessed in an activity chamber one hour after dosing. GDCA-treated mice exhibited reductions in distance traveled and related activity measures compared with vehicle-treated controls (Fig. 6F-G; Fig. S16B-D; Supplementary Table 14). Because bile acids can induce diarrhea, we also measured fecal pellet dry weight from pellets collected during the activity chamber assay in a subset of mice (*n =* 4 per group). Percent dry weight did not differ between GDCA- and vehicle-treated mice (Fig. S16H; Supplementary Table 14), and pellets appeared visually comparable between groups, indicating no detectable change in pellet consistency under these conditions.

To determine whether reduced locomotion reflected impaired motor coordination, we tested trained mice on a rotarod. Mice completed three 40s trials: 16rpm at 15min post-dose, 32rpm at 15min post-dose, and 16rpm at one hour post-dose. In both 16rpm trials, all mice remained on the rod for the full 40s trial, regardless of treatment group (Fig. S16E-F). By contrast, during the more challenging 32rpm trial, 8 of 15 GDCA-treated mice fell from the rod, compared with 2 of 14 vehicle-treated mice (Fig. S16G). This difference trended towards statistical significance by Fisher’s exact test (*p* = 0.0502; Supplementary Table 14), with latency to fall showing a similar trend (*p* = 0.057; Supplementary Table 14). These results suggest that GDCA may mildly affect motor coordination or balance, although additional testing will be required to confirm this effect.

Because all previous assays were conducted in male mice, we next asked whether GDCA’s effects were sex-specific, particularly given the proposed mechanism involving antagonism of the androgen receptor. In adult female mice treated with 50 mg/kg GDCA or vehicle (*n =* 15 per group), we observed no significant differences in Y-maze behavior, activity chamber measurements, rotarod performance, core body temperature, or fecal pellet percent dry weight (Fig. 6H-N; Fig. S16I-P; Supplementary Table 14). The absence of detectable effects in females could reflect lower baseline androgen signaling, reduced brain penetration of GDCA (Supplementary Table 15), or other sex-dependent factors.

## Discussion

The gut microbiome produces and impacts the systemic concentration of hundreds of metabolites, each of which may profoundly influence host organs, including the brain. However, systematic, high-throughput efforts to identify metabolites relevant to gut-brain signaling are lacking, partly because testing hundreds of molecules in animal models - the most accurate system to assess brain effects - is impractical. In this study, we integrated computational, *in vitro*, and *in vivo* approaches to identify age-related, microbiome-derived blood metabolites that - at physiologically relevant levels - induce significant molecular changes in the mouse brain. Among other changes, we observed impacts on RNA splicing and chromatin accessibility, both of which are altered in brain aging and in age-related neurodegeneration^79,120,121^. Furthermore, we found that a microbiome-derived secondary bile acid, GDCA, can reduce locomotor activity, which often declines with aging and neurodegenerative disease. These findings indicate that “microbiome aging” may modulate “brain aging” through impacts on the blood metabolome, and that targeting the microbiome and its metabolites may offer cognitive benefits during aging.

The aminoxide TMAO has been extensively studied in the context of cardiovascular disease^15,122^, and there is a growing body of research on its impact on the brain, where it has been shown to induce mitochondrial dysfunction, oxidative stress, neuroinflammation, and reductions in synaptic plasticity, although it may also confer beneficial effects on the blood-brain barrier^48,62^. Since TMAO is very likely to cross the blood-brain barrier^123^, its effects on the CNS may be mediated through direct central action. We found that TMAO also upregulates the glycolytic pathway in the brain, and impacts RNA splicing by inducing strong downregulation of splicing factors and driving dozens of differential splicing events. Although prior work has identified splicing changes in the brains of germ-free vs. conventionally raised animals^124^, our work provides the first evidence - to our knowledge - that specific microbiome-derived metabolites can alter host splicing patterns, whether in the brain or in any other host organ. While not necessarily surprising - given that splicing is responsive to a wide range of perturbations - this phenomenon is notably underexplored. Indeed, GDCA may also affect splicing patterns in the brain (Fig. 5C), raising questions about the relative impacts of individual metabolites and how they might converge to influence brain function.

Unlike TMAO, little is known about the effects of GDCA on the mammalian nervous system, with prior knowledge largely limited to its association with neurodegenerative disease and cognitive impairment in several human cohorts^29–37^. Here, we find that GDCA reduces chromatin accessibility at hundreds of genomic regions in the mouse brain, potentially explaining how it also silences genes required for stress and injury responses. Although chromatin accessibility changes in the aging and diseased brain are complex and nuanced^120,125^, large decreases in chromatin accessibility, such as those induced by GDCA, may be detrimental to neuronal stress resilience and cognitive function^117,126^. Consistent with this possibility, a very recent independent study reported that GDCA can impair recognition memory in mice^127^. There is precedent for microbiome-driven changes in the brain epigenome, for example, germ-free status is known to have an effect^128^. Previous metabolite-specific work has focused on the impact of short-chain fatty acids^129^, leaving open questions about the many other classes of microbial effectors, including bile acids.

Our data suggest that the effects of GDCA on the brain transcriptome and epigenome may be partially mediated by antagonism of the host androgen receptor (Figs. 5F and S14). We did not test this mechanism directly using an androgen receptor knockout model, but it is supported by prior work showing that related bile acids can antagonize the host androgen receptor^111^. It is also consistent with literature showing that GDCA improves phenotypes related to polycystic ovary syndrome, a condition that involves androgen excess^119^. In our study, GDCA reduced locomotion in male mice, consistent with reduced androgen signaling^115,116^, but did not impact locomotion in females, perhaps because females have lower baseline androgen levels. Lower brain penetration of GDCA in females may also contribute to this result (Supplementary Table 15). Further work will be needed to fully elucidate how GDCA affects cognition and motor behavior in each sex, whether these effects depend on androgen receptor signaling, and whether they arise through central or peripheral mechanisms.

Although our work clearly demonstrates that circulating microbiome-derived metabolites can induce significant molecular alterations in the brain, it has several limitations. First, we tested only short-term exposures. However, brain aging and neurodegeneration occur over long periods of time. It will thus be important to decipher the short- vs. long-term effects of metabolites of interest. *In vitro*, this may require determining the half-life of each metabolite and ensuring continuous exposure to the desired concentration. *In vivo*, metabolites may need to be administered via osmotic minipump or through the diet. To determine the most physiologic dosing regimens, pharmacokinetic (PK) studies - involving metabolomics at various timepoints - will be necessary. Second, the modest effects observed for PA and IAA in our acute-exposure experiments (Fig. 3B) should be interpreted with caution. These findings may simply reflect suboptimal bioavailability of the administered formulations, further underscoring the value of PK studies. In addition, because our omics analyses were powered primarily to detect moderate effects, limited statistical power may have precluded detection of smaller effects across all metabolites, including PA and IAA. Additionally, while we find interesting molecular phenotypes in adult mice, these results may or may not translate to aged animals. Future *in vivo* work in aged mice will be informative, in this regard, given that age may modify the brain’s response to each metabolite. Age-related comorbidities such as diabetes and cardiovascular disease may also impact host response to individual metabolites, highlighting the importance of testing metabolite effects in the relevant disease models. Finally, because our multi-omic analyses were conducted using bulk rather than single-cell methods, the cell-type-specific effects of TMAO and GDCA remain unclear.

Despite these limitations, our work underscores that the gut microbiome, which is potentially modifiable, is a valuable therapeutic target in the context of brain aging. We identify specific age-associated microbiome metabolites that impact the transcriptional and epigenetic landscape of the brain, with likely consequences for brain function. In addition, although we focused on brain aging in this work, the framework we describe - beginning with metabolomics data analysis, followed by *in vitro* transcriptomic screening, and concluding with *in vivo* studies of host organs and behavior - can be applied to identify the microbiome metabolites relevant to any host biology or pathology of interest, providing a powerful tool for the field of microbe-host interactions.

## Methods

### Identifying microbiome-associated metabolites whose blood levels change with age

To identify metabolites whose blood levels change with aging, we leveraged data from three large cohort studies^23–25^ listed in a recent review^22^ on the metabolomics of aging. From Menni *et al*.^23^, we retained all metabolites from Table S2, since only metabolites significantly associated with age were included in this table. From Dunn *et al*.^24^, we retained metabolites with p_adj <= 0.05 under the test labeled “_age.g2” in Supplementary Material 2 (“AGE” tab). From Darst *et al*.^25^, we retained metabolites with p_age <= 0.05 in Table S1.

Separately, we curated a list of 329 human-associated metabolites that have been reported to be produced by microbes and/or are the products of host-microbe cometabolism. To create the list, we first downloaded Table S3 from a recent study on interactions between human-associated metabolites and GPCRs^26^, and filtered to metabolites that included “Bacteria” in the “Source” column. After filtering, there were 320 metabolites. We then added the following nine metabolites known to be microbially producible and/or producible through host-microbe cometabolism^28,41,130–132^: acetic acid, butyric acid, indole-3-carboxaldehyde, indolelactic acid, enterolactone, imidazole propionate, p-cresol sulfate, hippuric acid and indoleacetic acid.

Then, we intersected the age-associated metabolites from each study with the curated list of metabolites associated with microbial metabolism. The intersection method was different per study given the differing formats of the tables. Specifically, for Darst *et al.*, we were able to intersect based on HMDB ID. We also noted that there was one metabolite that matched on CAS number but not on HMDB ID, given that it was listed as “ornithine HCl” in our curated list of microbiome-associated metabolites, but simply “ornithine” in the Darst study. So, we added ornithine manually to the intersection output. For the Menni and Dunn datasets, which lacked standardized identifiers, we used fuzzy string matching (via the Python package fuzzywuzzy, v0.18.0) to identify metabolites with similar names, followed by manual curation to confirm true matches. We also added two intersecting metabolites that were missed by fuzzy matching - indoleacetate and hippurate.

This process yielded 114 overlapping metabolites, 79 of which were unique based on HMDB ID. To narrow the list to metabolites most strongly associated with the gut microbiome, we leveraged two additional resources: **(1)** a recent study comparing blood metabolite levels between germ-free and conventionally raised mice^14^; and **(2)** the GUTSY Atlas^27^, which reported the proportion of variance in blood metabolite levels “explained” by the gut microbiome for 1,168 metabolites based on paired stool-blood data for over 8,000 humans. Metabolites were retained for the final panel if they met at least one of two criteria: **(1)** showed significantly different blood levels between germ-free and conventionally raised mice^14^; **(2)** ranked in the top 20% of blood metabolites for the GUTSY variance-explained metric. The 20% threshold was chosen to yield a workable number of metabolites for downstream experiments. We eliminated alanine and uridine, the former due to its minimal expected impact on cellular processes^38^, and the latter because its blood levels increased with age in one study^23^ but decreased with age in another^25^.

The final table of 30 metabolites and their properties is contained in Supplementary Table 1. The table also specifies the exact product that was ordered for testing in the *in vitro* transcriptomic screen.

Of note, at a later date, after conducting the *in vitro* RNA-seq experiments, we discovered that the p_age values in Table S1 of the Darst study had not been corrected for multiple hypothesis testing. However, even after we manually corrected for multiple hypothesis testing (using a False Discovery Rate approach in R v4.3.2), all the metabolites from the Darst study in our final panel maintained significance using an FDR threshold of padj ≤ 0.1, with all but cholate (padj = 0.065) and glycodeoxycholate (padj = 0.061) also having a padj ≤ 0.05.

### Cell culture

All cultures were maintained in a 37°C, 5% CO_2_ incubator on tissue culture-treated plates. Testing for mycoplasma contamination was conducted at six-month intervals during periods of active culture using the ATCC Universal Mycoplasma Detection Kit (ATCC, Cat #30-1012K).

Human cerebral microvascular endothelial cells (hCMEC/D3; Sigma, Cat #SCC066) were obtained as a gift from the lab of Prof. Mark Kay at Stanford University. Cell line authentication was conducted using the ATCC Human Cell STR Profiling Service prior to initiating the experiments described in this manuscript. The cell medium was prepared from the EGM2-MV BulletKit (Lonza, Catalog #CC-3202). The VEGF supplement was excluded from the medium based on literature precedent^41,62^ and due to VEGF’s ability to impair the key barrier-forming properties of the cells^133^.

Human neuroblastoma cells (SH-SY5Y; ATCC, Cat #CRL-2266) were obtained as a gift from the lab of Prof. Aaron Gitler at Stanford University. Cell line authentication was conducted using the ATCC Human Cell STR Profiling Service prior to initiating the experiment described in this manuscript. The cell medium was DMEM/F-12 with GlutaMAX™ supplement (Thermo Fisher, Cat #10565018) with 1% Penicillin/Streptomycin (Cytiva, Cat #SV30010) and 10% Fetal Bovine Serum (Thermo Fisher, Cat #A5209401).

### *In vitro* transcriptomic screen - experimental details

The screen was conducted in four separate batches using 12-well plates. Prior to cell seeding, each well was coated with 0.5ml of a 1:20 dilution of rat tail collagen (Sigma, Catalog #08-115) in DPBS (Fisher, Catalog #MT21031CV). Collagen was incubated on the plates at 37°C for 1 hour and rinsed off twice with DPBS before cell seeding. hCMEC/D3 cells were seeded at passages #9-10, at a density of 100,000 cells per well. Seeded cells were grown for six days prior to treatment, based on prior literature^41,62^. The cell culture medium was refreshed every two days.

Each 12-well plate was organized to include three wells for vehicle controls and nine wells for testing three metabolites, with three biological replicates per metabolite. All metabolites tested on a given plate were dissolved in the same solvent (H_2_O or DMSO) and vehicle controls included an equivalent volume (1%) of the solvent in the final media used for the assays. Because urate was insufficiently soluble in H_2_O or DMSO alone, the final treatment media for this metabolite included a small percentage (approximately 0.1%) of NaOH in addition to 1% H_2_O.

Each batch consisted of three 12-well plates, which allowed room for testing nine metabolites per batch. To allow for inclusion of an LPS control condition per batch, eight candidate metabolites were tested per batch. Specifically, batches #2-4 tested 8 metabolites each and batch #1 tested 6, to cover the entire set of 30 metabolites (see Supplementary Table 3). Cells were treated with a metabolite, vehicle, or LPS for 3h before being harvested in TRIzol and stored at - 20°C for downstream RNA extraction and sequencing.

For each metabolite, we determined a physiologically relevant concentration to add to cells (Supplementary Table 1), primarily based on information listed in the Human Metabolome Database (HMDB) as of November 2023. Specifically, for each metabolite, we searched for the metabolite in HMDB, clicked the “Concentrations” tab of the relevant entry, examined the concentrations listed for “normal” adult blood samples (which are recorded in Supplementary Note 1), and selected a representative concentration based on those listings. HMDB did not provide blood concentrations for four metabolites: glycodeoxycholate, imidazole propionate, methyl indole-3-acetate, and N-acetyl-L-alanine. For glycodeoxycholate, we chose the concentration based on Table S2 of Qing *et al.*, *Schizophrenia* 2022^134^. For imidazole propionate, a relevant physiological concentration was determined based on Fig. 1 of Molinaro *et al.*, *Nature Communications* 2020^130^. For methyl indole-3-acetate, we were not able to find literature on the exact concentrations in adult blood samples. We tested 1µM because it was between previously reported concentrations of the metabolite in adult urine^135^ and the chosen test concentration of the related metabolite indole-3-acetate. For N-acetyl-L-alanine, we tested 3µM based on previous literature that included *in vitro* experiments with this metabolite (Yang *et al.*, *Int J Mol Sci* 2023; Section 2.2^136^).

RNA was extracted using the following protocol. Samples were thawed to room temperature and incubated for an additional five minutes after thaw. 200µl chloroform was then added per 1ml of TRIzol used; then, samples were briefly vortexed and incubated 2-3 minutes at room temperature, prior to a 15-minute centrifugation at 17,000 x g at 4°C. The aqueous phase was transferred to a new tube. Steps 2-7 of the “Part 1” quick-start protocol for the RNeasy Mini Kit (Qiagen, Catalog #74106) were then executed, skipping the on-column DNase digestion but including the optional dry spin. DNase digestion was conducted using TURBO DNase (Fisher Scientific, Catalog #AM2239). Then, the “RNA cleanup” section of the RNeasy Mini handbook was executed. Finally, an ethanol precipitation was conducted for all samples to clean up any ethanol contamination, based on the relevant protocol from Cold Spring Harbor Laboratories^137^. RNA samples were sent to Novogene for quality control, mRNA library preparation (poly A enrichment), and sequencing (NovaSeq 6000, 150bp paired end reads, 6Gb raw data per sample).

### *In vitro* transcriptomic screen - computational analysis

Computational analysis of the *in vitro* RNAseq data was performed separately for each sequencing batch, as follows. First, FASTQ files from Novogene were trimmed to remove adapter and low-quality bases using TrimGalore v0.6.10.

Trimmed paired reads were pseudoaligned to the human reference transcriptome using kallisto v0.48.0^138^. The reference transcriptome was downloaded from Ensembl (release 110, Homo_sapiens.GRCh38.cdna.all.fa.gz) and an index was built using the kallisto index command. Samples were then grouped based on the solvent included in the media (DMSO or H_2_O), and a principal component analysis (PCA) was performed separately for each solvent group, as follows. First, transcript abundance tables generated by kallisto were normalized and filtered using the kallisto_table function from the sleuth R package (v0.30.2), using the options normalized = TRUE and use_filtered = TRUE. The resulting tables were log-transformed using the natural logarithm of (x+0.5) prior to PCA, which was conducted using the prcomp function in R v4.3.2.

PCA plots were used to identify potential outlier samples within each group - defined as samples positioned far from: (1) the vast majority of samples in the plot, and (2) the other replicates within their treatment condition. Due to the subjective nature of visual outlier identification, samples were only excluded from downstream analyses if their removal was additionally supported by deviant quality control (QC) statistics from Novogene. **Note:** The PCA plots for each sequencing batch are displayed in Figs. S4 and S5, and the corresponding sample lists and QC statistics are contained in Supplementary Table 3.

Based on the above criteria, the following 5 (out of 138) samples were excluded from downstream analysis. From batch 2, oxalate replicate #1 and carnitine replicate #2 were removed (supporting QC statistic: low %GC content compared to the other samples in the batch). From batch 3, glycodeoxycholate replicate #1 and quinolinate replicate #2 were removed (supporting QC statistic: low %GC content compared to other samples in the batch). From batch 4, DMSO control replicate #5 was removed (supporting QC statistic: low RNA concentration compared to all other DMSO control samples).

Following PCA and outlier removal, DESeq2^139^ v1.42.1 was used to determine differentially expressed genes between each set of metabolite-treated samples and the corresponding vehicle controls in the sequencing batch, using the default significance cutoff of alpha=0.10. As specified in Supplementary Table 3, the number of vehicle controls per solvent per sequencing batch ranged from three to nine. Gene abundance values were determined from the per-transcript kallisto outputs using tximport (v1.31.0). When applicable, the corrplot R package (v0.95) was used to verify that all vehicle controls of the same solvent (e.g., 1% DMSO controls) on different plates could be grouped, based on strong correlations - Pearson’s *r* > 0.98- of log-transformed gene abundances.

As recommended by the DESeq2 developers, the function lfcShrink^69^ was applied to conservatively shrink log2 fold changes, using the shrinkage estimator ‘apeglm.’ Additionally, to avoid overcounting the number of significant hits per metabolite, we filtered the DESeq2 output of differentially expressed genes to eliminate lines that were exact duplicates of previous lines besides the Ensembl ID field, which can occur when two different Ensembl gene IDs are mapped to the same physical gene. Finally, genes with an absolute shrunken log2 fold change ≤ 0.1 were not counted as DEGs, even if they met the significance threshold of adjusted *p* ≤ 0.1.

The KEGG pathway enrichment analyses shown in Figs. 2B and S2 were conducted using g:Profiler (https://biit.cs.ut.ee/gprofiler/gost)^140^ on August 17, 2025 using the default g:SCS significance threshold and a custom list of background genes - specifically, all genes that passed DESeq2’s internal filters in the corresponding DESeq2 analysis. For the analysis of DEGs in the Alzheimer and Parkinson’s KEGG pathways (Fig. 2C), the pathway gene lists were retrieved on July 19, 2025 from https://rest.kegg.jp/link/hsa/path:hsa05010 and https://rest.kegg.jp/link/hsa/path:hsa05012, respectively. The numbers reported in Fig. 2C represent numbers of unique genes (based on Entrez ID).

### *In vitro* cytotoxicity testing

hCMEC/D3 cells (passage #22) were seeded on Day 0 at a density of 100,000 cells per well in 12-well plates. Prior to seeding, each well was coated with 23μg rat tail collagen (Sigma, Cat. #08-115) in 1mL DPBS (Fisher, Catalog #MT21031CV) and rinsed twice with DPBS following a 10-minute incubation at room temperature.

On Day 2, cells were treated for 3 hours with EGM2-MV BulletKit medium (Lonza, Catalog #CC-3202) excluding the VEGF supplement and supplemented with either: **[1]** metabolites at the concentrations shown in Fig. S7, **[2]** vehicle control (i.e., 1% DMSO or 1% H_2_O), or **[3]** 20% DMSO (cytotoxic control).

Following treatment, cells were detached with 0.05% trypsin-EDTA (Fisher Scientific, Catalog #25-300-054), combined with the original well media to retain non-adherent or dead cells, pelleted by centrifugation (100 x g, 5min, 4°C), and resuspended in 150μl serum-free, phenol red-free EGM2-MV medium (basal medium: Lonza, Catalog #CC-3129; supplements: Lonza EGM2-MV SingleQuots kit, Catalog #CC-4147, excluding FBS and VEGF). 30μl of Propidium Iodide ReadyFlow reagent (Thermo Fisher, Catalog #R37169) was added to the suspension, and fluorescence was measured using a NovoCyte Quanteon flow cytometer at the Stanford Shared FACS Facility.

### Animals

Male and female C57BL/6 mice (3-4 months old for all experiments) were obtained from Jackson Laboratories (stock # Jax 000664). All mice were kept on a 12 h-12 h light-dark cycle and provided *ad libitum* access to food and water. All animal care and procedures complied with the Animal Welfare Act and were in accordance with institutional guidelines and approved by the institutional administrative panel of laboratory animal care at Stanford University (PIs of animal protocols: T. Wyss Coray, M. Shamloo; protocol #s 29100 and 18466).

### Mouse brain transcriptomics after metabolite injection

Guided by prior studies investigating TMAO and the microbiome metabolite p-Cresol sulfate^41,62^, we conducted bulk transcriptomics on mouse half-brains following a 2-hour exposure to each metabolite of interest or the corresponding vehicle control. As detailed in Figs. 2-3 and the accompanying text, four metabolites were tested in mice - palmitic acid (PA), indoleacetic acid (IAA), trimethylamine N-oxide (TMAO), and glycodeoxycholate (GDCA). Intravenous retro-orbital injection under isoflurane anesthesia was used as the administration route.

Metabolite dosing was based on precedent from the study on TMAO that guided experimental design^62^. For TMAO (Cayman Chemicals, Item #17354, Batch #0522898-67), we used the same dose as the original study, 1.8 mg/kg. This dose was described as “calculated to approximate human circulating TMAO levels.” Thus, for the remainder of the metabolites, we examined the targeted metabolomics literature^31,33,66,67,141,142^ to identify the range of circulating levels previously observed in human seniors, and calculated mouse-equivalent doses of observed human concentrations, using well-established guidelines^143,144^. PA (Sigma, Catalog #P0500-10G, Lot #0000253065) was administered at a final dose of 22.2 mg/kg (converted from ∼100µM human circulating concentration). IAA (Sigma, Catalog #I2886-5G, Lot #0000249926) was administered at a final dose of 1.5 mg/kg (converted from ∼10µM human circulating concentration). GDCA (Sigma, Catalog #361311-5GM, Lot #4045885) was administered at a final dose of 0.4 mg/kg (converted from ∼1µM human circulating concentration).

Two experiments were conducted in total. **Experiment 1** contained 8 mice; 4 mice received 1.8 mg/kg TMAO in 100µl saline, and 4 mice received 100µl saline (vehicle control). **Experiment 2** contained 20 mice, in two groups. <u>Group 1</u> contained 8 mice; 4 mice received 0.4 mg/kg GDCA in 100µl saline, and 4 mice received 100µl saline (vehicle control). <u>Group 2</u> contained 12 mice; 4 mice received 22.2 mg/kg PA in 100µl saline containing 6% DMSO, 4 mice received 1.5 mg/kg IAA in 100µl saline containing 6% DMSO, and 4 mice received 100µl saline also containing 6% DMSO (vehicle control). ***Note:*** unlike TMAO and GDCA, PA and IAA are only soluble in organic solvents such as DMSO, hence the use of DMSO above. Metabolites were stored at −20°C (IAA and TMAO) or room temperature (PA and GDCA) and dissolved immediately before use.

For sacrifice, mice were anesthetized using 2.5% Avertin (v/v). As per the original TMAO study^62^, mice were transcardially perfused after sacrifice and prior to brain removal, in order to remove contaminating blood. Specifically, mice were perfused with cold PBS (volume 40-50mL, rate 5mL/min) until clear saline exited the body and the brain and peripheral tissues appeared pale. Forceps used for brain extraction were pre-treated with RNaseZap (Thermo Fisher, Catalog #AM9780) and subsequently rinsed twice with RNase-free water to minimize the risk of RNA degradation. Brains were sagittally hemisected at the corpus callosum prior to storage. Left hemispheres were stored in 1ml RNAlater overnight at 4°C, and then dabbed on a Kimwipe to remove excess RNAlater before transfer to a fresh tube for storage at −80°C. Right hemispheres were either processed in the same manner or flash-frozen directly at −80°C. Specifically, for the TMAO-treated and corresponding vehicle control brains - which were part of the first experiment performed - the right hemispheres were processed identically to the left. In the second experiment, however, the right hemispheres were flash-frozen at −80°C without RNAlater incubation.

RNA was extracted from frozen left hemispheres using a modified version of the protocol from the QIAWave RNA Mini Kit (Qiagen, Catalog #74536), as follows. A bead beater (Qiagen Tissue Lyser II) was used to homogenize the left hemisphere in a collection microtube (Qiagen Catalog #19560) containing 700μL of pre-chilled QIAzol (Qiagen, Catalog #79306) and stainless 5mm steel bead (Qiagen Catalog #69989). For bead beating, the tube was positioned in between cold metal blocks prechilled at −20°C. Samples were homogenized at 30Hz for five minutes, kept on ice for 1 minute, then homogenized at 30Hz for an additional 2 minutes. Following homogenization, the tube was incubated at room temperature for 3-5 minutes. 200µL was then set aside for RNA extraction in a DNA LoBind tube; the remaining volume was stored in a fresh DNA LoBind tube at −80°C. 500µL of additional QIAzol was added to the 200µL tube to make 700µL; then, all the liquid was transferred to a DNA LoBind tube containing 200µL chloroform. The tube was capped and vortexed for 15 seconds, then incubated at room temperature for 2-3 minutes and centrifuged at 12,000xg for 15min at 4°C. 150µL x 2 times (300µL total) of aqueous phase was then transferred to a DNA LoBind tube containing 300µL 70% ethanol. The tube was then inverted 10 times to mix, and briefly centrifuged. ∼600µl of sample was transferred to the RNeasy Mini spin column, then spun at 10,000xg for 30 seconds at room temperature. Flow-through was discarded, and 350µL RW1 was added to the column. The column was spun at 10,000xg for 30 seconds at room temperature, and the flow-through was again discarded. 500µL RPE was added to the column, and the column was spun at 10,000xg for 30 seconds at room temperature, after which flow-through was discarded. The preceding RPE wash and spin was then repeated for 2 minutes. The tube was then centrifuged at maximum speed for one minute at room temperature. The column was transferred to a new 1.5ml DNA LoBind tube, to which 50µL RNAse-free water was added. The lid was closed and the tube was incubated at room temperature for 1-2 minutes. For elution, the tube was spun for one minute at 10,000xg at room temperature. The same 50µL of eluate was used to repeat the previous step.

RNA samples were sent to Novogene for quality control, mRNA library preparation (poly A enrichment), and sequencing (NovaSeq 6000, 150bp paired end reads, 6Gb raw data per sample). A sample list and QC statistics are included in Supplementary Table 3.

### Mouse brain transcriptomics - preliminary data analysis

The following steps were performed separately for each experiment defined above. First, FASTQ files from Novogene were trimmed to remove adapter and low-quality bases using TrimGalore v0.6.10. Trimmed paired-end reads were pseudoaligned to the standard mouse transcriptome index provided with the v1 release of kallisto^138^, using kallisto v0.50.1.

Principal component analysis (PCA) was conducted using the following functions available in DESeq2 (v1.42.1): vst (setting blind = TRUE) and plotPCA. PCA revealed two key insights that guided subsequent decisions: (1) For experiment 1 - TMAO replicate #3 was positioned away from every other TMAO-treated sample along the first two components (Fig. S6A), necessitating further investigation of this sample prior to downstream analysis. (2) For experiment 2 - PCA revealed strong separation of samples by RNA extraction batch (Fig. S6B), necessitating the inclusion of this variable as a covariate in downstream differential expression analysis.

TMAO replicate #3 was ultimately excluded from all subsequent analyses, including DESeq2 differential expression analysis, based on three additional facts beyond its placement in PCA. First, TMAO replicate #3 exhibited high leverage in the initial DESeq2 run; i.e., with TMAO replicate #3 included, it showed the highest mean Cook’s distance across tested genes, more than 1.5x that of any other sample, indicating a disproportionate influence on the linear models (Fig. S6C). Second, DESeq2 differential gene expression analysis comparing TMAO replicate #3 vs. all other samples in the experiment revealed enrichment of blood markers (*Hba-a1*: log₂FC = 2.28, padj = 3.9 × 10E-4; *Hbb-bt*: log₂FC = 1.72, padj = 0.03; Supplementary Table 7 and Fig. S6D), suggesting potential incomplete perfusion and/or other blood contamination. Third, excluding TMAO replicate #3 brought our differential expression results into closer agreement with the broader literature on TMAO, strengthening confidence in the results.

DESeq2^139^ v1.42.1 was used to determine differentially expressed genes between metabolite-treated samples and matched vehicle controls, using the default significance cutoff of alpha=0.10. For experiment 1, TMAO replicate #3 was excluded for the reasons detailed above, yielding *n =* 3 TMAO brains and *n =* 4 control brains for the final differential expression analysis. For experiment 2, all *n =* 4 brains per condition were included in the final analysis, and the RNA extraction batch was included as a covariate, based on the PCA results described above (Fig. S6B). As recommended by the DESeq2 developers, the function lfcShrink^69^ was applied to conservatively shrink log2 fold changes, using the shrinkage estimator ‘apeglm.’ All final DESeq2 differential gene expression results for PA, IAA, TMAO and GDCA are contained in Supplementary Table 7. Genes with an absolute shrunken log2 fold change ≤ 0.1 were not counted as DEGs, even if they met the significance threshold. All per-gene abundances for DESeq2 analysis were determined from the per-transcript kallisto outputs using tximport (v1.30.0).

### Mouse brain transcriptomics - gene set enrichment analysis

Gene set enrichment analysis (GSEA) was conducted using the R package fgsea (v1.28.0). The relevant gene sets (hallmark gene sets and REACTOME pathways for mouse) were downloaded from https://www.gsea-msigdb.org/gsea/msigdb/mouse/collections.jsp on July 4, 2025. Enrichments with padj ≤ 0.05 were considered statistically significant.

### Mouse brain transcriptomics - marker gene analysis

Marker genes for six different brain cell classes were determined using the following command in the R package Seurat (v5.3.0): FindAllMarkers(seurat_object, group.by = "ident", layer = “data”, only.pos = TRUE, min.pct = 0.25, logfc.threshold = 0.5). The resulting marker genes were further filtered to those with adjusted *p*-value ≤ 0.05; the filtered markers are listed in Supplementary Table 9. The Seurat object was constructed using single cell data from the mouse brain, specifically data from the 8 young mice (approximately age-matched to those in our study) included in Ximerakis *et al.*, 2019^73^. Expression data, and metadata specifying the class of each cell, were downloaded from https://singlecell.broadinstitute.org/single_cell/study/SCP263/aging-mouse-brain on August 22, 2025.

### Mouse brain transcriptomics - differential splicing analysis

Differential splicing events were identified using rMATS-turbo^80^ (v4.1.1), as follows. First, STAR (v2.7.11b) was used to align trimmed paired-end reads to the mouse genome downloaded from GENCODE (Release M37, i.e., GRCm39).

The resulting bam files were then run through rMATS-turbo with the following key parameters: -t paired --libType fr-unstranded --readLength 150 --variable-read-length --nthread 10 --cstat 0.1 --novelSS --allow-clipping. Based on the “JCEC” output files, this command initially nominated 126 differential splicing events between TMAO-treated and saline-treated brains (FDR ≤ 0.05), which we further narrowed to 44 using the following four pre-specified filters: **(1)** ≥ 5 read counts on average per group, **(2)** exclusion of events with average PSI < 0.05 or > 0.95 in both groups, **(3)** FDR ≤ 0.01, and **(4)** ΔPSI ≥ 0.10.

All sashimi plots were generated using ggsashimi^145^ (v1.1.5).

### RT-PCR validation of differential splicing

An intron retention event in *Rsrp1*, originally identified by rMATS-turbo, was validated by RT-PCR (Figs. S9 and S17) using excess RNA that had not been used for the sequencing experiment. 1.1μg RNA per brain was reverse-transcribed using the Verso cDNA Synthesis Kit (Thermo Fisher, Catalog #AB1453B). The resulting cDNA was subsequently diluted 1:4. PCR primers were designed to bind within the exons flanking the intron of interest, with the downstream primer also crossing into the subsequent exon to prevent amplification of gDNA. Primer sequences were 5’-ACGCAGCAAAAGCTCTGGGA-3’ (upstream) and 5’-GCTTTGCTACGGAATTATTGGAGCT-3’ (downstream). GoTaq Green Master Mix (Promega, Catalog #M7122) was used for PCR. The reaction mix contained 12.5μL Master Mix, 1μL of each 10 μM primer, 1μL diluted cDNA, and 9.5μL nuclease-free water. The PCR program was as follows: (1) initial denaturation at 95°C for 2min, (2) 30 cycles of 95°C for 30s, 67°C for 30s, and 72°C for 95s, and (3) final extension at 72°C for 5min. PCR products were run on a 2% agarose gel with the TriDye 1kb Plus DNA Ladder (NEB, Catalog #N3270S). The gel was imaged using an iBright™ CL750 Imaging System (Invitrogen).

### SH-SY5Y splicing experiment

SH-SY5Y cells at passage 5 were seeded in 12-well plates at a density of 0.3 × 10⁶ cells per well. The next day, cells were treated with 30μM TMAO (Cayman Chemical, Cat #17354) for 3 hours, 1% H₂O for 3 hours (vehicle control for TMAO), 100 nM pladienolide B (Tocris, Cat #6070) for 4 hours, or 0.01% DMSO for 4 hours (vehicle control for pladienolide B). After treatment, cells were harvested in TRIzol and stored at −20°C before RNA extraction, which was performed as described for the *in vitro* transcriptomic screen above. RNA samples were sent to Novogene for quality control, mRNA library preparation (poly A enrichment), and sequencing (NovaSeq X Plus, 150bp paired end reads, 15Gb raw data per sample). Read trimming and differential splicing analyses were performed as described for the mouse brain transcriptomics experiment, except using the human genome annotation, GENCODE human v47 (GRCh38).

### Bulk ATAC-seq of brains of GDCA-exposed vs. vehicle control mice

As described above under “Mouse brain transcriptomics after metabolite injection,” only the left hemisphere of each brain in these experiments underwent RNA-sequencing. The right hemispheres from GDCA-exposed and corresponding vehicle control mice were flash-frozen at - 80°C, enabling ATAC-seq.

Mouse brain nuclei were isolated following procedures established for processing mammalian brains^146^ and then ATAC-seq was carried out as previously described^147^ with some modifications.

Briefly, dissected brain tissue was added to 1 mL nuclei extraction buffer (NEB: 250 mM Sucrose, 65 mM B-glycerol, 1x protease inhibitor, 25 mM KCl, 5 mM MgCl2, 20mM HEPES-KOH, 0.5% IGEPAL CA-630, 1 mM DTT, 0.2 mM Spermine, 0.5 mM Spermidine, 2-5% NGSerum) and incubated on ice for 10-15 minutes, then dounced 20 times with pestle A and another 20 times with pestle B. Another 300 µL of NEB was added and the mixture was transferred to a 2mL tube and incubated on a rotator at 4°C for 5 minutes, then filtered through a 70µm Flowmi strainer (Sigma, Catalog #BAH136800070) into a fresh 2mL tube. The volume was adjusted to 1,200 µL with more NEB if necessary and nuclei were lightly crosslinked by adding 6.52 µl of 37% FA, incubating for 4 minutes at room temperature, and quenching with 133 µL 2.5 M Glycine solution.

Nuclei were finally isolated using density gradient separation, by adding 1,100 µL of O-50% solution (generated by mixing 18,751 µL O-60%/OptiPrep/iodaxinol solution, 3242 µL H20, 450 µL NGS, and 22.5 µL 1M DTT) in a 5mL Eppendorf tube, then creating a density gradient by adding 1,000 µL O-44% (8,800 µL O-50% plus 1,200 µL H_2_O) to the very bottom of the tube and 500 µL O-22% (2,200 µL O-50% plus 2,800 µL NEB buffer) in between the O-44% and the top layer with the nuclei. Tubes were then centrifuged for 30 minutes at 4°C with brakes off (deceleration 0, acceleration 5), and nuclei were extracted from the middle layer in the density gradient.

After nuclei isolation, nuclei were counted and approximately 100,000 were used as input into each transposition reaction in four technical replicates for each individual mouse brain. Transposition was carried out in 50 µL transposition mix, consisting of 25 µL 2x TD buffer (20 mM Tris-HCl pH 7.6, 10 mM MgCl2, 20% Dimethyl Formamide), 22.5 µL nuclease-free H_2_O, and 2.5 µL loaded Tn5 transposase (produced and assembled following previously established protocols^148^). Transposition reactions were incubated in an Eppendorf ThermoMixer at 37°C at 1,000 rpm for 30 minutes. The transposition reactions were then stopped immediately by adding 150 µL reverse crosslinking buffer (1% SDS, 0.1M NaHCO3), and reverse crosslinking was carried out for 8 hours at 65°C.

Transposed DNA was isolated using the MinElute PCR Purification Kit (Qiagen, Catalog #28004) by adding 600 µL PB Buffer and eluting in 20 µL EB Buffer heated to 65°C, then PCR amplified by mixing 20 µL transposed DNA, 2.5 µL i5 and 2.5 µL i7 PCR primers, and 25 µL 2x NEBNext High-Fidelity PCR Master Mix (New England Biolabs, Catalog #M0541L), with the following settings: 72°C for 5 minutes, 98°C for 30 seconds, followed by 10 cycles of 98°C for 10 seconds, 63°C for 30 seconds, and 72°C for 45 seconds.

Final libraries were purified using the MinElute PCR Purification Kit (Qiagen, Catalog #28004), QC-ed using a Qubit 1X dsDNA kit (Thermo Fisher Scientific, Catalog #Q33230) and an Agilent TapeStation (High Sensitivity D1000 Screen Tape® Assay), and sequenced on a NextSeq 550 instrument in a 2×38 format.

### ATAC-seq - data analysis

ATAC-seq data analysis was conducted largely as previously described^149^, with brief specifics provided below.

Two replicates (independent sets of nuclei) were sequenced per brain sample, yielding *n =* 16 total replicates from *n =* 8 mice, of which *n =* 4 were GDCA-exposed mice and *n =* 4 were the corresponding vehicle control mice (Supplementary Table 3).

Forward and reverse reads from each replicate were first trimmed to a length of 36bp using the script PEFastqToTabDelimited.py available at the following Github repository: https://github.com/georgimarinov/GeorgiScripts. Trimmed reads were aligned to both the entire mouse genome (mm10) and to the mitochondrial portion alone (chrM) using bowtie (v1.0.1). The following parameters were used for alignment to the mitochondrial genome: -p 20 -v 2 -a -t –best --strata -q -X 1000 --sam --12. The following parameters were used for alignment to the entire genome: -p 20 -v 2 -k 2 -m 1 -t --best --strata -q -X 1000 --sam --12.

The mitochondrial alignments were used to calculate the fraction of mitochondrial reads in each replicate (reported in Supplementary Table 3). For all subsequent analyses, the entire-genome alignments were used, after filtering out reads mapping to chrM.

The remaining, non-mitochondrial reads were deduplicated using the MarkDuplicates tool from Picard Tools (v1.99) using the following parameters: VALIDATION_STRINGENCY=LENIENT ASSUME_SORTED=true REMOVE_DUPLICATES=true. The resulting deduplicated .bam files were indexed using samtools v0.1.18 and converted to .wig using the script makewigglefromBAM-NH.py from the Github repository linked above. Reads per million (RPM) was used as the normalization method. The .wig files were converted to .bigWig using wigToBigWig from UCSC Utils (version from 2017-07-13). TSS enrichment scores reported in Supplementary Table 3 were calculated using the scripts signalAroundCoordinate-BW.py and ATACTSSscore.py, from the Github repository linked above. TSS coordinates were based on the UCSC mm10 refFlat gene annotation table (dated 2014-10-26).

Peaks were called for each replicate using the callpeak function from MACS2 (v2.1.1) using the following parameters: -g mm -f BAMPE --to-large -p 1e-1 --keep-dup all --nomodel. Then, reproducible peaks between replicates were identified using the previously described irreproducible discovery rate (IDR) framework^149^. IDR v2.0.4.2 was used for this step. The required script BAMPseudoReps.py from the Github repository linked above was converted to Python 3 using the 2to3 Python program prior to use.

The reproducible peaks between replicates per sample were then merged across samples using the createIterativeOverlapPeakSet.R script available at the following Github repository - https://github.com/corceslab/ATAC_IterativeOverlapPeakMerging - described in Grandi *et al.*, *Nature Protocols* 2022^150^. Blacklist regions provided to the script were downloaded from https://github.com/Boyle-Lab/Blacklist/tree/master/lists (file mm10-blacklist.v2.bed.gz) on July 11, 2025. The spm was set to 0 because confident peaks had already been identified. Additional relevant parameters included: --rule "(n+1)/2" --extend 250. This step yielded a .bed file with the coordinates of the final set of merged peaks, which was converted to SAF format and used to create a read counts table (rows = merged peaks, columns = samples) using featureCounts (v2.0.6).

Samples were retained for subsequent analysis if both replicates had TSS enrichment scores ≥ 10, the standard deemed as “acceptable” under ENCODE guidelines for mm10 (https://www.encodeproject.org/atac-seq/, last accessed July 28, 2025). As per Supplementary Table 3, 7 samples (*n* = 4 from GDCA-exposed mice, and *n* = 3 from control mice) passed this filter.

Differentially accessible peaks were determined using DESeq2 (v1.42.1) using the default significance cutoff of alpha=0.10. These peaks were further annotated using rGREAT (v2.4.0) and using the annotatePeaks.pl script from HOMER, which was installed using the configureHomer.pl script downloaded from http://homer.ucsd.edu/homer/ on July 14, 2025.

Coverage of differentially accessible peaks (Fig. S13A-B) was determined as follows. First, BAM files for individual brains within each condition were merged using samtools v1.21. The resulting two files (one for control brains and one for GDCA-exposed brains) were sorted and indexed with samtools v1.21, and subsequently converted to bigWig format using the bamCoverage function from deepTools v3.3.0, using a bin size of 100 bp and specifying counts per million (CPM) as the normalization method. Coverage matrices per condition were generated from each bigWig file using the signalAroundCoordinate-individual.py script available from https://github.com/georgimarinov/GeorgiScripts, using the previously specified parameters^149^, after conversion to Python 3 using the 2to3 Python program.

chromVAR^109^ analysis (Figs. 5F and S14) was conducted using chromVAR 1.24.0, referencing the 0.99.6 version of the JASPAR 2024 database in R version 4.3.2.

### Behavioral testing and related measurements

Behavioral testing was conducted at the Stanford Behavioral and Functional Neuroscience Laboratory. After arrival from Jackson Laboratories, mice were acclimated to a reverse 12h-12h light-dark cycle for at least one week prior to initiation of experiments, ensuring that daytime testing coincided with their naturally active (dark) phase. During this acclimation period, mice were periodically handled to facilitate habituation to the experimenter.

For experiments with **male mouse cohort 1**, mice were dosed for three consecutive days with GDCA (0.4 mg/kg) or saline vehicle (*n =* 15 mice per group, *n =* 30 total). Intravenous retroorbital injection was used as the administration route, with isoflurane as the anesthetic. The total injection volume was 50µL. Behavioral testing was conducted after the final injection (activity chamber: one day after final injection, Y-maze: two days after final injection). Mice were observed for one hour in the activity chamber and for five minutes in the Y-maze.

For experiments with **male mouse cohort 2**, mice were administered 50 mg/kg GDCA, 10 mg/kg GDCA, or saline vehicle *per os* (*n =* 15 mice per group, *n =* 45 total). Y-maze testing was conducted one hour post-dosing, across two consecutive days to accommodate all mice. Six days later, all mice were redosed (keeping the same group assignments), and core body temperature was measured using a mouse rectal probe. All *n =* 15 mice per group were measured at baseline, prior to dosing; *n =* 5 mice per group were measured at each subsequent timepoint (10min, 30min, and 60min post-dosing). Eight days later, *n =* 3 mice per group were redosed (keeping the same group assignments). Sixty minutes after dosing, mice were anesthetized with isoflurane, and plasma and brain tissues were collected for metabolomics analysis. Blood was collected by cardiac puncture, approximately 500µL per mouse, into EDTA-treated plasma collection tubes (BD Microtainer, Cat #365974). Tubes were inverted ten times, placed on wet ice, and centrifuged within 60 minutes of collection at 8,000 x g for 4 minutes at 4°C. Plasma was transferred to individual 1.5mL microcentrifuge tubes, frozen on dry ice, and stored at −80°C at the end of the day. Immediately after blood collection, mice were transcardially perfused with cold 0.9% saline. Perfusion was considered complete when clear saline exited the body and the paws and liver appeared pale. Brains were harvested, flash frozen on dry ice, and stored at −80°C at the end of the day. GDCA concentrations in plasma and brain samples were quantified by LC-MS/MS at Frontage Laboratories, Inc. in Hayward, CA. Brain tissue was homogenized in four volumes of 20% acetonitrile in water. Plasma or brain homogenate aliquots were precipitated with acetonitrile containing ammonium acetate, acetic acid, and d4-GDCA as an internal standard, then centrifuged at 3,220 x g for 15 minutes. Supernatants were analyzed using a Shimadzu LC-30AD HPLC coupled to a Sciex 6500 mass spectrometer operating in negative ionization mode with multiple reaction monitoring.

For experiments with **male mouse cohort 3**, mice were administered 50 mg/kg GDCA or saline vehicle *per os* (*n =* 15 mice in the GDCA group and *n =* 14 mice in the vehicle group; one mouse, originally intended to belong to the vehicle group, died during the acclimation/handling period). The same group assignments were maintained across all assays. Activity chamber testing was conducted one hour post-dosing. Mice were observed for one hour in the activity chamber, after which fecal pellets excreted during testing were collected from a subset of mice (*n =* 4 mice per group). The percent dry weight of fecal pellets was calculated by weighing each tube containing a pellet before and after drying on a heat block at 100°C for 23 hours. Five days after activity chamber testing, all mice were trained on a rotarod in three sessions at 16rpm for 40s, with 30min inter-session intervals. The following day, mice were dosed with GDCA or vehicle; 60 minutes after dosing, they were tested on the rotarod at 16rpm for 40s. Six days later, they were redosed and tested at the 15 minute timepoint, first at 16rpm for 40s, and then at 32rpm for 40s.

For experiments in the **female mouse cohort**, mice were administered 50 mg/kg GDCA or saline vehicle *per os*, with *n =* 15 mice per group. The same group assignments were maintained across all assays. Mice underwent activity chamber testing and fecal collection as described for male mouse cohort 3, followed one week later by rotarod testing as described for male mouse cohort 3, but with a three-day interval between the two rotarod testing days. Y-maze testing was conducted five days after the final rotarod test, as described for male mouse cohort 2. Body temperature was measured six days after Y-maze testing, as described for male mouse cohort 2, except that measurements were collected only at baseline and 60 minutes post-dosing, for all *n =* 15 mice per group. Metabolomics was performed as described for male mouse cohort 2, with final dosing and tissue collection occurring eight days after body temperature measurement, using *n =* 4 mice per group.

### Statistical analysis

Statistical tests were performed in R v4.3.2 unless otherwise stated. Statistical analyses performed outside R used the software specified in the Methods, including HOMER, g:Profiler and rMATS-turbo. Multiple testing correction was applied within each distinct family of hypotheses, using the correction method appropriate for each analysis; for example, the default Benjamini-Hochberg correction was applied within each DESeq2 result set, and Dunnett’s test was used for post hoc comparisons following each one-way ANOVA.

## Data availability

All raw sequencing data are available under NCBI BioProject ID PRJNA1328687^151^.

## Code availability

Code is described in the Methods section and available at https://github.com/meenachakra/metabolites_paper.

## Acknowledgements

We thank Jeff Rubin, Ren Song, Ayan Mondal, Hongchao Guo, Michael Bassik, Betsy Mellins, and Mark Kay for guidance surrounding brain endothelial cell culture and experiments. Jeff Rubin and Mark Kay provided us with the hCMEC/D3 cell line. We thank Aaron Gitler and Caiwei Guo for providing SH-SY5Y cells and guidance on culturing them. We also thank Jakub Rajnuk for advice during the metabolite selection process, Anshul Kundaje for guidance on RNA-seq data analysis, Garrett Patrick for reviewing the manuscript during its preparation, Ariel Gulasch for assistance with *in vitro* transcriptomic screening, and Rudy Wycallis at the Stanford Shared FACS Facility for assistance with flow cytometry. Several non-data figures were created using BioRender.com. This work was funded, in part, by NIH R01AI148623 and R01AI143757, NCI R01 CA301727, a Stand Up 2 Cancer Grant, and the Stanford Wu Tsai Neurosciences Institute. M.C. was supported by the National Defense Science and Engineering Graduate Fellowship. I.E.P. was supported by the Stanford REACH PhD fellowship. S.M.S. was supported by a Stanford Bio-X Bowes Fellowship. Computing for this project was performed, in part, on the Stanford SCG Bioinformatics Cluster (RRID:SCR_026876). The Stanford Behavioral and Functional Neuroscience Laboratory is supported by the NIH S10 Shared Instrumentation for Animal Research (1S10OD030452-01).

## Author contributions

**MC:** conceptualization, investigation, methodology, data curation, formal analysis, visualization, writing - original draft, writing - review and editing. **SMS:** investigation, methodology. **IEP:** methodology. **DJR:** investigation, methodology. **GKM:** investigation, methodology, data curation, formal analysis. **MNM:** investigation, methodology, data curation, formal analysis. **AAM:** investigation, writing - review and editing. **JLEB:** investigation, methodology. **AN:** visualization. **JWJ:** methodology. **JCW:** supervision. **SXL:** methodology, supervision. **SMD:** methodology, supervision. **WJG:** supervision. **NLS:** investigation, methodology, data curation, formal analysis. **MS:** methodology, supervision. **AB:** funding acquisition, supervision. **TWC:** supervision. **ASB:** conceptualization, methodology, funding acquisition, supervision, writing - original draft, writing - review and editing. All authors read and approved the manuscript.

## Supplementary information

Supplementary Information is available for this paper, as follows:

1. PDF of Supplementary Figures

2. Supplementary Table 1 - Properties of the 30 metabolites in the final screening panel

3. Supplementary Table 2 - Enriched KEGG pathways for genes upregulated by LPS

4. Supplementary Table 3 - Sample information for omics assays

5. Supplementary Table 4 - DESeq2 results, screen batches 1 and 2

6. Supplementary Table 5 - DESeq2 results, screen batches 3 and 4

7. Supplementary Table 6 - DEGs in AD or PD pathways for 12 metabolites

8. Supplementary Table 7 - DESeq2 results, mouse brain RNA-seq

9. Supplementary Table 8 - GSEA results, mouse brain RNA-seq

10. Supplementary Table 9 - Marker genes per cell class, Ximerakis data

11. Supplementary Table 10 - Significant differential splicing events in TMAO brains

12. Supplementary Table 11 - Significant differential splicing events in SH-SY5Y

13. Supplementary Table 12 - Studies associating GDCA in blood with AD, PD, or CI

14. Supplementary Table 13 - ATAC-seq data analysis results

15. Supplementary Table 14 - Behavioral and temperature data, and related stats

16. Supplementary Table 15 - Metabolomics after oral GDCA administration

17. Supplementary Note 1 - Images of the HMDB concentration listings that were referenced when designing the *in vitro* transcriptomic screen

