## Supplementary Figures for "Age-related microbiome metabolites alter RNA splicing and chromatin accessibility in the brain"

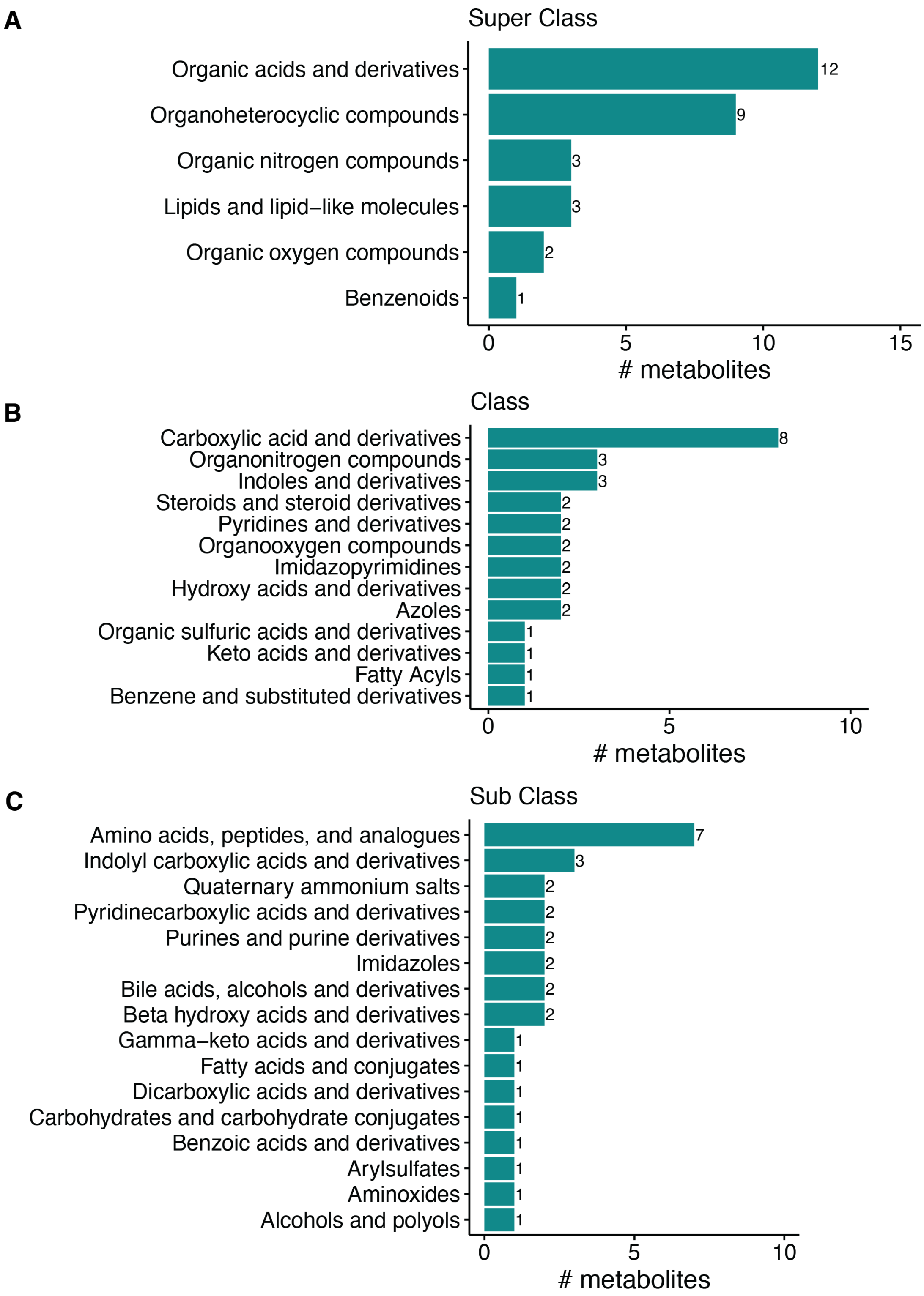


**Fig. S1.** Distribution of metabolites in the final screening panel according to the chemical taxonomy in the Human Metabolome Database (HMDB, version 5.0). **(A)** Number of metabolites per super class. **(B)** Number of metabolites per class. **(C)** Number of metabolites per sub class. Supplementary Table 1 specifies the taxonomic information for each individual metabolite.

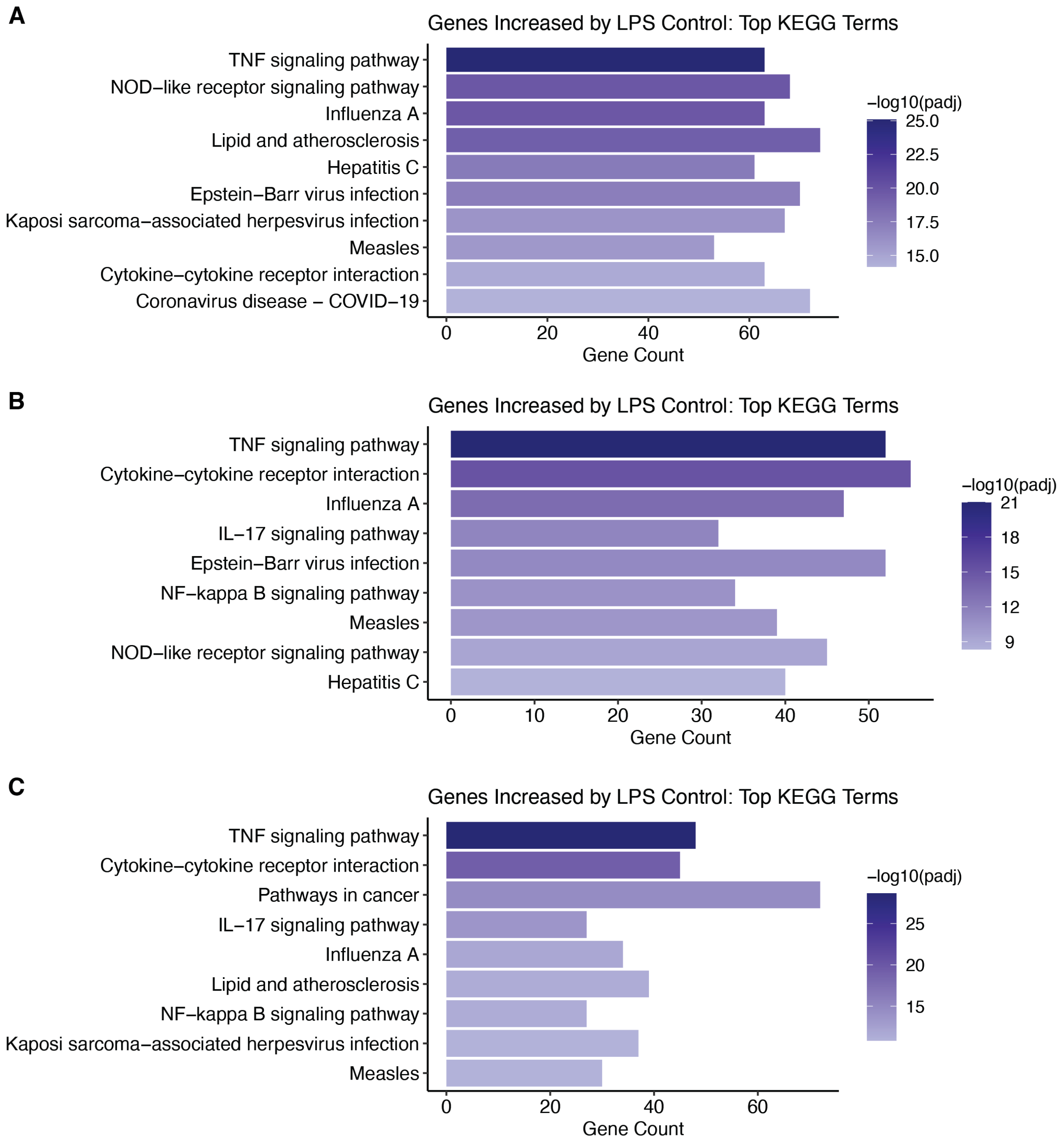


**Fig. S2.** Top enriched KEGG pathways for the genes significantly upregulated by the LPS control in sequencing batches **(A)** #2, **(B)** #3 and **(C)** #4 of the *in vitro* transcriptomic screen. The non-specific “KEGG root term” was filtered out. See Fig. 2B for the analogous graph from batch #1, and Supplementary Table 2 for exact *p*-values for all batches.

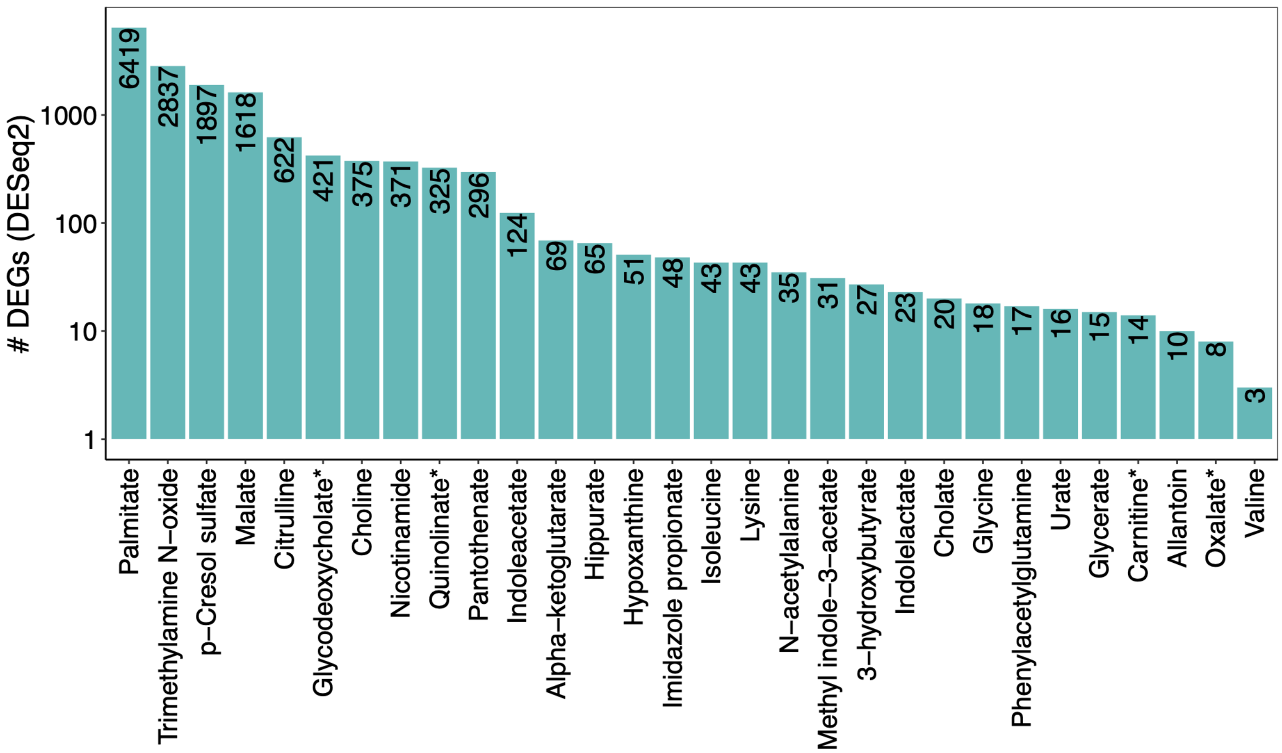


**Fig. S3.** Number of differentially expressed genes (DEGs) in hCMEC/D3 after 3h treatment with each of the 30 test metabolites. Asterisk = one of the three biological replicates for the metabolite treatment was removed as outlier, based on PCA plots and QC statistics (see Methods, Supplementary Table 3, and Figs. S4-S5).

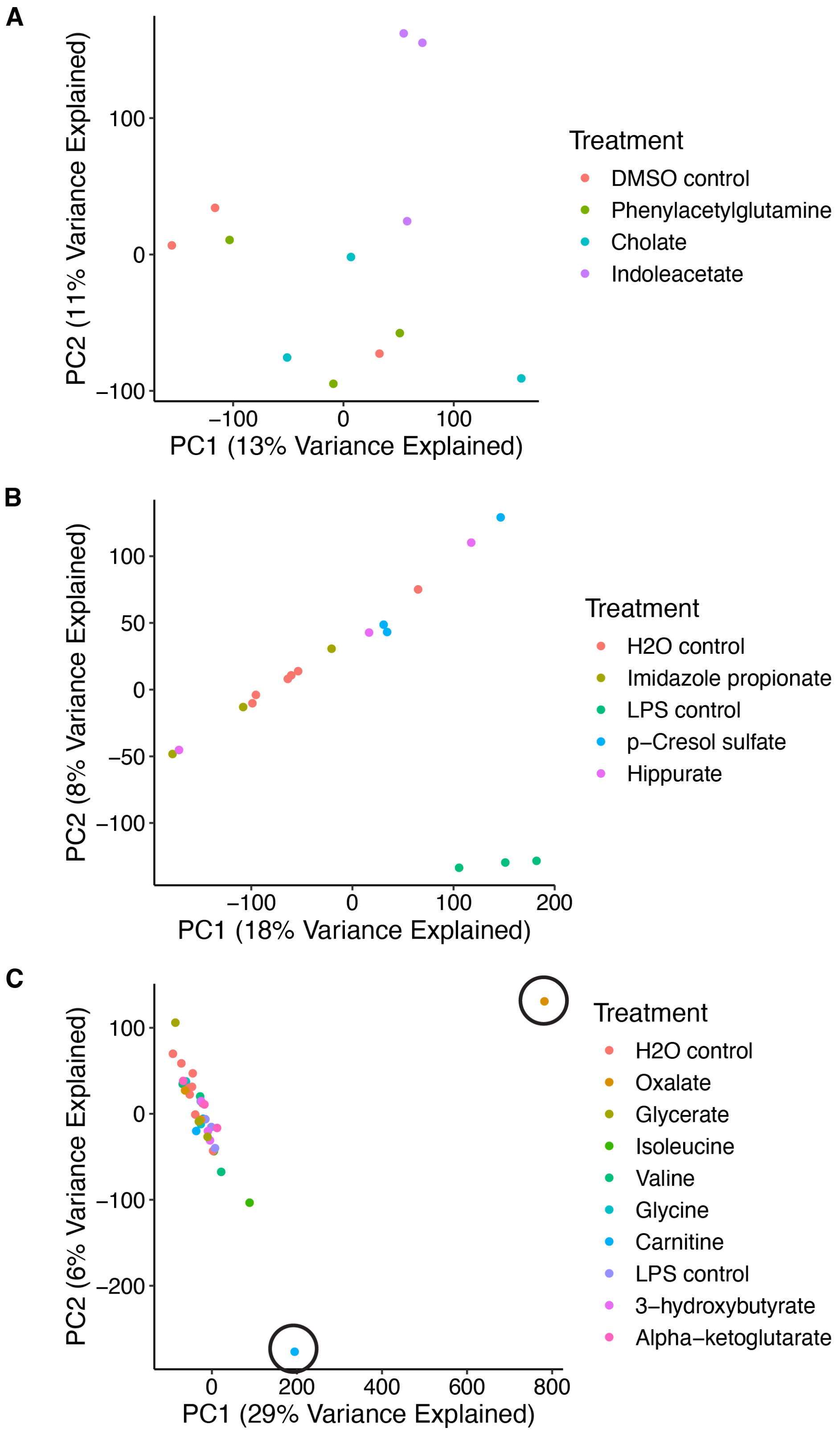


**Fig. S4.** PCA plots for RNA-seq data from the *in vitro* transcriptomic screen, batches #1 and 2, specifically: **(1)** All samples from batch 1 with 1% DMSO in the assay media, **(2)** All samples from batch 1 with 1% $H_2O$ in the assay media, **(3)** All samples from batch 2 (all of which had 1%

$H_2O$ in the assay media). Circled are the samples that were removed as outliers prior to downstream analysis, based on PCA and supporting QC statistics (see Methods and Supplementary Table 3).

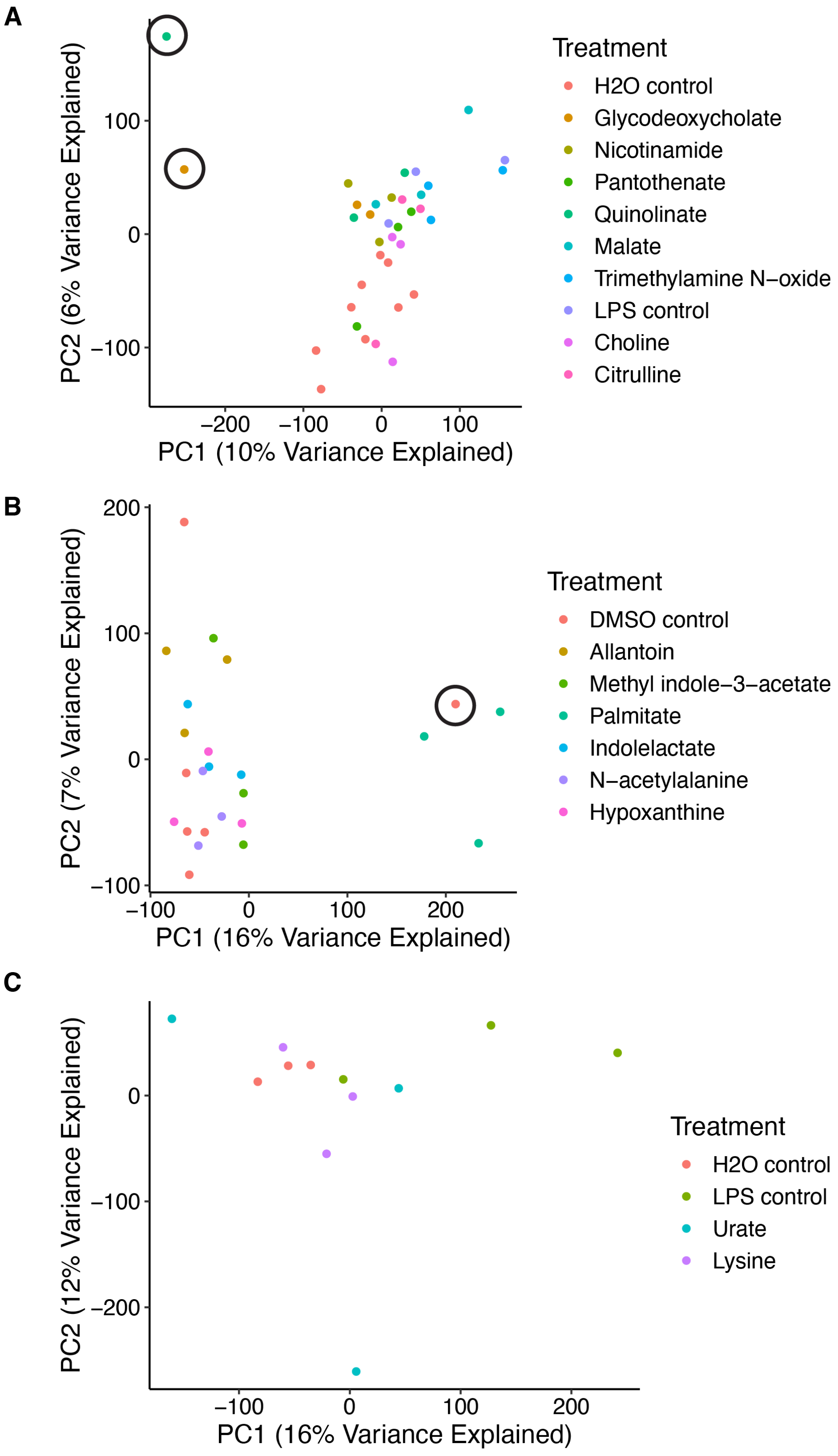


**Fig. S5.** PCA plots for RNA-seq data from the *in vitro* transcriptomic screen, batches #3 and 4, specifically: **(1)** All samples from batch 3 (all of which had 1% $H_2O$ in the assay media), **(2)** All samples from batch 4 with 1% DMSO in the assay media, **(3)** All samples from batch 4 with 1% $H_2O$ in the assay media. Circled are the samples that were removed as outliers prior to downstream analysis, based on PCA and supporting QC statistics (see Methods and Supplementary Table 3).

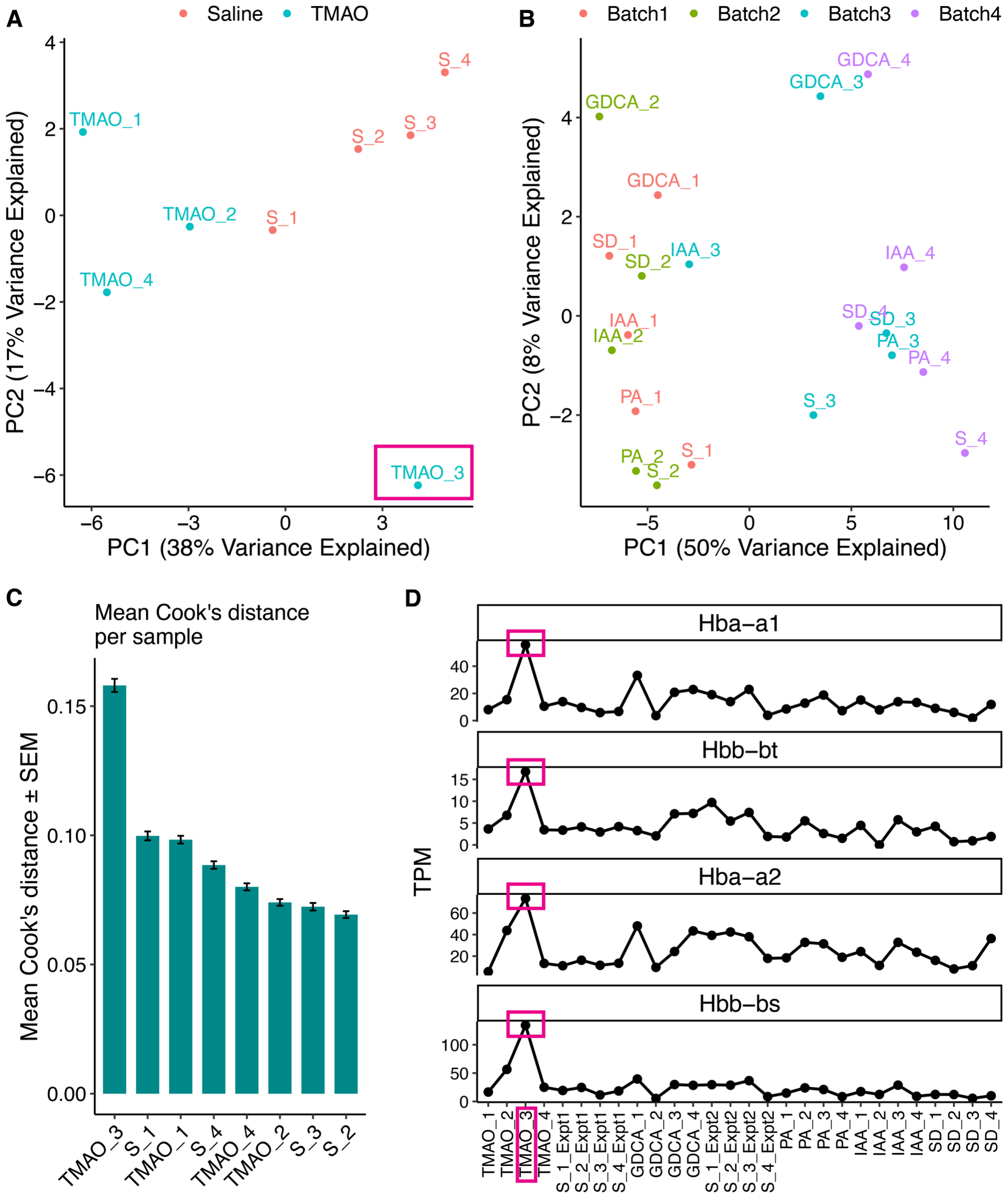


**Fig. S6.** Assessment of sample-level variation in mouse brain RNA-seq data. **(A-B)** Principal component analysis (PCA) of mouse brain RNA-seq samples from (A) Experiment 1 and (B) Experiment 2, as defined in the Methods section "Mouse brain transcriptomics after metabolite injection." PCA revealed two key features that guided downstream analyses. In Experiment 1, TMAO replicate #3 separated from the other TMAO-treated samples along the first two principal components, necessitating further investigation of this sample prior to downstream analysis (see Methods and subsequent panels). In Experiment 2, samples separated strongly by RNA extraction batch, necessitating the inclusion of batch as a covariate in downstream differential expression analysis. **(C)** In an initial differential expression analysis within Experiment 1, TMAO replicate #3 showed the highest mean Cook's distance across tested genes, more than 1.5x that of any other sample, indicating a disproportionate influence on the linear models. **(D)** Across samples from both experiments, TMAO replicate #3 showed higher levels of hemoglobin transcripts than the other samples. This observation was consistent with a differential gene expression analysis performed within Experiment 1, which identified elevated hemoglobin transcripts in TMAO replicate #3 relative to the other Experiment 1 samples, including transcripts encoding *Hba-a1* (log2FC = 2.28, padj = 3.9E-4; Supplementary Table 7) and *Hbb-bt* (log2FC = 1.72, padj = 0.03; Supplementary Table 7). As described in the Methods, TMAO replicate #3 was ultimately excluded from downstream analyses.

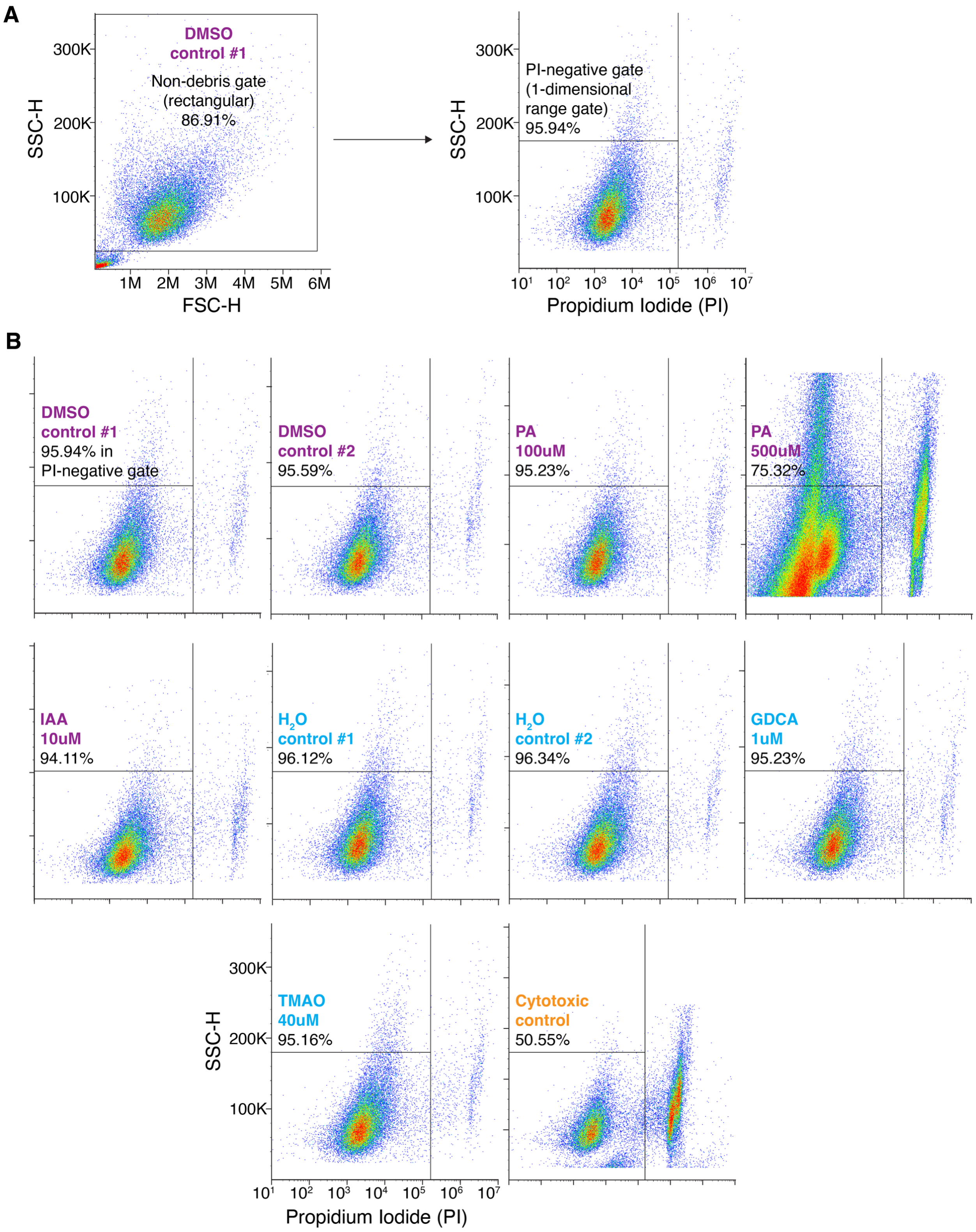


**Fig. S7.** Cytotoxicity testing in hCMEC/D3 cells following 3-hour exposure to various doses of the four metabolites tested in mice: palmitic acid (PA), indoleacetic acid (IAA), glycodeoxycholic acid (GDCA), and trimethylamine N-oxide (TMAO). Cytotoxicity was measured using the flow-cytometry-compatible dye propidium iodide (PI - Thermo Fisher, Cat #R37169), which stains dead cells. PA and IAA were dissolved in DMSO, so the cells exposed to PA and IAA were compared to cells exposed to the same concentration of DMSO in the assay media (1%). Analogously, GDCA and TMAO were dissolved in $H_2O$, so the cells exposed to these metabolites were compared to a “1% $H_2O$” vehicle control. Each plot represents one well of cells in a 12-well plate. **(A)** Gating strategy, shown for one of the DMSO control samples. First, a rectangular non-debris gate was defined, followed by a one-dimensional PI-negative gate (all cells with PI value less than ~$10^5$) representing live cells. The same gates were applied to all samples. **(B)** PI-negative gates for all samples. As shown in the bottom-left plot, the x-axis is displayed on a logarithmic scale with ticks at each power of ten, while the y-axis is linear with ticks at intervals of 100K. The “cytotoxic control” sample represents cells that were exposed to 20% DMSO in the assay media for 3 hours, based on prior literature on the cytotoxic effects of DMSO at high concentrations (PMID 29125561).

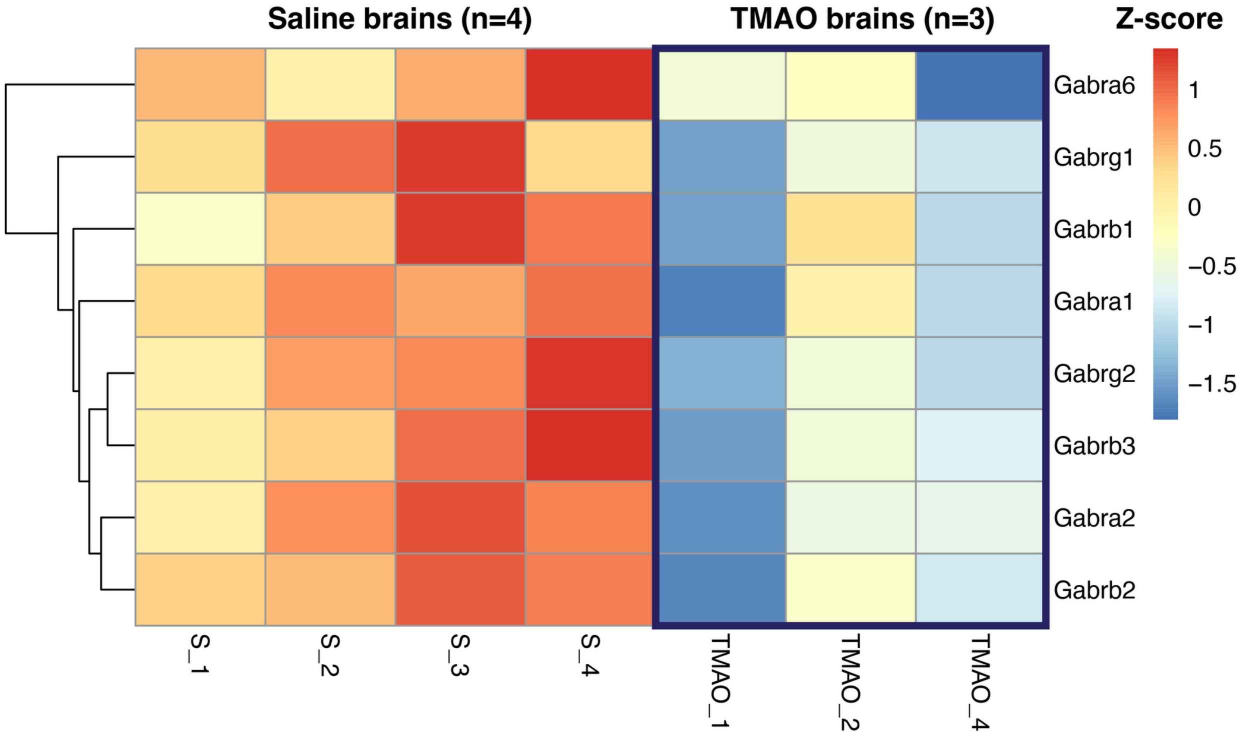


**Fig. S8.** rlog-stabilized counts, converted to Z-scores (# standard deviations away from row mean), for $GABA_A$ receptor subunits that were significantly downregulated in brains from mice exposed to TMAO, vs. brains from mice exposed to the saline vehicle control. As detailed in the Methods, one TMAO-exposed brain was excluded from analysis after QC, yielding *n* = 4 saline-exposed brains and *n* = 3 TMAO-exposed brains.

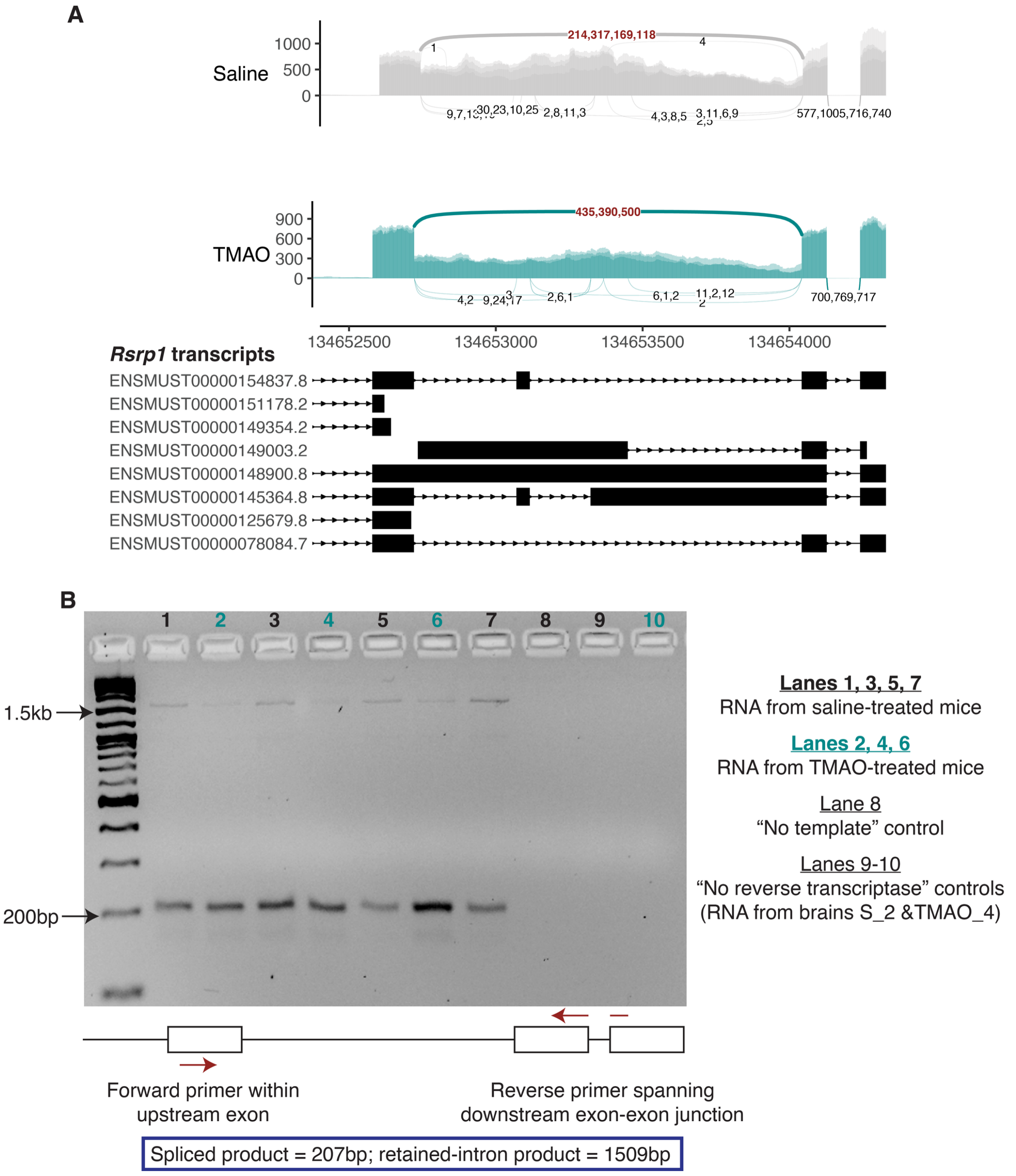


**Fig. S9.** Differential splicing of *Rsrp1* in brains from TMAO- vs. saline-treated mice (FDR = 1.96E-7 based on rMATS-turbo; see Supplementary Table 10). *n* = 3 TMAO samples and *n* = 4 saline samples were included in the analysis (see Methods). **(A)** Sashimi and coverage plot (*y*-axis = read counts) showing increased retention of the displayed intron in the saline condition. The base plot was generated using ggsashimi (PMID 30118475). **(B)** RT-PCR validation of the differential splicing event. See Fig. S17 for the unprocessed original gel image.

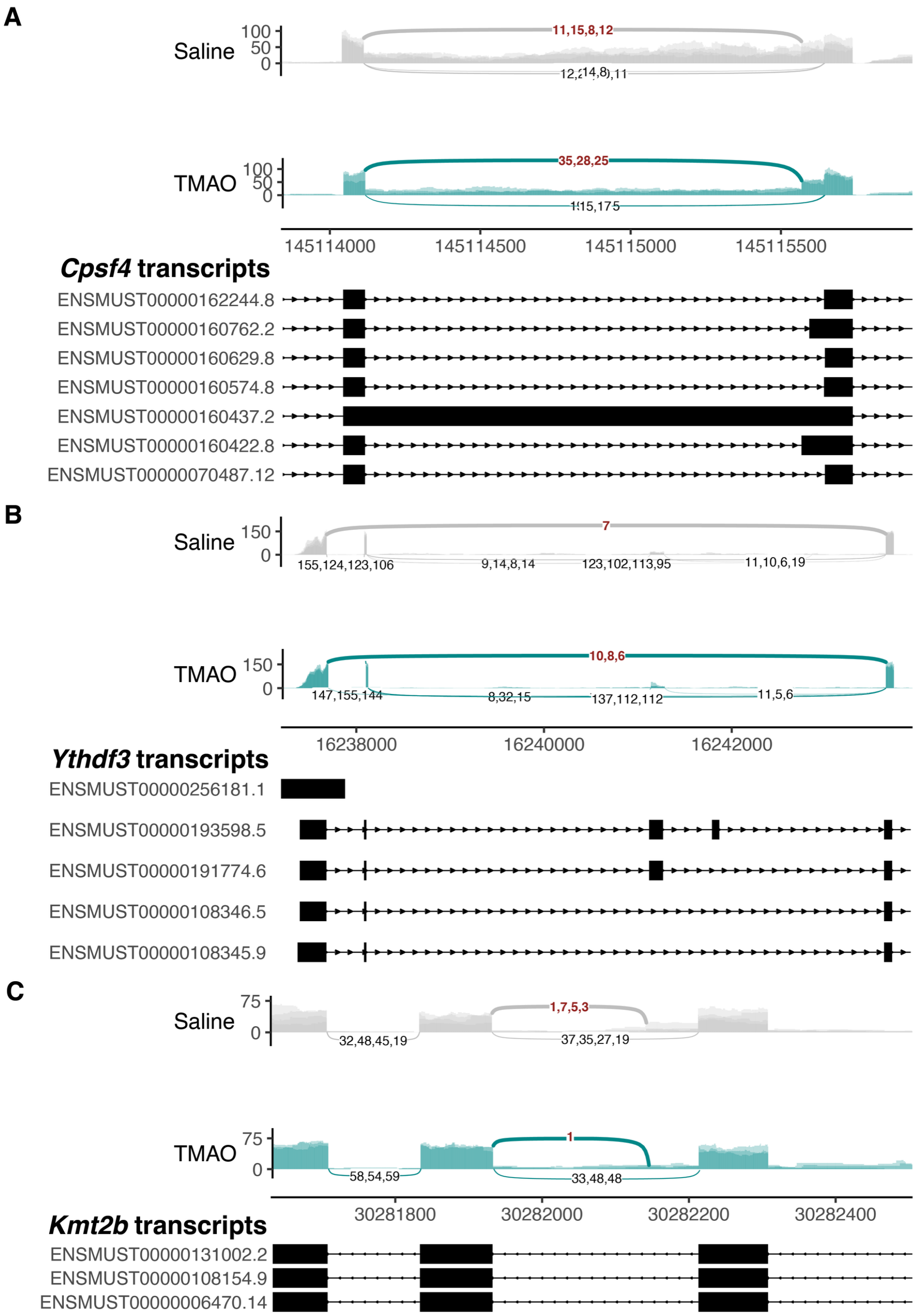


**Fig. S10.** Sashimi and coverage plots of differential splicing events in *Cpsf4*, *Ythdf3*, and *Kmt2b* in brains from TMAO- vs. saline-treated mice. The *y*-axis shows read counts. Base plots were generated using ggsashimi (PMID 30118475). Differential splicing events were identified by rMATS-turbo (PMID 38396040). *n* = 3 TMAO samples and *n* = 4 saline samples were included in the analysis (see Methods). **(A)** Saline-treated brains showed increased retention of the displayed *Cpsf4* intron relative to TMAO-treated brains; event FDR = 0.0005, see Supplementary Table 10. **(B)** TMAO-treated brains showed increased skipping of the displayed *Ythdf3* exon(s) relative to saline-treated brains; event FDR = 0.001, see Supplementary Table 10. The relevant exon-skipping junction was detected in only one of four saline-treated brains but in all three TMAO-treated brains. **(C)** rMATS-turbo identified a novel splice junction within an intron of canonical *Kmt2b* transcripts, defining a previously unannotated intron and creating an alternative 5’ splice site for the downstream exon. This splice junction was detected in one of three TMAO-treated brains, with one supporting read. By contrast, the junction was detected in all four saline-treated brains, with supporting read counts of 1, 7, 5, and 3. Event FDR = 5.46E-6, see Supplementary Table 10.

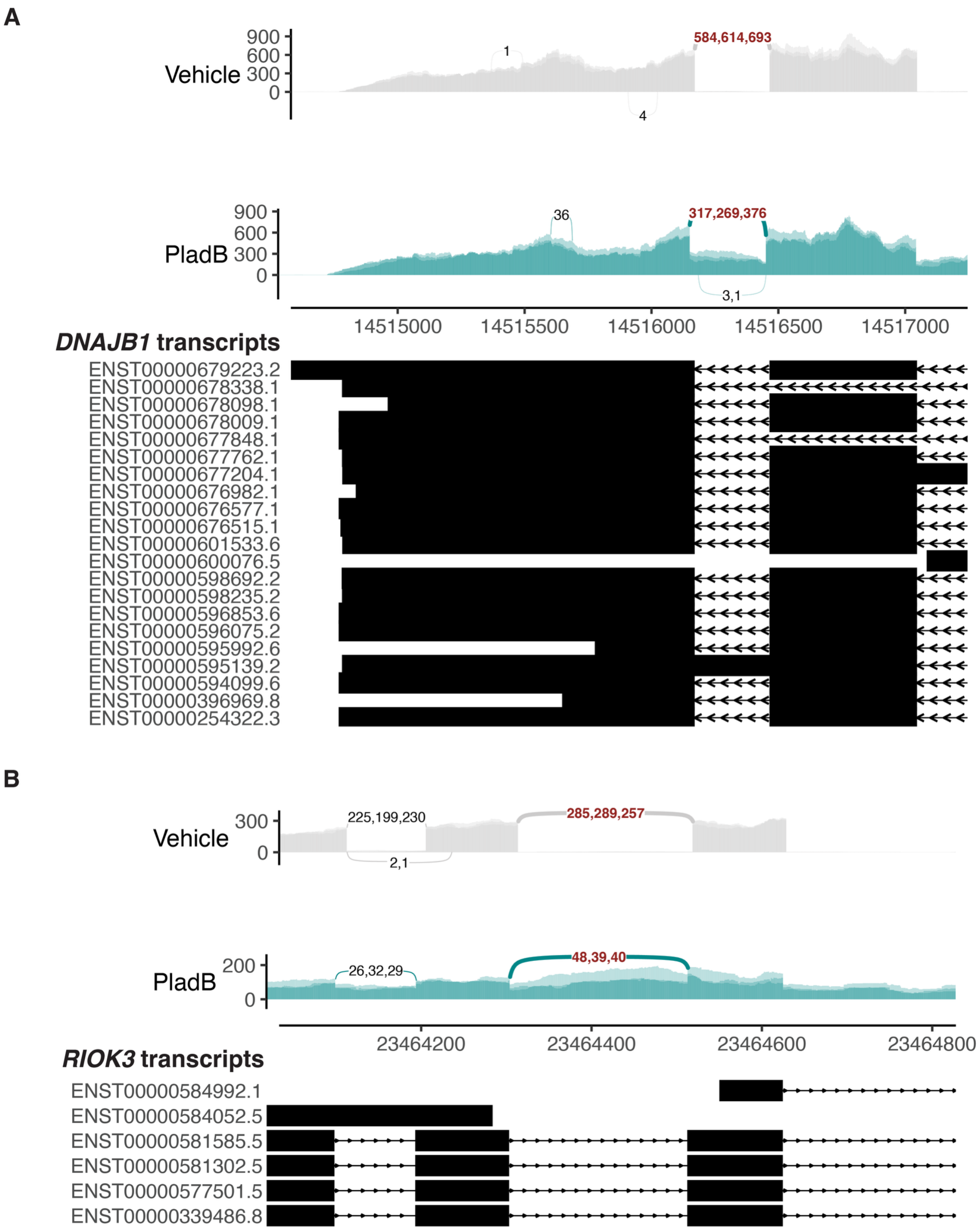


**Fig. S11.** Sashimi and coverage plots of differential splicing events in *DNAJB1* and *RIOK3* in SH-SY5Y human neuroblastoma cells treated with splicing inhibitor pladienolide B or vehicle for 4 hours (*n* = 3 replicates per condition). *Y*-axis shows read counts. Base plots were generated with ggsashimi (PMID 30118475), and differential splicing events were identified with rMATS-turbo (PMID 38396040). Pladienolide B increased retention of the introns displayed in panels **(A)** and **(B)**; both event FDRs were reported as 0 by rMATS-turbo (Supplementary Table 11).

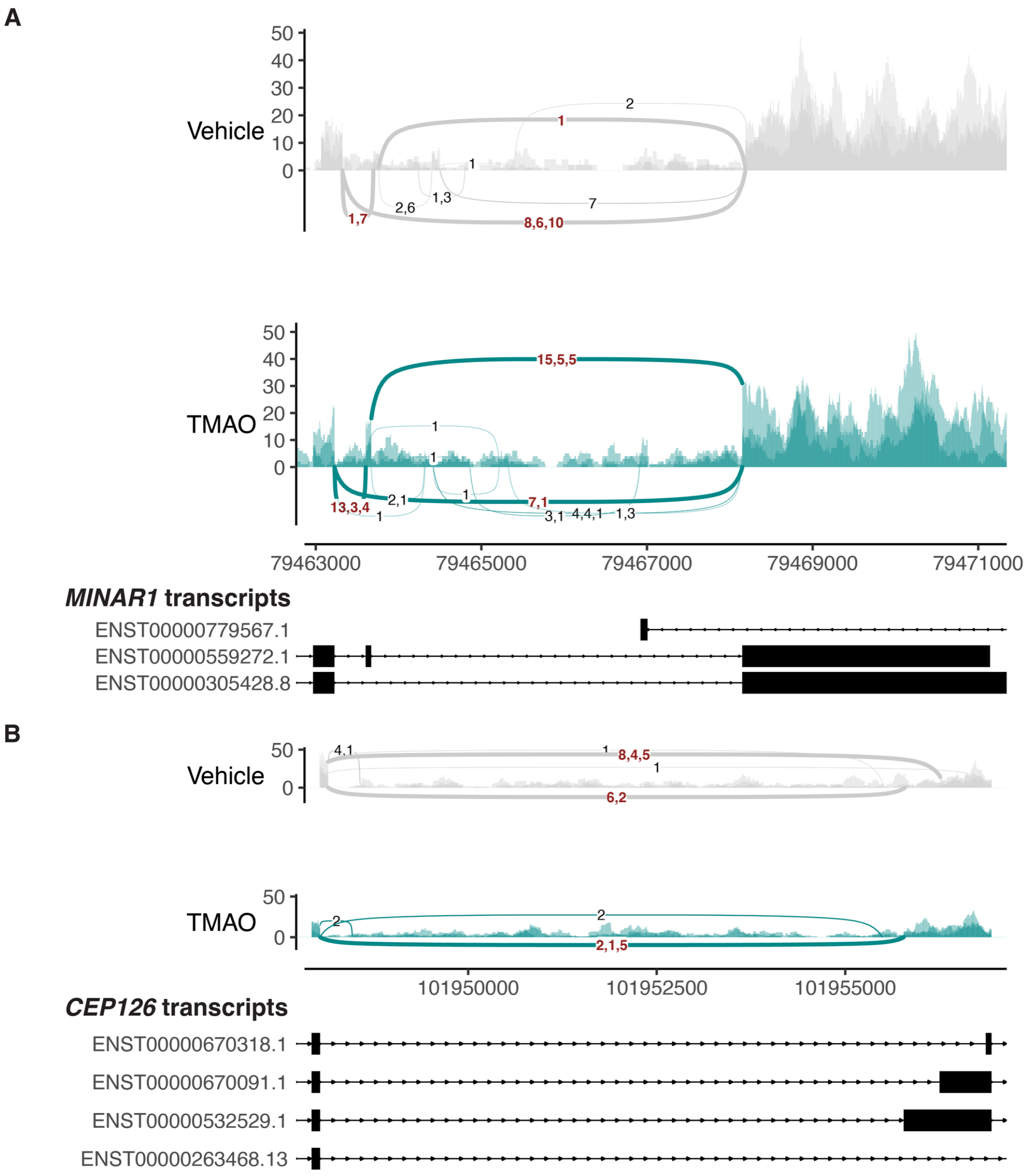


**Fig. S12.** Examples of differential splicing events in SH-SY5Y human neuroblastoma cells treated with TMAO or vehicle for 3 hours ($n$ = 3 replicates per condition). *Y*-axis shows read counts. Base plots were generated with ggsashimi (PMID 30118475), and differential splicing events were identified with rMATS-turbo (PMID 38396040). **(A)** An exon of *MINAR1* was skipped more frequently in vehicle-treated cells, vs. TMAO-treated cells (event FDR = 1E-3, Supplementary Table 11). **(B)** In *CEP126*, the junction corresponding to the alternative 3' splice site and shorter exon on the right was detected in all three vehicle-treated samples, but in none of the three TMAO-treated samples (event FDR = 1E-4; Supplementary Table 11).

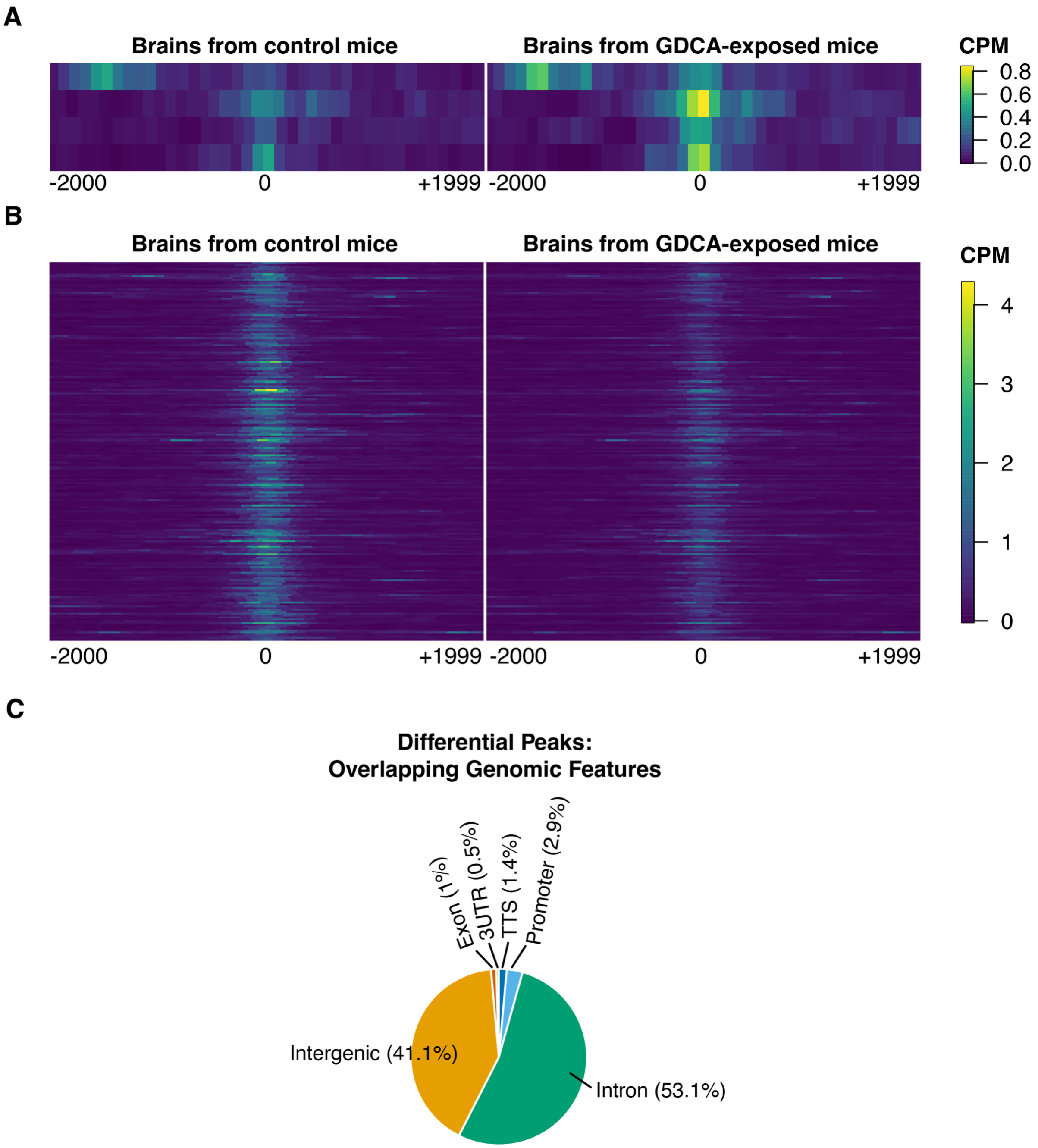


**Fig. S13.** Coverage and annotation of differentially accessible ATAC-seq peaks in brains from GDCA-exposed versus control mice. **(A)** Coverage heatmap for the 4 genomic peaks that were significantly more accessible in brains from GDCA-exposed mice, vs. control. Each row represents one peak, with 0 representing the central base of the peak. BAM files for individual brains within each condition were merged prior to visualization (Methods). **(B)** Same as (A), but for the 203 genomic peaks that were significantly less accessible in brains from GDCA-exposed mice vs. control. **(C)** Distribution of the 207 differentially accessible peaks, according to the genomic feature they overlap (intergenic region, exon, 3' UTR, transcription termination site (TTS), promoter, intron).

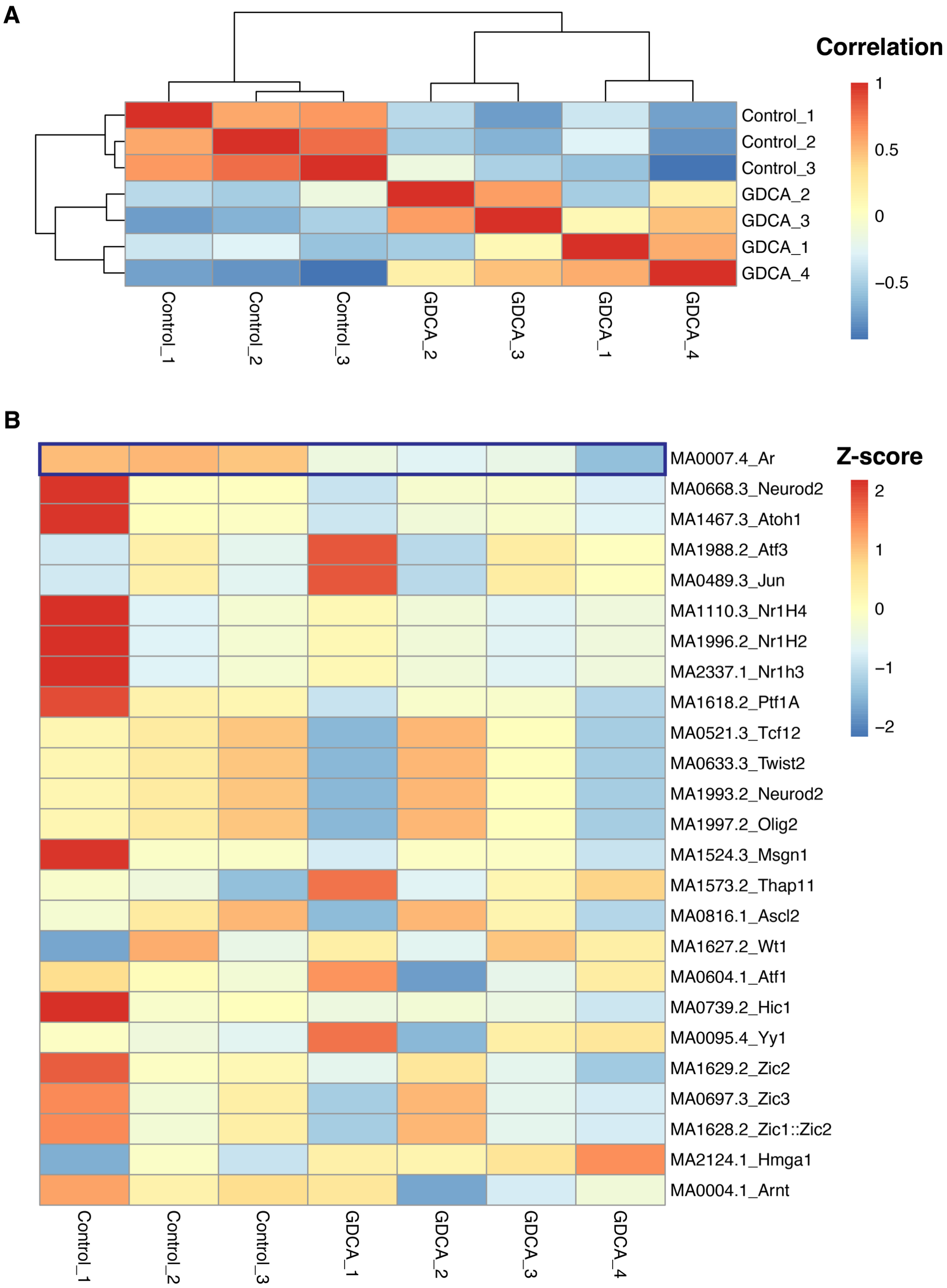


**Fig. S14.** ChromVAR analysis of GDCA ATAC-seq data quantifies transcription factor (TF) motif accessibility patterns across brain samples. **(A)** Correlation heatmap, showing the Pearson correlations between samples based on the accessibilities of the core collection of mouse TF motifs from the JASPAR 2024 database. Rows and columns are clustered by distance, i.e., by

1-correlation. Brains cluster by condition (GDCA-treated vs. control). **(B)** Heatmap of chromVAR deviation score (proxy for accessibility) for the 25 TF motifs with the greatest variability in accessibility across samples. Rows are centered and scaled (per-motif). The accessibility of *MA0007.4_Ar* was consistently decreased in brains from GDCA-exposed vs. control mice.

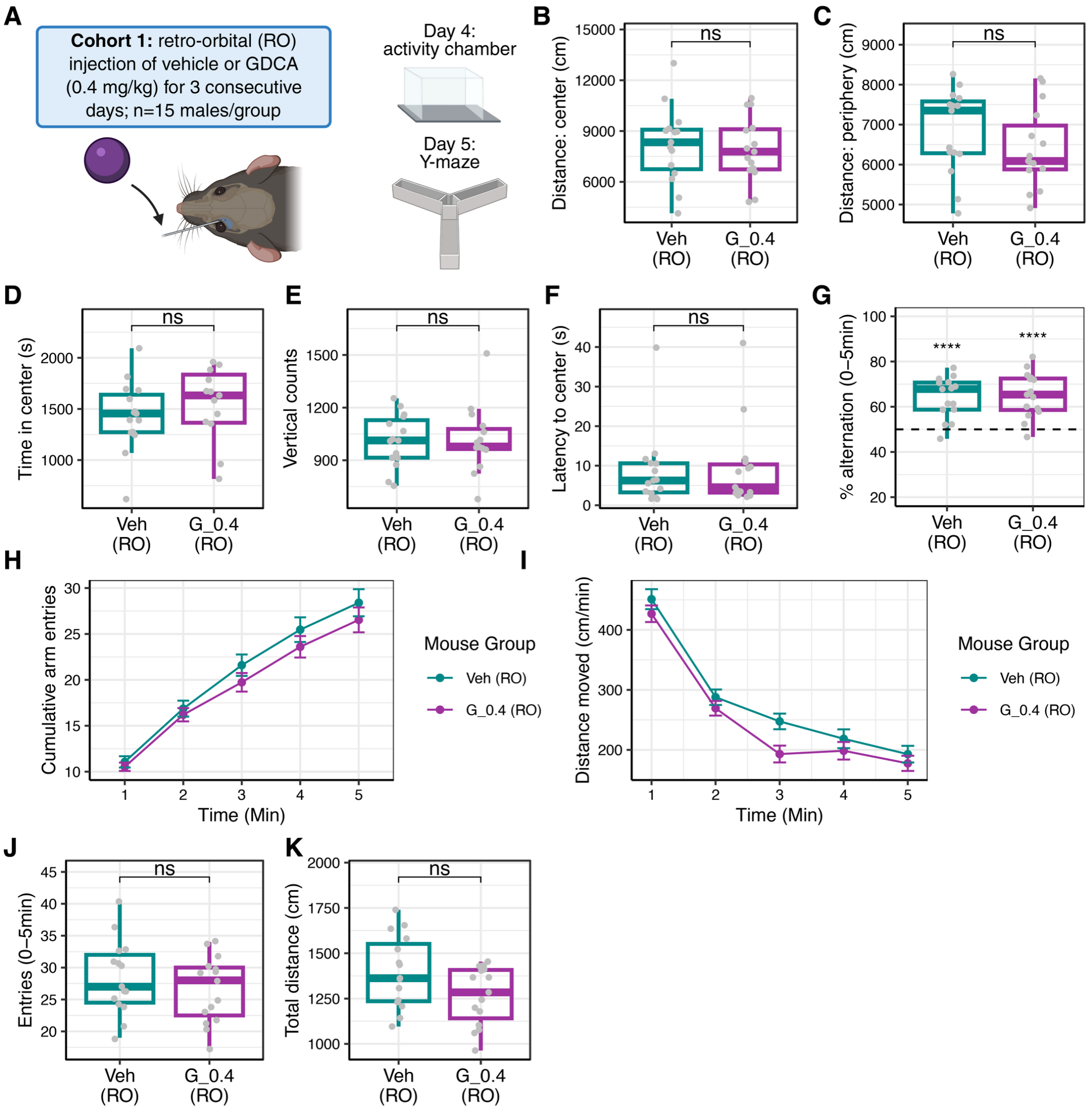


**Fig. S15.** Behavioral testing results following retro-orbital (RO) injection of vehicle or GDCA in the first behavioral cohort. Unless otherwise indicated, statistics were calculated using Welch's two-sample, two-sided *t*-test. **(A)** Schematic showing experimental design. Male mice received retro-orbital injections of vehicle or GDCA (0.4 mg/kg) once daily for 3 consecutive days, with $n$ = 15 mice per group. Activity chamber testing was performed for one hour on day 4, followed by Y-maze testing for five minutes on day 5. **(B)** Total distance traveled in the center of the activity chamber. **(C)** Total distance traveled in the periphery of the activity chamber. **(D)** Total time spent in the center of the activity chamber. **(E)** Total vertical counts in the activity chamber. **(F)** Latency to first entry into the center of the activity chamber. **(G)** Percent alternation among maze arms during the Y-maze test. An alternation was defined as entry into the novel arm after exiting a maze arm, rather than re-entry into either of the two most recently visited arms.

Percent alternation was calculated as 100 * (number of alternations) / (total number of alternation opportunities). Statistics compare alternation percentages within each group to chance level (50%) using Welch's one-sample, two-sided *t*-test. **(H)** Cumulative arm entries in the Y-maze. Lines show group means ± SEM at each time point. **(I)** Distance traveled per minute in the Y-maze, plotted as in (H). **(J)** Boxplot of total arm entries in the Y-maze. **(K)** Boxplot of total distance traveled in the Y-maze.

In all panels, ns indicates not statistically significant, * indicates $p \leq 0.05$, ** indicates $p \leq 0.01$, *** indicates $p \leq 0.001$, and **** indicates $p \leq 0.0001$. Full statistics are reported in Supplementary Table 14. In all boxplots, boxes represent the interquartile range (IQR), horizontal lines indicate medians, and whiskers extend to the most extreme values within 1.5 x IQR of the hinges.

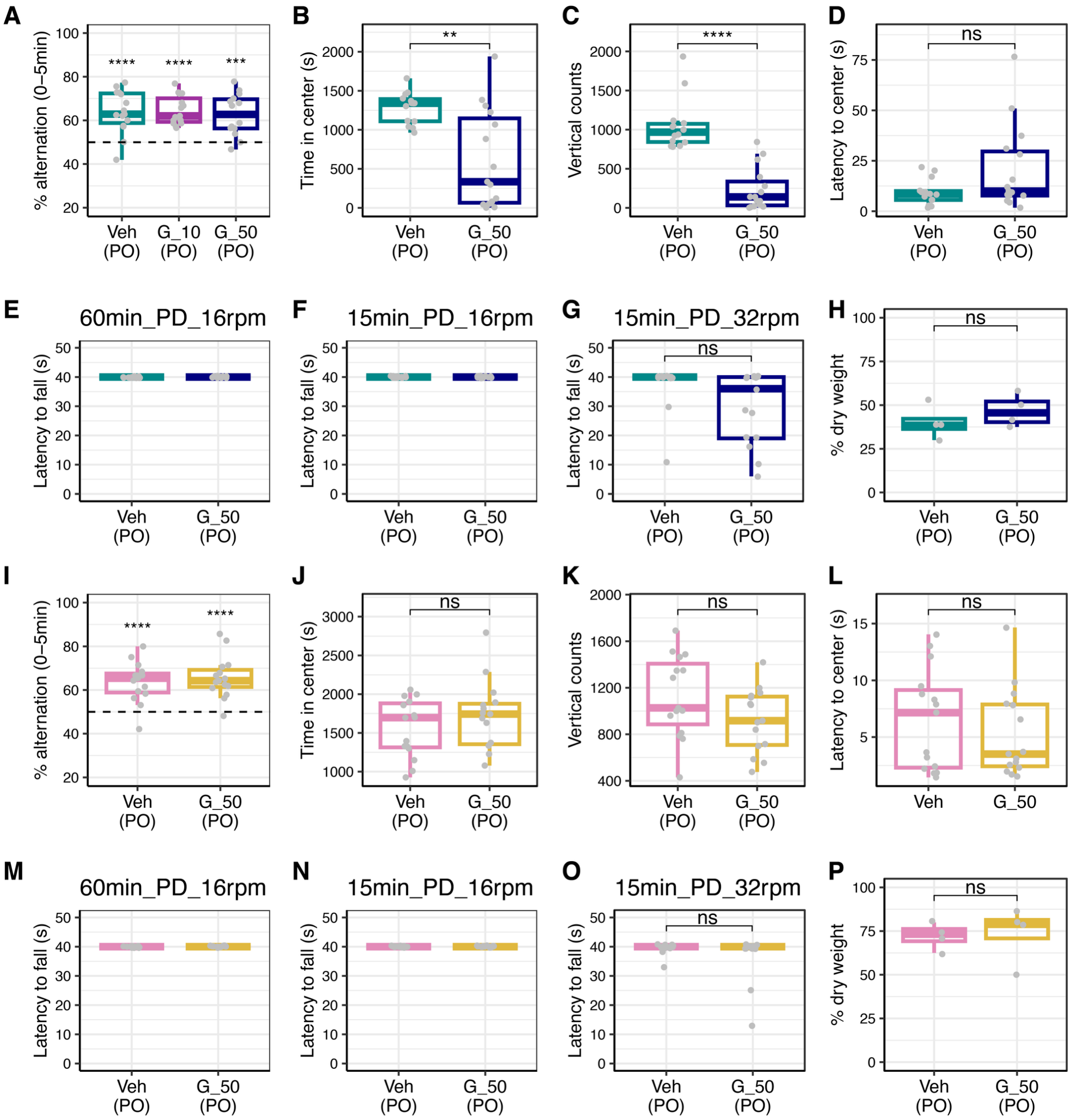


**Fig. S16.** Additional behavioral testing results from the oral gavage experiments introduced in Fig. 6. Panels (A)-(H) show experiments in male mice, and panels (I)-(P) show analogous experiments in female mice. In all panels, Veh (PO), G_10 (PO), and G_50 (PO) indicate oral gavage with vehicle, 10 mg/kg GDCA, and 50 mg/kg GDCA, respectively. PO = *per os*. **(A)** Percent alternation among maze arms in a Y-maze test conducted one hour after gavage. An alternation was defined as entry into the novel arm after exiting a maze arm, rather than re-entry into either of the two most recently visited arms. Percent alternation was calculated as 100 * (number of alternations) / (total number of alternation opportunities). Statistics compare alternation percentages within each group to chance level (50%) using Welch's one-sample, two-sided *t*-test. There were *n* = 15 mice per group; however, one mouse from the G_50 group was excluded from the percent alternation statistics because it had fewer than three arm entries and therefore no opportunities to alternate. **(B)** Total time spent in the center of an activity chamber over one hour, beginning one hour after gavage ($n$ = 14 mice in the vehicle group, $n$ = 15 mice in the GDCA group; see Methods). **(C)** Total vertical counts in the activity chamber. **(D)** Latency to first entry into the center of the activity chamber. **(E)** Latency to fall from a rotarod rotating at 16rpm. Testing was conducted 60min post-dosing and lasted 40s. No mice fell from the rod; therefore, latency to fall was 40s for all mice. There were $n$ = 14 mice in the vehicle group and $n$ = 15 mice in the GDCA group; see Methods. **(F)** Same as (E), except testing was conducted 15min post-dosing on a different day. **(G)** Same as (F), except the rotation speed was 32rpm. Testing for each mouse was conducted immediately after the test shown in (F). Groups were compared using Welch's two-sample, two-sided $t$-test ($p$ = 0.057). Fisher's exact test comparing the proportion of mice that fell from the rod in each group yielded $p$ = 0.05017; see Supplementary Table 14. **(H)** Percent dry weight of fecal pellets collected from $n$ = 4 mice per group during the one-hour activity chamber test. Groups were compared using Welch's two-sample, two-sided $t$-test. **(I)-(P)** Analogous plots for female mice. There were $n$ = 15 mice per group for all female experiments.

In all panels, ns indicates not statistically significant, * indicates $p \leq 0.05$, ** indicates $p \leq 0.01$, *** indicates $p \leq 0.001$, and **** indicates $p \leq 0.0001$. Full statistics are reported in Supplementary Table 14. In all boxplots, boxes represent the interquartile range (IQR), horizontal lines indicate medians, and whiskers extend to the most extreme values within 1.5 x IQR of the hinges.

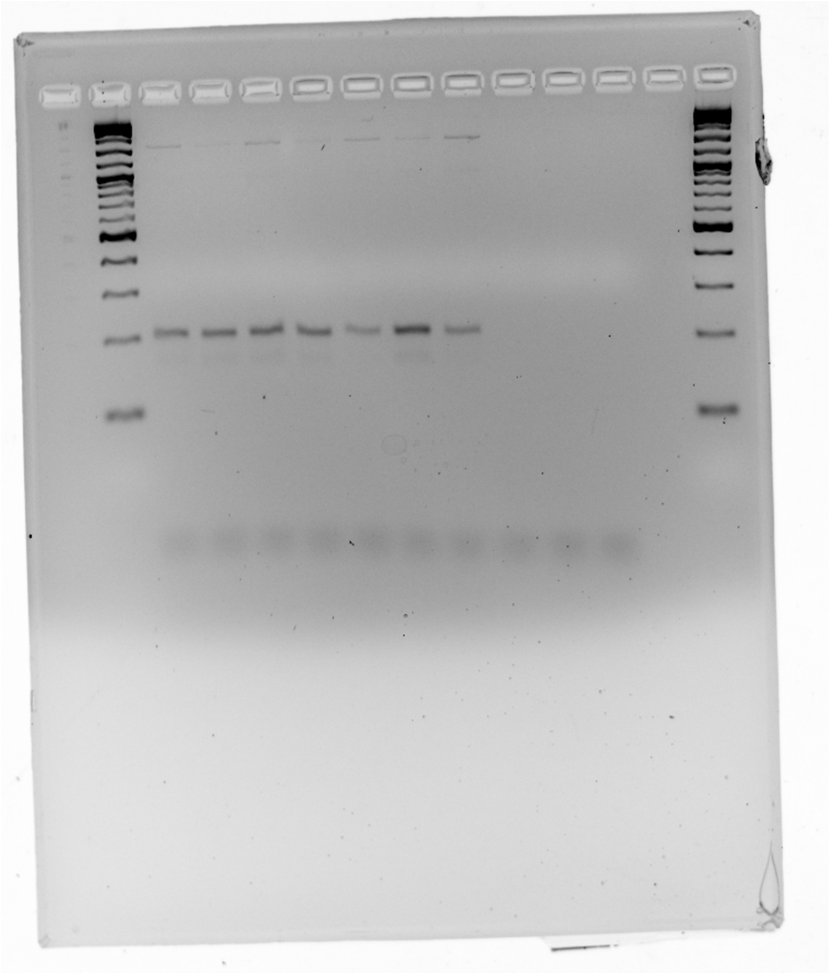

**Fig. S17.** Unprocessed original image of the gel presented in Fig. S9.
