## Supplementary Note 1 for "Age-related microbiome metabolites alter RNA splicing and chromatin accessibility in the brain"

This file contains archived images of the HMDB “Concentrations” entries for each metabolite included in the *in vitro* transcriptomic screen. The images document the concentration values reported for “Normal” blood samples in November 2023, when concentrations for the screen were selected (see Methods). Only values from adult samples were considered during concentration selection.

Alpha-ketoglutarate

| Biospecimen | Status | Value | Age | Sex | Condition |
|---|---|---|---|---|---|
| Blood | Detected but not Quantified | Not Quantified | Adult (>18 years old) | Both | Normal |
| Blood | Detected and Quantified | 7.0 (0.0-23.0) uM | Adult (>18 years old) | Both | Normal |
| Blood | Detected and Quantified | 8.6 +/- 2.6 uM | Children (1-13 years old) | Both | Normal |
| Blood | Detected and Quantified | 9.3 +/- 2.3 uM | Children (1-13 years old) | Both | Normal |
| Blood | Detected and Quantified | 8.9 +/- 2.7 uM | Adult (>18 years old) | Both | Normal |
| Blood | Detected but not Quantified | Not Quantified | Adult (>18 years old) | Both | Normal |
| Blood | Detected and Quantified | 4-13 uM | Not Specified | Not Specified | Normal |
| Blood | Detected and Quantified | 20.7 +/- 2.5 uM | Newborn (0-30 days old) | Both | Normal |
| Blood | Detected but not Quantified | Not Quantified | Adult (>18 years old) | Both | Normal |

Quinolinate

| Biospecimen | Status | Value | Age | Sex | Condition |
|---|---|---|---|---|---|
| Blood | Detected but not Quantified | Not Quantified | Adult (>18 years old) | Both | Normal |
| Blood | Detected and Quantified | 0.599 +/- 0.299 uM | Adult (>18 years old) | Both | Normal |
| Blood | Detected and Quantified | 0.47 +/- 0.047 uM | Adult (>18 years old) | Both | Normal |

#### 3-hydroxybutyrate (BHBA)

| Biospecimen | Status | Value | Age | Sex | Condition |
|---|---|---|---|---|---|
| Blood | Detected and Quantified | 60.0 +/- 20.0 uM | Adult (>18 years old) | Both | Normal |
| Blood | Detected and Quantified | 40.0 +/- 10.0 uM | Adult (>18 years old) | Both | Normal |
| Blood | Detected and Quantified | 323.0 +/- 29.0 uM | Adult (>18 years old) | Female | Normal |
| Blood | Detected and Quantified | 36 +/- 20 uM | Adult (>18 years old) | Both | Normal |
| Blood | Detected and Quantified | 86 +/- 53 uM | Adult (>18 years old) | Both | Normal |
| Blood | Detected and Quantified | 38 +/- 31 uM | Adult (>18 years old) | Both | Normal |
| Blood | Detected and Quantified | 50.0 (0.0-100.0) uM | Adult (>18 years old) | Both | Normal |
| Blood | Detected and Quantified | 46.5 +/- 10.1 uM | Adult (>18 years old) | Both | Normal |
| Blood | Detected and Quantified | 100.0 +/- 90.0 uM | Adult (>18 years old) | Female | Normal |
| Blood | Detected and Quantified | 36.0 (13.0-95.0) uM | Adult (>18 years old) | Both | Normal |
| Blood | Detected and Quantified | 80-3600 uM | Not Specified | Not Specified | Normal |
| Blood | Detected and Quantified | 64.88(60.4) uM | Adult (>18 years old) | Both | Normal |
| Blood | Detected and Quantified | 10.60-143.2 uM | Adult (>18 years old) | Both | Normal |
| Blood | Detected and Quantified | 76.9 +/- 66.3 uM | Adult (>18 years old) | Both | Normal |
| Blood | Detected and Quantified | 180.0 (22.0-700.0) uM | Adult (>18 years old) | Both | Normal |

#### Allantoin

| Biospecimen | Status | Value | Age | Sex | Condition |
|---|---|---|---|---|---|
| Blood | Detected and Quantified | 2.1 +/- 1.1 uM | Adult (>18 years old) | Both | Normal |
| Blood | Detected but not Quantified | Not Quantified | Adult (>18 years old) | Both | Normal |
| Blood | Detected but not Quantified | Not Quantified | Adult (>18 years old) | Both | Normal |

#### Cholate

| | | | | | |
|---|---|---|---|---|---|
| Blood | Detected and Quantified | 1.7 +/- 0.3 uM | Newborn (0-30 days old) | Both | Normal |
| Blood | Detected and Quantified | 0.1(0-0.1) uM | Infant (0-1 year old) | Both | Normal |
| Blood | Detected but not Quantified | Not Quantified | Adult (>18 years old) | Both | Normal |
| Blood | Detected and Quantified | 0.72 +/- 0.24 uM | Adult (>18 years old) | Both | Normal |

#### Choline

| Biospecimen | Status | Value | Age | Sex | Condition |
|---|---|---|---|---|---|
| Blood | Detected and Quantified | 6.0 +/- 0.3 uM | Adult (>18 years old) | Male | Normal |
| Blood | Detected but not Quantified | Not Quantified | Adult (>18 years old) | Both | Normal |
| Blood | Detected and Quantified | 27.5 +/- 3.3 uM | Newborn (0-30 days old) | Both | Normal |
| Blood | Detected and Quantified | 10.6 +/- 1.9 uM | Adult (>18 years old) | Both | Normal |
| Blood | Detected and Quantified | 6.8 +/- 0.3 uM | Adult (>18 years old) | Female | Normal |
| Blood | Detected and Quantified | 8.70-19.8 uM | Adult (>18 years old) | Both | Normal |
| Blood | Detected and Quantified | 14.5 +/- 5.3 uM | Adult (>18 years old) | Both | Normal |

### Glycine

| | | | | | |
|---|---|---|---|---|---|
| Blood | Detected and Quantified | 242.0 +/- 44.0 uM | Adult (>18 years old) | Male | Normal |
| Blood | Detected and Quantified | 258.0 +/- 64.0 uM | Adult (>18 years old) | Female | Normal |
| Blood | Detected and Quantified | 106-272 uM | Newborn (0-30 days old) | Both | Normal |
| Blood | Detected and Quantified | 150-350 uM | Infant (1 - 3 months old) | Both | Normal |
| Blood | Detected and Quantified | 125-400 uM | Children (3 months - 6 years old) | Both | Normal |
| Blood | Detected and Quantified | 140-490 uM | Children (6 - 18 years old) | Both | Normal |
| Blood | Detected and Quantified | 329.9 +/- 105.6 uM | Adult (>18 years old) | Both | Normal |
| Blood | Detected and Quantified | 147-321 uM | Not Specified | Not Specified | Normal |
| Blood | Detected but not Quantified | Not Quantified | Adult (>18 years old) | Both | Normal |
| Blood | Detected and Quantified | 149-301 uM | Infant (0-1 year old) | Not Specified | Normal |
| Blood | Detected and Quantified | 184-356 uM | Newborn (0-30 days old) | Not Specified | Normal |
| Blood | Detected and Quantified | 140-420 uM | Infant (0-1 year old) | Not Specified | Normal |
| Blood | Detected and Quantified | 200-600 uM | Newborn (0-30 days old) | Not Specified | Normal |
| Blood | Detected and Quantified | 80-341 uM | Infant (0-1 year old) | Not Specified | Normal |
| Blood | Detected but not Quantified | Not Quantified | Adult (>18 years old) | Both | Normal |
| Blood | Detected and Quantified | 100-320 uM | Newborn (0-30 days old) | Not Specified | Normal |
| Blood | Detected and Quantified | 126-384 uM | Newborn (0-30 days old) | Not Specified | Normal |
| Blood | Detected and Quantified | 280.0 (140.0-420.0) uM | Newborn (0-30 days old) | Both | Normal |
| Blood | Detected and Quantified | <300 uM | Children (1 - 13 years old) | Male | Normal |
| Blood | Detected and Quantified | 100 - 310 uM | Children (1 - 13 years old) | Not Specified | Normal |
| Blood | Detected and Quantified | 120-320 uM | Adult (>18 years old) | Both | Normal |
| Blood | Detected and Quantified | 213 +/- 35 uM | Infant (0-1 year old) | Both | Normal |
| Blood | Detected and Quantified | 56-308 uM | Infant (0-1 year old) | Both | Normal |
| Blood | Detected and Quantified | 60-310 uM | Children (1-13 years old) | Both | Normal |
| Blood | Detected and Quantified | 180-709 uM | Newborn (0-30 days old) | Not Specified | Normal |
| Blood | Detected and Quantified | 212.4 +/- 57.4 uM | Adult (>18 years old) | Male | Normal |
| Blood | Detected and Quantified | 150.0-440.0 uM | Adult (>18 years old) | Both | Normal |
| Blood | Detected and Quantified | 220 +/- 33 uM | Children (1 - 13 years old) | Both | Normal |
| Blood | Detected and Quantified | 300 +/- 114 uM | Adult (>18 years old) | Female | Normal |
| Blood | Detected and Quantified | 255.4 +/- 65.9 uM | Adult (>18 years old) | Both | Normal |
| Blood | Detected and Quantified | 306.63 uM | Children (1-13 years old) | Not Specified | Normal |
| Blood | Detected and Quantified | 236 +/- 43 uM | Adult (>18 years old) | Male | Normal |
| Blood | Detected and Quantified | 304.4(255.1-362.8) uM | Children (1-13 uears old) | Both | Normal |
| Blood | Detected and Quantified | 230.0 (178.0-282.0) uM | Adult (>18 years old) | Both | Normal |
| Blood | Detected and Quantified | 325.4 +/- 126.8 uM | Adult (>18 years old) | Both | Normal |
| Blood | Detected but not Quantified | Not Quantified | Adult (>18 years old) | Both | Normal |
| Blood | Detected and Quantified | 460.0 +/- 275.0 uM | Newborn (0-30 days old) | Not Specified | Normal |
| Blood | Detected and Quantified | 234.0 +/- 34.0 uM | Children (1-13 years old) | Male | Normal |

Glycodeoxycholate
N/A, see Methods

Hippurate

| Biospecimen | Status | Value | Age | Sex | Condition |
|---|---|---|---|---|---|
| Blood | Detected and Quantified | 3.0 (0.0-5.0) uM | Adult (>18 years old) | Both | Normal |
| Blood | Detected but not Quantified | Not Quantified | Adult (>18 years old) | Both | Normal |
| Blood | Detected and Quantified | 16.74 +/- 11.16 uM | Adult (>18 years old) | Both | Normal |
| Blood | Detected and Quantified | 1.000-28.00 uM | Adult (>18 years old) | Both | Normal |
| Blood | Detected and Quantified | <27.933 uM | Adult (>18 years old) | Both | Normal |
| Blood | Detected but not Quantified | Not Quantified | Adult (>18 years old) | Both | Normal |

Hypoxanthine

| | | | | | |
|---|---|---|---|---|---|
| Blood | Detected and Quantified | 2.300 +/- 1.100 uM | Adult (>18 years old) | Not Specified | Normal |
| Blood | Detected but not Quantified | Not Quantified | Adult (>18 years old) | Both | Normal |
| Blood | Detected and Quantified | 11.02 +/- 3.67 uM | Adult (>18 years old) | Both | Normal |
| Blood | Detected and Quantified | 1.7 +/- 0.4 uM | Adult (>18 years old) | Male | normal |
| Blood | Detected and Quantified | 2.2 +/- 1.1 uM | Adult (>18 years old) | Male | Normal |
| Blood | Detected and Quantified | 11.0294 +/- 3.676 uM | Adult (>18 years old) | Both | Normal |
| Blood | Detected and Quantified | 5.6 (3.1-7.1) uM | Adult (>18 years old) | Female | Normal |
| Blood | Detected and Quantified | 1.0 +/-0.9 uM | Adult (>18 years old) | Male | Normal |
| Blood | Detected and Quantified | 4.87 +/- 0.36 uM | Adult (>18 years old) | Both | Normal |
| Blood | Detected and Quantified | 1.0 +/- 0.9 uM | Adult (>18 years old) | Male | Normal |
| Blood | Detected and Quantified | 5.6 (3.1-7.1) uM | Adult (>18 years old) | Female | Normal |
| Blood | Detected and Quantified | 0.38 +/- 0.18 uM | Adult (>18 years old) | Male | Normal |
| Blood | Detected and Quantified | 34.2 +/- 10.3 uM | Adult (>18 years old) | Both | Normal |

Imidazole propionate
NA, see Methods

Indoleacetate, Indole-3-acetic acid

| Biospecimen | Status | Value | Age | Sex | Condition |
|---|---|---|---|---|---|
| Blood | Detected but not Quantified | Not Quantified | Adult (>18 years old) | Both | Normal |
| Blood | Detected and Quantified | 2.85 +/- 1.71 uM | Adult (>18 years old) | Both | Normal |
| Blood | Detected and Quantified | 0.100 +/- 0.100 uM | Adult (>18 years old) | Both | Normal |
| Blood | Detected and Quantified | 0.05 (0.0-0.118) uM | Adult (>18 years old) | Both | Normal |
| Blood | Detected and Quantified | 0.05 (0.0- 0.1) uM | Adult (>18 years old) | Both | Normal |

Indolelactate

| Biospecimen | Status | Value | Age | Sex | Condition |
|---|---|---|---|---|---|
| Blood | Detected and Quantified | 2.8 (0.5-5.0) uM | Adult (>18 years old) | Both | Normal |
| Blood | Detected but not Quantified | Not Quantified | Adult (>18 years old) | Both | Normal |

glycerate, glyceric acid

| Biospecimen | Status | Value | Age | Sex | Condition |
|---|---|---|---|---|---|
| Blood | Detected and Quantified | 10.0 (0.0-24.0) uM | Adult (>18 years old) | Both | Normal |
| Blood | Detected but not Quantified | Not Quantified | Adult (>18 years old) | Both | Normal |
| Blood | Detected and Quantified | <5 uM | Adult (>18 years old) | Both | Normal |
| Blood | Detected but not Quantified | Not Quantified | Adult (>18 years old) | Both | Normal |
| Blood | Detected but not Quantified | Not Quantified | Adult (>18 years old) | Both | Normal |
| Blood | Detected but not Quantified | Not Quantified | Adult (>18 years old) | Both | Normal |
| Blood | Detected but not Quantified | Not Quantified | Adult (>18 years old) | Female | Normal |
| Blood | Detected but not Quantified | Not Quantified | Adult (>18 years old) | Female | Normal |

Malate

| Biospecimen | Status | Value | Age | Sex | Condition |
|---|---|---|---|---|---|
| Blood | Detected and Quantified | 12.0 (0.0-21.0) uM | Adult (>18 years old) | Both | Normal |
| Blood | Detected but not Quantified | Not Quantified | Adult (>18 years old) | Both | Normal |
| Blood | Detected and Quantified | 3.2 +/- 0.9 uM | Adult (>18 years old) | Both | Normal |
| Blood | Detected but not Quantified | Not Quantified | Adult (>18 years old) | Both | Normal |

Carnitine

| Biospecimen | Status | Value | Age | Sex | Condition |
|---|---|---|---|---|---|
| Blood | Detected and Quantified | 29.74 +/- 7.55 uM | Adult (>18 years old) | Both | Normal |
| Blood | Detected and Quantified | 25.4-54.1 uM | Adult (>18 years old) | Female | Normal |
| Blood | Detected but not Quantified | Not Quantified | Adult (>18 years old) | Both | Normal |
| Blood | Detected and Quantified | 38.2 +/- 5.4 uM | Adult (>18 years old) | Female | Normal |
| Blood | Detected and Quantified | 43.0 (26.0-79.0) uM | Adult (>18 years old) | Both | Normal |
| Blood | Detected and Quantified | 43.57(9.97) uM | Adult (>18 years old) | Both | Normal |
| Blood | Detected and Quantified | 39.33(13.25) uM | Adult (>18 years old) | Both | Normal |
| Blood | Detected and Quantified | 19.0-65.0 uM | Adult (>18 years old) | Both | Normal |
| Blood | Detected and Quantified | 35.3 +/- 7.0 uM | Adult (>18 years old) | Both | Normal |
| Blood | Detected and Quantified | 45.7 +/- 11.6 uM | Adult (>18 years old) | Both | Normal |

Citrulline

| | | | | | |
|---|---|---|---|---|---|
| Blood | Detected and Quantified | 38.0 (30.0-46.0) uM | Adult (>18 years old) | Both | Normal |
| Blood | Detected and Quantified | <43 uM | Adult (>18 years old) | Both | Normal |
| Blood | Detected and Quantified | 29.8 +/- 7.91 uM | Adult (>18 years old) | Both | Normal |
| Blood | Detected and Quantified | 35 +/- 10 uM | Adult (>18 years old) | Female | Normal |
| Blood | Detected and Quantified | 37.0 +/- 9.0 uM | Adult (>18 years old) | Male | Normal |
| Blood | Detected and Quantified | 35.0 +/- 10.0 uM | Adult (>18 years old) | Female | Normal |
| Blood | Detected and Quantified | 37 +/- 9 uM | Adult (>18 years old) | Male | Normal |
| Blood | Detected and Quantified | 38.0 (30.0-46.0) uM | Adult (>18 years old) | Not Specified | Normal |

Isoleucine

| | | | | | |
|---|---|---|---|---|---|
| Blood | Detected and Quantified | 84.0 +/- 18.0 uM | Adult (>18 years old) | Male | Normal |
| Blood | Detected and Quantified | 81.0 +/- 18.0 uM | Adult (>18 years old) | Male | Normal |
| Blood | Detected and Quantified | 56.0 +/- 12.0 uM | Adult (>18 years old) | Female | Normal |
| Blood | Detected and Quantified | 68 +/- 11 uM | Adult (>18 years old) | Male | Normal |
| Blood | Detected and Quantified | 30-108 uM | Adult (>18 years old) | Not Specified | Normal |
| Blood | Detected and Quantified | 53 +/- 7 uM | Adult (>18 years old) | Female | Normal |
| Blood | Detected and Quantified | 64 +/- 13 uM | Adult (>18 years old) | Female | Normal |
| Blood | Detected and Quantified | 40.00-100.0 uM | Adult (>18 years old) | Both | Normal |
| Blood | Detected and Quantified | 77.5 +/- 15.4 uM | Adult (>18 years old) | Both | Normal |
| Blood | Detected and Quantified | 84 +/- 18 uM | Adult (>18 years old) | Male | Normal |
| Blood | Detected and Quantified | 62.0 (48.0-76.0) uM | Adult (>18 years old) | Both | Normal |
| Blood | Detected and Quantified | 60.7 +/- 18.6 uM | Adult (>18 years old) | Both | Normal |

Lysine

| | | | | | |
|---|---|---|---|---|---|
| Blood | Detected and Quantified | 198.0 +/- 31.0 uM | Adult (>18 years old) | Male | Normal |
| Blood | Detected and Quantified | 183.0 +/- 34.0 uM | Adult (>18 years old) | Female | Normal |
| Blood | Detected and Quantified | 71-151 uM | Adult (>18 years old) | Not Specified | Normal |
| Blood | Detected and Quantified | 189 +/- 27 uM | Adult (>18 years old) | Male | Normal |
| Blood | Detected and Quantified | 161 +/- 21 uM | Adult (>18 years old) | Female | Normal |
| Blood | Detected and Quantified | 105.100 +/- 30.100 uM | Not Specified | Not Specified | Normal |
| Blood | Detected and Quantified | 434.0 +/- 23.0 uM | Adult (>18 years old) | Both | Normal |
| Blood | Detected and Quantified | 110.0-240.0 uM | Adult (>18 years old) | Both | Normal |
| Blood | Detected and Quantified | 197.4 +/- 30.7 uM | Adult (>18 years old) | Both | Normal |
| Blood | Detected but not Quantified | Not Quantified | Adult (>18 years old) | Both | Normal |
| Blood | Detected and Quantified | 197 (184-216) uM | Adult (>18 years old) | Both | Normal |
| Blood | Detected and Quantified | 178.6 +/- 58.2 uM | Adult (>18 years old) | Both | Normal |
| Blood | Detected and Quantified | 188.0 (156.0-220.0) uM | Adult (>18 years old) | Both | Normal |

Valine

| Biospecimen | Status | Value | Age | Sex | Condition |
|---|---|---|---|---|---|
| Blood | Detected and Quantified | 212.3 +/- 61.3 uM | Adult (>18 years old) | Both | Normal |
| Blood | Detected but not Quantified | Not Quantified | Adult (>18 years old) | Both | Normal |
| Blood | Detected and Quantified | 266.3 +/- 61 uM | Adult (>18 years old) | Both | Normal |
| Blood | Detected and Quantified | 233.0 (190.0-276.0) uM | Adult (>18 years old) | Both | Normal |
| Blood | Detected and Quantified | 252.0 +/- 37.0 uM | Adult (>18 years old) | Male | Normal |
| Blood | Detected and Quantified | 209.0 +/- 31.0 uM | Adult (>18 years old) | Female | Normal |
| Blood | Detected and Quantified | 119-336 uM | Adult (>18 years old) | Not Specified | Normal |
| Blood | Detected and Quantified | 257 +/- 39 uM | Adult (>18 years old) | Male | Normal |
| Blood | Detected and Quantified | 210 +/- 19 uM | Adult (>18 years old) | Female | Normal |
| Blood | Detected and Quantified | 209 +/- 31 uM | Adult (>18 years old) | Female | Normal |
| Blood | Detected and Quantified | 243.6 +/- 45.2 uM | Adult (>18 years old) | Both | Normal |
| Blood | Detected and Quantified | 252 +/- 37 uM | Adult (>18 years old) | Male | Normal |
| Blood | Detected and Quantified | 140.0-300.0 uM | Adult (>18 years old) | Both | Normal |

methyl indole-3-acetate
N/A, see Methods

N-acetylalanine
N/A, see Methods

Phenylacetylglutamine

| Biospecimen | Status | Value | Age | Sex | Condition |
|---|---|---|---|---|---|
| Blood | Detected but not Quantified | Not Quantified | Adult (>18 years old) | Both | Normal |
| Blood | Detected but not Quantified | Not Quantified | Adult (>18 years old) | Both | Normal |
| Blood | Detected and Quantified | 3.34 +/- 0.31 uM | Adult (>18 years old) | Both | Normal |
| Blood | Detected and Quantified | <17.803 uM | Adult (>18 years old) | Both | Normal |

Nicotinamide

| Biospecimen | Status | Value | Age | Sex | Condition |
|---|---|---|---|---|---|
| Blood | Detected but not Quantified | Not Quantified | Adult (>18 years old) | Both | Normal |
| Blood | Detected and Quantified | 0.03 +/- 0.01 uM | Adult (>18 years old) | Both | Normal |
| Blood | Detected and Quantified | 0.44 +/- 0.0054 uM | Adult (>18 years old) | Both | Normal |

oxalate (ethanedioate)

| Biospecimen | Status | Value | Age | Sex | Condition |
|---|---|---|---|---|---|
| Blood | Detected and Quantified | 6.43 +/- 1.06 uM | Adult (>18 years old) | Both | Normal |
| Blood | Detected and Quantified | 22.2 uM | Adult (>18 years old) | Both | Normal |
| Blood | Detected and Quantified | 9.2 +/- 2.7 uM | Adult (>18 years old) | Male | Normal |
| Blood | Detected and Quantified | 3.33 +/- 1.11 uM | Adult (>18 years old) | Both | Normal |
| Blood | Detected and Quantified | 3.333 +/- 1.111 uM | Adult (>18 years old) | Both | Normal |

p-cresol sulfate

| Biospecimen | Status | Value | Age | Sex | Condition |
|---|---|---|---|---|---|
| Blood | Detected but not Quantified | Not Quantified | Adult (>18 years old) | Both | Normal |
| Blood | Detected and Quantified | 27.099 +/- 13.28 uM | Adult (>18 years old) | Both | Normal |

palmitate (16:0)C

| Biospecimen | Status | Value | Age | Sex | Condition |
|---|---|---|---|---|---|
| Blood | Detected and Quantified | 122 +/- 48 uM | Adult (>18 years old) | Both | Normal |
| Blood | Detected but not Quantified | Not Quantified | Adult (>18 years old) | Both | Normal |
| Blood | Detected and Quantified | 66.010 +/- 9.879 uM | Adult (>18 years old) | Both | Normal |
| Blood | Detected and Quantified | 1060.743 +/- 35.0981 uM | Adult (>18 years old) | Male | Normal |
| Blood | Detected and Quantified | 1064.643 +/- 50.697 uM | Adult (>18 years old) | Male | Normal |
| Blood | Detected and Quantified | 63.8 +/- 0.400 uM | Adult (>18 years old) | Both | Normal |
| Blood | Detected and Quantified | 30.49 +/- 2.64 uM | Adult (>18 years old) | Both | Normal |
| Blood | Detected and Quantified | 22.7-32.9 uM | Adult (>18 years old) | Female | Normal |
| Blood | Detected and Quantified | 366 uM | Adult (>18 years old) | Both | Normal |
| Blood | Detected and Quantified | 424 uM | Adult (>18 years old) | Both | Normal |
| Blood | Detected and Quantified | 1415.468 +/- 323.410 uM | Adult (>18 years old) | Both | Normal |
| Blood | Detected and Quantified | 1215.798 +/- 212.0316 uM | Adult (>18 years old) | Female | Normal |
| Blood | Detected and Quantified | 2500 +/- 630 uM | Adult (>18 years old) | Female | Normal |
| Blood | Detected and Quantified | 2360 +/- 430 uM | Adult (>18 years old) | Male | Normal |

*Continued on next page →*

| | | | | | |
|---|---|---|---|---|---|
| Blood | Detected and Quantified | 1052.943 +/- 54.597 uM | Adult (>18 years old) | Male | Normal |
| Blood | Detected and Quantified | 26.7 +/- 4.4 uM | Adult (>18 years old) | Both | Normal |
| Blood | Detected and Quantified | 1029.544 +/- 62.397 uM | Adult (>18 years old) | Male | Normal |
| Blood | Detected and Quantified | 52 uM | Adult (>18 years old) | Both | Normal |
| Blood | Detected and Quantified | 53 uM | Adult (>18 years old) | Both | Normal |
| Blood | Detected and Quantified | 74 uM | Adult (>18 years old) | Both | Normal |
| Blood | Detected and Quantified | 80 uM | Adult (>18 years old) | Both | Normal |
| Blood | Detected and Quantified | 95 uM | Adult (>18 years old) | Both | Normal |
| Blood | Detected and Quantified | 114 uM | Adult (>18 years old) | Both | Normal |
| Blood | Detected and Quantified | 129 uM | Adult (>18 years old) | Both | Normal |
| Blood | Detected and Quantified | 131 uM | Adult (>18 years old) | Both | Normal |
| Blood | Detected and Quantified | 147 uM | Adult (>18 years old) | Both | Normal |
| Blood | Detected and Quantified | 22.3-31.1 uM | Adult (>18 years old) | Male | Normal |
| Blood | Detected and Quantified | 347 uM | Adult (>18 years old) | Both | Normal |

Pantothenate

| Biospecimen | Status | Value | Age | Sex | Condition |
|---|---|---|---|---|---|
| Blood | Detected but not Quantified | Not Quantified | Adult (>18 years old) | Both | Normal |
| Blood | Detected and Quantified | 2.86 +/- 0.94 uM | Adult (>18 years old) | Female | Normal |
| Blood | Detected and Quantified | 4.91 +/- 0.38 uM | Adult (>18 years old) | Both | Normal |
| Blood | Detected but not Quantified | Not Quantified | Adult (>18 years old) | Both | Normal |
| Blood | Detected and Quantified | 0.30-1.80 uM | Adult (>18 years old) | Both | Normal |

trimethylamine N-oxide

| Biospecimen | Status | Value | Age | Sex | Condition |
|---|---|---|---|---|---|
| Blood | Detected but not Quantified | Not Quantified | Adult (>18 years old) | Both | Normal |
| Blood | Detected and Quantified | 1.81 +/- 1.60 uM | Adult (>18 years old) | Both | Normal |
| Blood | Detected and Quantified | 38.81 +/- 20.37 uM | Adult (>18 years old) | Both | Normal |
| Blood | Detected but not Quantified | Not Quantified | Adult (>18 years old) | Both | Normal |
| Blood | Detected and Quantified | 37.8 +/- 20.4 uM | Adult (>18 years old) | Both | Normal |

urate

| | | | | | |
|---|---|---|---|---|---|
| Blood | Detected and Quantified | 279.578 +/- 23.794 uM | Adult (>18 years old) | Male | normal |
| Blood | Detected and Quantified | 303 +/- 59 uM | Adult (>18 years old) | Male | Normal |
| Blood | Detected and Quantified | 150-360 uM | Adult (>18 years old) | Female | Normal |
| Blood | Detected and Quantified | 140-410 uM | Adult (>18 years old) | Female | Normal |
| Blood | Detected and Quantified | 377.6 +/- 82.6 uM | Adult (>18 years old) | Male | Normal |
| Blood | Detected and Quantified | 494.2 uM | Adult (>18 years old) | Both | Normal |
| Blood | Detected and Quantified | 372.0 (238.0-506.0) uM | Adult (>18 years old) | Male | Normal |
| Blood | Detected and Quantified | 298.0 (149.0-446.0) uM | Adult (>18 years old) | Female | Normal |
| Blood | Detected and Quantified | 271.95 +/- 43.13 uM | Adult (>18 years old) | Both | Normal |
| Blood | Detected and Quantified | 256.8 +/- 7.0 uM | Adult (>18 years old) | Both | Normal |
| Blood | Detected and Quantified | 291.475 +/- 59.485 uM | Adult (>18 years old) | Not Specified | Normal |
| Blood | Detected and Quantified | 240.91 +/- 82.68 uM | Adult (>18 years old) | Both | Normal |
| Blood | Detected and Quantified | 302.0 +/- 60.0 uM | Adult (>18 years old) | Male | Normal |
| Blood | Detected and Quantified | 234.0 +/- 52.0 uM | Adult (>18 years old) | Female | Normal |
| Blood | Detected and Quantified | 200-420 uM | Adult (>18 years old) | Both | Normal |
